# Population coding in the cerebellum and its implications for learning from error

**DOI:** 10.1101/2020.05.18.102376

**Authors:** Reza Shadmehr

## Abstract

The cerebellum resembles a feedforward, three-layer network of neurons in which the “hidden layer” consists of Purkinje cells (P-cells), and the output layer consists of deep cerebellar nucleus (DCN) neurons. However, unlike an artificial network, P-cells are grouped into small populations that converge onto single DCN neurons. Why are the P-cells organized in this way, and what is the membership criterion of each population? To consider these questions, in this review I apply elementary mathematics from machine learning and assume that the output of each DCN neuron is a prediction that is compared to the actual observation, resulting in an error signal that originates in the inferior olive. This signal is sent to P-cells via climbing fibers that produce complex spikes. The same error signal from the olive must also guide learning in the DCN neurons, yet the olivary projections to the DCN are weak, particularly in adulthood. However, P-cells that form a population exhibit a special property: they can synchronize their complex spikes, which in turn suppresses activity of the DCN neuron that produced the erroneous output. Viewed in the framework of machine learning, it appears that the olive organizes the P-cells into populations so that through complex spike synchrony each population can act as a surrogate teacher for the DCN neuron it projects to. This error-dependent grouping of P-cells into populations gives rise to a number of remarkable features of behavior, including multiple timescales of learning, protection from erasure, and spontaneous recovery of memory.

---

*Ever tried. Ever failed. No Matter. Try again. Fail again. Fail better*.

Samuel Beckett

## Introduction

During electrophysiological recording from Purkinje cells (P-cells), listening to the sound of spikes as they are reported by an electrode, one cannot help but be impressed by the complex spike. Whereas the simple spikes appear ordinary and common, rain drops falling on the roof, the complex spike is more like lightening, a thunderous event that makes the P-cell pause its production of simple spikes. Indeed, after generating a complex spike, a P-cell requires 10-20 ms of recovery before it resumes production of simple spikes (Thach, 1967).

The significance of the complex spike as an event in the life of a P-cell is illustrated by the fact that persistent stimulation of climbing fibers, the sole source of complex spikes, can lead to excitotoxic damage of P-cells (Slemmer et al., 2005). Indeed, sustained increase or decrease in the rate of complex spikes above or below baseline damages the P-cells (O’Hearn and Molliver, 1997). Thus, survival of a P-cell depends on its ability to tame the production of complex spikes, regulating it to around baseline (De Schutter, 1995; Mauk and Donegan, 1997).

Complex spikes arise from climbing fibers that originate from cells in the inferior olive (Eccles et al., 1966). The olive cells receive inhibitory projections from the neurons that reside in the deep cerebellar nucleus (DCN) (de Zeeuw et al., 1988), the cells that carry the output of the cerebellum (Kim et al., 1998; Rasmussen et al., 2015; Rasmussen and Hesslow, 2014). The olive cells receive excitatory inputs from other regions of the brain (Kralj-Hans et al., 2007). In analogy to a neural network, activity of DCN neurons are a form of prediction (Doya, 1999), allowing the olive cells to compare this prediction to the actual observations. Indeed, a complex spike is often generated after unexpected occurrence of a broad class of sensory inputs (Andersson and Armstrong, 1987; Ju et al., 2019), motor actions (Welsh et al., 1995), or rewarding events (Heffley et al., 2018).

Thus, as a first approximation the signal in the climbing fiber is the difference between what was reported to the olive (the observation), and what was outputted by the cerebellum (the prediction). Said in another way, neurons in the inferior olive are in position to compare the desired response that they have received, i.e., the excitatory input from a region of the brain, with the actual output of a DCN neuron, and produce an error signal that reflects the difference between the two (De Zeeuw et al., 1998; Kitazawa et al., 1998; Simpson et al., 1996; Soetedjo et al., 2008). From a machine learning perspective, this signal is termed a prediction error.

Prediction errors are fundamental to learning in artificial neural networks. In such networks, the rules of learning stipulate that the error associated with a given output neuron must be conveyed to that neuron as well as to all other neurons that project directly or indirectly to it. Surprisingly, the DCN neurons receive a rather weak signal from the olive (Lu et al., 2016), a signal that weakens further as the animal reaches adulthood (Najac and Raman, 2017). In comparison, the middle layer neurons (the P-cells) receive an extraordinarily strong signal from the olive. However, despite the fact that each P-cell projects to about 4 DCN neurons (Person and Raman, 2012b), a P-cell receives only a single climbing fiber.

The weak olivary input to the DCN is puzzling because it suggests that the error signal from the olive will have difficulty acting as a teacher for the output layer neurons. The single climbing fiber to a P-cell is also puzzling because it implies that the error signal is not a fair reflection of the entire error space, but rather biased to provide only a limited view.

To illustrate the implications of these puzzles, consider the following example. Suppose you are playing basketball and shoot from the free throw line and your ball misses the basket. If you are looking at the basket as the ball passes to one side, say to the right, the event will produce an increase in probability of complex spikes among P-cells that prefer rightward prediction errors in visual space (Herzfeld et al., 2015; Soetedjo et al., 2008; Soetedjo and Fuchs, 2006). That same error will reduce probability of complex spikes in P-cells that prefer leftward prediction errors. However, the rightward error will have no consequence for the complex spikes of P-cells that prefer upward or downward errors. As a result, each P-cell receives a limited view of the error space: some care about rightward errors and some care about leftward errors, but none care about all parts of the error space. In addition, because the olivary projections to the DCN are weak (Lu et al., 2016), learning from error faces a formidable obstacle: how can the DCN neurons that contributed to your movement learn about their erroneous output?

Thus, when we view the cerebellum from the perspective of machine learning, two puzzles emerge. It is surprising that the error information from the olive is conveyed strongly to the middle layer neurons (the P-cells), but not the output layer neurons (the DCN). Furthermore, it is surprising that the olivary signal to each P-cell consists of a single input that is biased toward a specific part of the error space.

We will suggest that the very limited distribution of output from a single olive cell to a handful of P-cells organizes the P-cells into populations that share the same teacher (De Zeeuw et al., 2011; Heck et al., 2013): P-cells that learn from the same teacher likely project together as a population to a single neuron in the DCN. Viewed from the perspective of machine learning, this organization of P-cells serves a critical function: it allows the olive to not only convey error information to a small group of P-cells, but through synchronous production of complex spikes, also convey the same error information to the DCN neuron that was responsible for the error (Chaumont et al., 2013; Tang et al., 2019). The result is a framework in which each olivary cell is the teacher to a handful of P-cells (Gao et al., 2012), and those P-cells are the teachers for the DCN neuron that produced the erroneous output (Medina and Mauk, 1999).

This error-dependent organization of P-cells into populations can produce a number of remarkable features. Because an error may be preferred by some P-cell populations, increasing their probability of complex spikes, while the same error will be anti-preferred for other populations, decreasing their probability of complex spikes, the differing sensitivities to error, as reflected in the differing rates of plasticity associated with the presence vs. absence of a complex spike (Herzfeld et al., 2018; Yang and Lisberger, 2014a), are likely to contribute to multiple timescales of adaptation, some fast, others slow.

In addition, the personalized view of error afforded to each P-cell may provide protection from erasure. That is, behavior becomes easier to learn than to unlearn. Rather than countering plasticity in the P-cells that learned from error, reversal of error will engage learning in a separate population of P-cells, possibly in a different olivocerebellar module. As a result, organizing P-cells into error-dependent populations may be responsible for a paradoxical aspect of behavior: learning followed by extinction returns behavior to baseline, but following passage of time, actions revert back toward that which was learned initially, thus producing spontaneous recovery of memory (Criscimagna-Hemminger and Shadmehr, 2008; Kojima et al., 2004; Sarwary et al., 2018; Smith et al., 2006).

Our theoretical analysis relies on the assumption that the olivary input to the cerebellum reflects the erroneous output of a DCN neuron. Although this assumption is consistent with anatomical (de Zeeuw et al., 1988) and physiological data (Soetedjo et al., 2008), it remains a speculation because there are no simultaneous recordings from olive projecting DCN neurons and the feedback that they receive from the olive. Indeed, the assumption that climbing fiber activity reflects a prediction error is not universally accepted (Horn et al., 2004), and faces challenges including the fact that the probability of complex spikes poorly encodes the magnitude of prediction error (Catz et al., 2005), and the fact that cerebellar dependent learning can take place without obvious modulation of complex spikes (Hewitt et al., 2015; Ke et al., 2009). In some cases, modulation of simple spikes alone is sufficient to produce a form of learning (Nguyen-Vu et al., 2013). Cognizant of these observations, we will develop the theoretical results based on the assumption that olivary input provides an error signal, and then consider the limitations of our assumption.

## Prediction errors of the cerebellum

The anatomy of the cerebellum resembles a feedforward network (Fig. 1A) (Raymond and Medina, 2018). Like an artificial neural network, inputs (mossy fibers) bring information to the first layer of neurons (granule cells), which distribute them to the second layer (P-cells), which in turn project to the output layer (DCN neurons) (Fig. 1B). To be sure, this resemblance is an approximation. The input from the granule cells to the P-cells is direct, as well as indirect (via molecular layer interneurons). P-cells send a few collateral axons to neighboring P-cells (Witter et al., 2016), and at least in some lobules, P-cells also send a few collaterals to granule cells (Guo et al., 2016). As a result, there is a degree of synchrony among neighboring P-cells (Sedaghat-Nejad et al., 2019), a synchrony that remains even when chemical synapses are inactivated (Han et al., 2018). Despite these complications, a feedforward network is a useful approximation, particularly from the P-cell layer to the output layer, which is the focus of our analysis.

**Figure 1.**
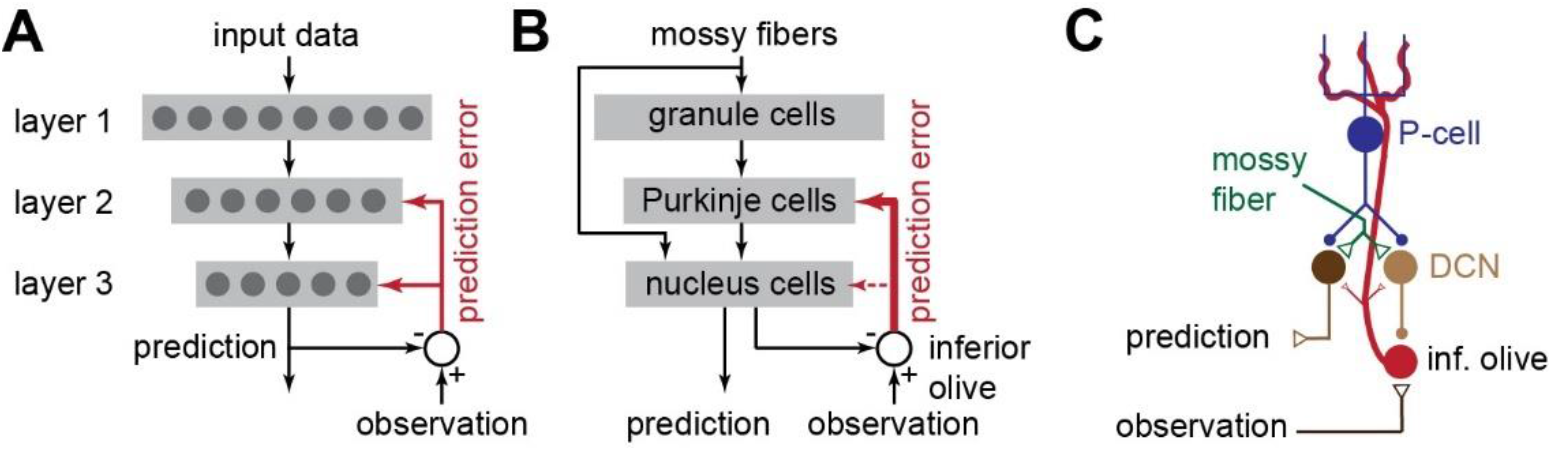
A feedforward network as a simplified model of the cerebellum. **A**. In an artificial neural network with three layers, the third layer provides the predictions, which are compared to observations and then fed back via an error signal to units in layers 2 and 3. **B**. In the cerebellum, the 3-layers are comprised of granule cells, Purkinje cells, and deep cerebellar nucleus neurons. The predictions of the cerebellum are conveyed via GABA-ergic DCN neurons to the inferior olive, which in turn provides the cerebellum with an error signal. This signal is conveyed strongly to the P-cells, but weakly to the DCN neurons (dashed line). The predictions of the cerebellum are also conveyed via non-GABA-ergic DCN neurons to the rest of the brain. **C**. A single P-cell projects to both GABA-ergic and non-GABA-ergic DCN neurons. However, the error signal sent from the olive to the cerebellum depends directly on the GABA-ergic DCN neurons, but not on the non-GABA-ergic neurons. Filled circles are inhibitory synapses, triangles are excitatory synapses.

In a typical artificial neural network, all cells in each layer connect to all cells in the next layer. However, in the cerebellum (of rodents), about 50 P-cells converge upon a single DCN neuron (Person and Raman, 2012a). That is, P-cells organize into small populations and with their simple spikes modulate the output of the DCN neurons. What determines the membership in this population? That is, what property must a P-cell have to make it eligible to be a member of the select group that projects to a given DCN neuron? To answer this question, let us consider this problem from the perspective of machine learning.

The activities of the DCN neurons represent the output of the cerebellum, but the nature of these outputs is diverse because the cerebellum projects to numerous sensory (May et al., 1990), motor (Noda et al., 1990), and reward related structures (Carta et al., 2019). We label the activation, i.e. firing rate, of each DCN neuron as 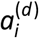, where the superscript *d* refers to DCN neurons. Some of the DCN neurons are GABA-ergic and inhibit the olive (Bazzigaluppi et al., 2012; Lefler et al., 2014). Activity of these DCN neurons is labeled as 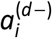. Another subset is non-GABA-ergic and send their axons to other regions, e.g., brainstem, superior colliculus, red nucleus, thalamus, etc. It is with these non-GABA-ergic neurons that the cerebellum influences behavior. We label activity of the non-olive projecting DCN neurons as 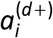. Thus, in the output layer the superscript *d* includes GABA-ergic as well as non-GABA-ergic projecting DCN neurons: 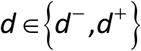.

We assume that a cell in the olive compares the output 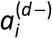 that it receives from a DCN neuron with the observed event 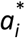. For example, suppose the output 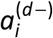 is predicting location of the visual information that should be present following conclusion of a saccadic eye movement. In this case, the actual location of the visual event 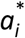 is conveyed from the superior colliculus to the olivary cell that receives the cerebellar output. If the cerebellar output does not match the actually observed collicular activity, then the olivary cell responds, producing spikes in its cerebellar projections (Kojima and Soetedjo, 2018, 2017). Because the collicular activity reflects not only the position of the visual stimulus, but also its reward value (Ikeda and Hikosaka, 2003), a difference in the predicted reward value of the stimulus as compared to the observed value will also be an error signal that is conveyed to the cerebellum (Heffley et al., 2018).

Thus, given output 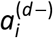 from the GABA-ergic DCN neurons, we assume that the activity in the inferior olive neuron that projects back to the DCN and the rest of the cerebellum is an error signal that conveys the difference between what was observed (excitatory input to the olive from a brain region), and the cerebellar output (inhibitory output from the DCN to the olive). Notably, this error is only associated with activity of GABA-ergic DCN neurons 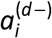. Indeed, De Zeeuw et al. (1997) were the first to note existence of projections from the olive back to the GABA-ergic neurons of the DCN. Let us label this error signal as 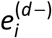:

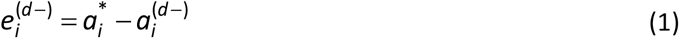

In the above equation, the error signal depends on the activity 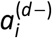 of GABA-ergic olive-projecting DCN neurons, but not directly on the activity 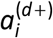 of the non-GABA-ergic, non-olive projecting DCN neurons. This produces our first puzzle:

**Puzzle P1.** *Despite the fact that non-GABA-ergic DCN neurons are responsible for conveying output of the cerebellum, the error associated with their output is not directly a part of the olivary signal back to the cerebellum. How do non-GABA-ergic DCN neurons receive information regarding the error in their activities?*

The consequence of computing error information based on the activity of GABA-ergic DCN neurons, and not the other output neurons, can be illustrated by an experiment by Kim et al. (1998). The authors presented rabbits a tone followed by an air puff, teaching them to blink just before arrival of the air puff. In naïve animals the air puff alone produced complex spikes in P-cells. Early in training with the tone+air puff trials the P-cells continued to produce complex spikes following the air puff, but late in training tone+air puff no longer produced complex spikes: as performance improved and errors were reduced, so were the complex spikes. Now the authors took the well-trained animals and injected a GABA antagonist into the inferior olive. This effectively eliminated the efficacy of the DCN input to the olive. However, blocking the DCN input re-introduced the complex spikes during tone+air puff trials. That is, although the animal continued to blink in response to the tone, the olive nevertheless signaled prediction errors to the cerebellum. This suggests that even when non-GABA-ergic DCN neurons are correctly predicting an output 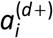, if the inhibitory input to the olive is suppressed, the error signal to the cerebellum returns.

To consider this puzzle, let us further develop the mathematics associated with training our network. A generic DCN neuron’s activity 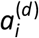 depends partly on the inputs that it receives, and partly on its internal state. The inputs are the weighted activities of neurons that project to it. This includes inputs from P-cells (via simple spikes that act on inhibitory synapses), and inputs from mossy fibers (via excitatory synapses), as shown in Fig. 1C. We approximate the inputs to a generic DCN neuron via the following equation:

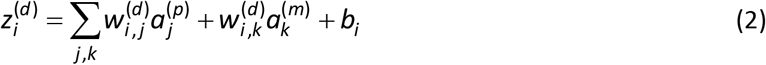

In the above equation, 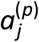 is the activity of P-cells that project to a given DCN neuron *i*, 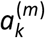 is the activity conveyed via mossy fibers to that neuron, and *b_i_* is the internal bias of that neuron (making it possible for the neuron to fire despite having inhibitory inputs).

Notice that in Eq. (2) we omitted the influence of projections to the DCN neuron from the olive. This is because the axons that bring olivary input to the cerebellum fire at rates that are two orders of magnitude smaller than P-cells and mossy fibers. Furthermore, in the adult animal (mice), the excitatory post-synaptic currents (EPSCs) that are produced in a typical DCN neuron following activation of olivary axons are remarkably small (Lu et al., 2016). For example, activation of olivary axons can produce an EPSC event with an amplitude of 2 pA in a P-cell, 0.4 pA in the DCN neuron of a juvenile mouse (Najac and Raman, 2017), and only 8×10^−3^ pA in the DCN neuron of an adult mouse (Lu et al., 2016). Therefore, in the adult animal the input to the DCN from the olive is weak. This fact simplifies our equation, but also introduces a second puzzle:

**Puzzle P2.** *Given that the olive’s input to the DCN is weak, how does a DCN neuron receive information about the error in its output?*

To summarize, an error signal is needed to teach the output layer of our network (the DCN neurons). We assume that this error signal is computed in the olive. However, projections from the olive to the DCN represent the error that was made by only a subset of output neurons: GABA-ergic olive projecting DCN neurons. Furthermore, the axons that project from the olive to the DCN have weak synapses in adulthood, and are essentially silent as compared to mossy fiber and P-cell axons that converge on the same neurons. How can the olive be an effective teacher for the DCN neurons?

## The problem of teaching a deep cerebellar nucleus neuron

The purpose of learning in a neural network is to minimize a loss function, typically the sum of squared errors. For our network, the sum of errors is a function of all output neurons, i.e., both olive-projecting and non-olive-projecting DCN neurons:

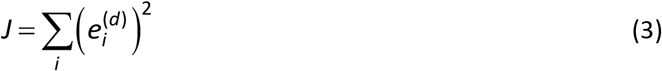

To minimize this loss, we need to change the activity of both the GABA-ergic DCN neurons 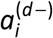, and the non-GABA-ergic DCN neurons 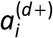. This is done by changing the weights associated with the inputs that these neurons receive, as well as their internal biases, i.e., their intrinsic excitability.

The activity (firing rate) of a generic neuron in our network is related to its inputs and internal biases via a non-linearity, for example a sigmoidal function:

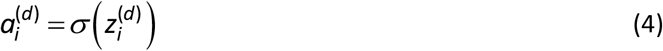

To change the synaptic weight of a given input to a neuron in the output layer, we compute the gradient of the loss function with respect to that weight. For example, the weight change for a GABA-ergic DCN neuron is negatively proportional to the following gradient:

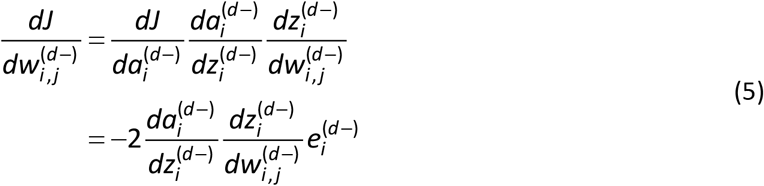

The above expression has a simple meaning: the gradient of a DCN neuron’s input *z_i_* with respect to its mossy fiber synaptic weights is positive (because those inputs are excitatory), and thus positive errors (e.g., olivary output above baseline) should increase the mossy fiber synaptic weights. In comparison, the gradient of *z_i_* with respect to P-cell synaptic weights is negative (because those inputs are inhibitory), and thus positive errors should decrease those synaptic weights. Similarly, to change the intrinsic excitability of the DCN neuron, we compute the gradient with respect to the bias:

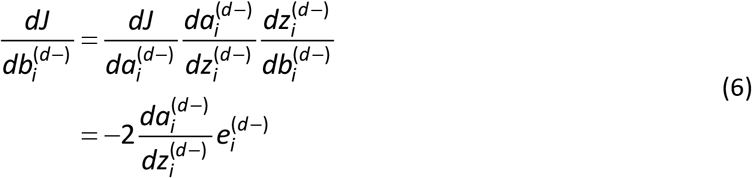

Aside from providing general rules for learning in the DCN, the above expressions state that the gradients of our loss function with respect to both weights and internal biases of a given DCN neuron are proportional to the prediction errors of that neuron. This makes an important anatomical prediction: if a DCN neuron projects to the olive, it must receive error information associated with its own output, not the output of some other DCN neuron. This leads to our first conjecture:

**Conjecture C1**. *A DCN neuron receives error feedback from precisely the same olive neurons it inhibits*.

Indeed, olivary projections to the DCN are reciprocal: if a region in the olive projects to a region in the DCN, that DCN region also projects back to that specific region of the olive (Ruigrok and Voogd, 2000). However, even if conjecture C1 were true, we still have DCN neurons that do not project to the olive, but rather project to some other region of the brain (puzzle P1). In principle, because the non-GABA-ergic neurons do not project to the olive, we have no error signal with which to teach them.

There are several ways to consider this problem. First, it is possible that as the non-GABA-ergic DNC neurons project to their destination, they synapse on GABA-ergic neurons, and those GABA-ergic neurons then project back to the olive. In this way, the olivary cell would indirectly receive the output of the non-GABA-ergic DCN neuron. However, this introduces a delay in communication, and importantly, imposes a timing difference in the comparison of outputs with observations for some DCN neurons (non-GABA-ergic) with respect to others (GABA-ergic). Given that timing of complex spikes depends on the precise internal state of olive neurons, which oscillate at 4-5 Hz (Khosrovani et al., 2007), and plasticity in P-cells depends on timing of the complex spikes (Herzfeld et al., 2018; Suvrathan et al., 2016), a timing difference may not be a good solution.

A different way to consider puzzle P1 is to note that a single P-cell projects to only 4-5 individual DCN neurons (Person and Raman, 2012b), and critically, this small group of DCN neurons includes both GABA-ergic, as well as non-GABA-ergic neurons (De Zeeuw and Berrebi, 1995; Teune et al., 1998). Therefore, for each GABA-ergic DCN neuron that projects to the olive with activity 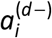, there are one or two non-GABA-ergic DCN neurons that receive input from the same P-cell. In order for the non-olive projecting DCN neuron 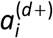 to receive the appropriate error signal, it can pair its activity with a “sister” DCN neuron that projects to the olive 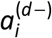. The two sister DCN neurons would have to receive inputs from the same P-cells and mossy fibers so that their activities are similar (Fig. 1C).

If a relationship between a pair of non-GABA-ergic and GABA-ergic DCN neurons existed such that 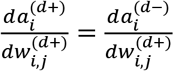, then we can compute the gradient of the loss function with respect to the weights of the non-olive projecting DCN neurons:

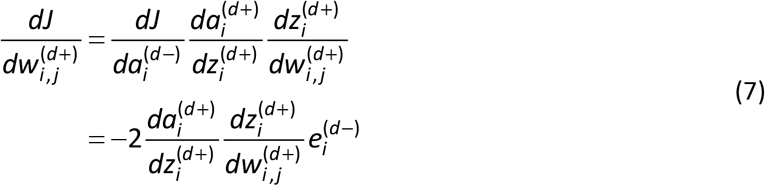

The above expression implies that a non-olive-projecting DCN neuron should share the error signal with its “sister” olive-projecting DCN neuron. From this equation we arrive at our second conjecture:

**Conjecture C2**. *The prediction error signal conveyed by an olive neuron must result in plasticity in at least one olive projecting GABA-ergic neuron, and one non-olive projecting, non-GABA-ergic neuron. These two DCN neurons should be coupled in the sense that their activities should be roughly equal at all times*.

At this writing, we do not know how the non-GABA-ergic DCN neurons receive their error information. Conjecture C2 provides the possibility that if the activities in the two DCN neurons were identical, then the olivary signal that reflects error for one DCN neuron can also be the signal that teaches the other DCN neuron. Unfortunately, even if conjectures C1 and C2 were true, we still have puzzle P2: how can a DCN neuron learn about its error when it does not receive an effective signal from the olive?

The trivial answer is to suggest that perhaps DCN neurons have essentially static weights and internal biases and do not learn from their errors. But this is clearly not the case, as evidenced by the work of Mauk and colleagues (Ohyama et al., 2003; Ohyama and Mauk, 2001), De Zeeuw and colleagues (Boele et al., 2013; Carulli et al., 2020), and Nagao and colleagues (Shutoh et al., 2006). For example, in a classical conditioning task in which animals hear a tone and learn to predict the arrival of an aversive stimulus and close their eyes, training produces plasticity in the P-cells (Jirenhed and Hesslow, 2016), as well as the DCN, as evidence by the fact that after conclusion of training, disconnection of the P-cell input to the DCN (via GABA antagonists) produces the conditioned behavior, albeit at an earlier time with respect to the tone onset (Medina et al., 2001). Furthermore, this training coincides with mossy fiber axonal growth and synaptic genesis in the DCN (Boele et al., 2013), as well as changes to the extra-cellular matrix of molecules that surround the synapses that contact DCN neurons (Carulli et al., 2020). Thus, experience of error leads to plasticity in the P-cells as well as the DCN. But given the weak olivary input to the DCN, how is this possible?

Let us consider two scenarios. On the one hand, perhaps error information is not conveyed from the olive to the DCN, but from the simple spikes of P-cells to the DCN. In this scenario, the olive does not play a role in computing error information for the purpose of teaching the DCN. Rather, the P-cells indirectly receive error information from the mossy fibers, and through their converging simple spikes teach their downstream DCN neurons.

Indeed, the synapses that P-cells make upon a DCN cell can undergo plasticity based on the history of the P-cell’s simple spikes (Telgkamp and Raman, 2002). Furthermore, the synapses that mossy fibers make upon a DCN neuron can also undergo plasticity: when a period of excitation (150 ms) in the DCN neuron precedes a period of strong inhibition (250 ms), excitatory synapses strengthen (McElvain et al., 2010; Person and Raman, 2010; Pugh and Raman, 2008). Thus, the sequential pattern of excitation from the mossy fibers and inhibition from the P-cells can, in principle, serve as a teacher for changing the synaptic weights of the inputs onto DCN neurons (De Zeeuw et al., 2011).

Alternatively, it is possible that error information is conveyed from the olive to a population of P-cells via climbing fibers, and then through synchrony of complex spikes the P-cells convey occurrence of the error event to the DCN. In this scenario, climbing fiber activity not only guides plasticity in the P-cells, but also organizes the P-cells anatomically into populations that share a common error signal. The resulting temporal synchrony in the complex spikes produces a period of suppression in the DCN neuron, which may result in plasticity in the mossy fiber and P-cell synapses that act on that DCN neuron. Let us consider these two possibilities in detail.

## Possibility 1: transmitting error information to the deep nucleus via simple spikes

Recent results have demonstrated that modulation of P-cell simple spikes alone can drive learning in the cerebellum. Nguyen-Vu et al. (2013) considered the vestibular ocular reflex (VOR), a behavior in which a vestibular input is repeatedly paired with a visual stimulus. Usually, P-cells in the flocculus have simple spikes that reflect both the vestibular stimulus and the visual feedback, while complex spikes reflect retinal error. The authors hypothesized that modulation of simple spikes alone could produce learning in the cerebellum independent of complex spikes. To test for this, they used optogenetics in mice to stimulate P-cell activity, generating simple spikes that were paired with the vestibular stimulus in the absence of visual input. During training, a sinusoidal vestibular stimulus was used in the dark while each side of the cerebellum was stimulated during ipsiversive or contraversive turning. Pairing optogenetic pulses of stimulation with ipsiversive turning produced learning that reduced the VOR gain. Similarly, pairing optogenetic stimulation with contraversive turning produced learning that increased the VOR gain. Because optogenetic pulses likely synchronized production of simple spikes across populations of P-cells, the work illustrated that production of synchronous simple spikes was sufficient to produce learning.

How did this learning compare with complex spike driven adaptation? Nguyen-Vu et al. (2013) paired stimulation of climbing fibers with a vestibular stimulus and found that this form of stimulation also produced learning, but the patterns were somewhat different than direct P-cell stimulation. Like simple spike stimulation, ipsiversive turning paired with climbing fiber stimulation increased the VOR gain, but climbing fiber stimulation during contraversive turning did not reduce the VOR gain. Thus, activation of P-cells and production of synchronous simple spikes could produce motor learning, a role that had generally been ascribed to complex spikes.

The results of Lee et al. (2015) extended this work, demonstrating that synchronized patterns of simple spikes produced DCN plasticity. Using optogenetic pulsed stimulation of P-cells in the forelimb region of the anterior cerebellum of mice, they found that onset of P-cell inhibition was followed by an arm movement (raising the arm upward), whereas offset of a period of P-cell excitation produced a similar movement. They then paired excitation or inhibition with an auditory tone and made a remarkable observation: the animal raised its arm when the tone was presented without P-cell stimulation. Notably, optogenetic stimulation of P-cells induced strong synchrony in their simple spike activities, and this coincided with structural changes in the mossy fiber inputs to the DCN. Thus, synchronized P-cell simple spikes led to plasticity in the DCN.

Ke et al. (2009) asked whether learning could be induced with error driven changes in simple spikes but without significant changes in complex spikes. They trained monkeys in the VOR task by rotating their heads but manipulated the visual input so that there was a difference in how the visual target moved as compared to how the background visual scene moved. This novel stimulus appeared to reduce the complex spike response to error (retinal slip). However, the simple spike response with the novel stimulus resembled the response during natural gain-down training. Following one or two hours of training with the novel stimulus they assessed learning by measuring eye movements in response to head movements in darkness and found that the VOR gain had declined. Using a linear model they fitted the amount of behavioral adaptation in the various paradigms as a function of changes in the complex and simple spikes. Whereas changes in complex spikes accounted for roughly 65% of changes in behavior, roughly 35% of adaptation appeared to be driven by changes in simple spikes.

Correlates of an error-like variable in simple spikes were first noted by Kase et al. (1979) who trained monkeys in two tasks, one in which they actively pursued a sinusoidally moving dot with their eyes, and another in which they fixated but the dot moved sinusoidally. They recorded from P-cells along the vermis of lobules VI-VII and found that 14/89 cells showed simple spike modulation both during active pursuit, and during fixation as the target moved with respect to the fovea.

More recently, correlates of error were found in simple spikes as monkeys were trained to use a robotic arm to move a cursor and track a moving target that traced a random path. Ebner and colleagues (Hewitt et al., 2015, 2011; Popa et al., 2012; Streng et al., 2018) recorded from hundreds of P-cells in the intermediate and lateral regions of lobule IV-VI and used linear regression to first account for the variance in the simple spike rate with respect to motion of the arm, and then determined how much of the residual simple spike rate could be associated with error, that is, the instantaneous distance between the hand-held cursor and the target. The regression with respect to error generally exhibited two peaks: one at a lead (−223 ms prediction about a future error), and the other at a lag (+227 ms feedback about the past error) (Popa et al., 2017). The r-squares for the delayed response of simple spikes to error were greater than the r-squares for the prediction of future error. The coefficient of the linear fit that predicted the error was often negative of the coefficient that responded to that error.

To check that the simple spikes were indeed responding to visual error and not other confounding variables, Streng et al. (2018) introduced a delay between position of the cursor and the position of the hand. When the cursor was not delayed, simple spikes were correlated with error at approximately −250 ms and at +200 ms. However, in the visual delay block, the peak of the correlation in the negative range shifted by 100 and 200 ms, but did not shift in the positive range. This means that the simple spikes predicted hand position (not cursor position) with respect to the target with a lead of around +250 ms. Importantly, simple spikes responded to error with a 200 ms delay and this delay remained stable even when there was an artificial delay in cursor feedback. The results provided further evidence that following simple spikes were modulation following an error event.

Thus, simple spikes not only can carry error information, but that information is also sufficient to induce adaptation in the DCN, particularly if the simple spikes are synchronized. However, Eq. (5) and Eq. (6) impose a stringent requirement: the network must transmit the error that has been made by a single DCN neuron specifically to that DCN neuron. This means that if the P-cells that converge on a DCN neuron carry error information, that information must be the difference between the DCN neuron’s output and the desired output. At this writing, it is unclear how the P-cells through their simple spikes might convey this specific kind of error signal.

## Possibility 2: transmitting error information to the deep nucleus via complex spikes

If we assume that the olivary information regarding the error that the DCN neuron has made is not conveyed to the DCN neuron directly (via a projection from the olive), we are left with the possibility that the nucleus neuron becomes “aware” of its error through an indirect pathway. The obvious indirect pathway is from the parent P-cells (Lang et al., 2017; Medina, 2011).

A complex spike is a significant event at the soma of a P-cell, generating multiple spikelets. However, the P-cell’s axon transmits this event to the DCN via ordinary spikes (Ito and Simpson, 1971): typically one axonal spike at the onset of the complex spike (with near 100% probability) and another one at the final somatic spikelet (about 60% probability) (Khaliq and Raman, 2005; Monsivais et al., 2005). Thus, the information in the climbing fiber is transmitted from a P-cell to the DCN as a sequence of one or two ordinary spikes. This makes it unlikely that a DCN neuron could dissociate between a complex spike and a simple spike. If synchronous simple spikes can produce plasticity in the DCN (Lee et al., 2015), perhaps synchronous complex spikes can also accomplish this feat.

When the inferior olive is electrically stimulated, at 5-10 ms latency there is occasionally a single spike in a DCN neuron (Bengtsson et al., 2011; Hoebeek et al., 2010). This is likely due to the direct excitatory projections from the olive to the DCN. This occasional spike is then followed by suppression of activity that lasts around 50 ms (Fig. 2) (which can be followed by a rebound burst of activity). The suppression is a consistent feature in many DCN neurons, including output neurons that do not project to the olive (e.g., non-GABA-ergic neurons that project to the red nucleus) (Hoebeek et al., 2010). If we view olivary stimulation as an artificial means for providing error feedback to the cerebellum, then that error leaves its impression not via a strong EPSC in the DCN neuron at the time of error, but surprisingly, via a strong suppression of activity during a short period that follows.

**Figure 2.**
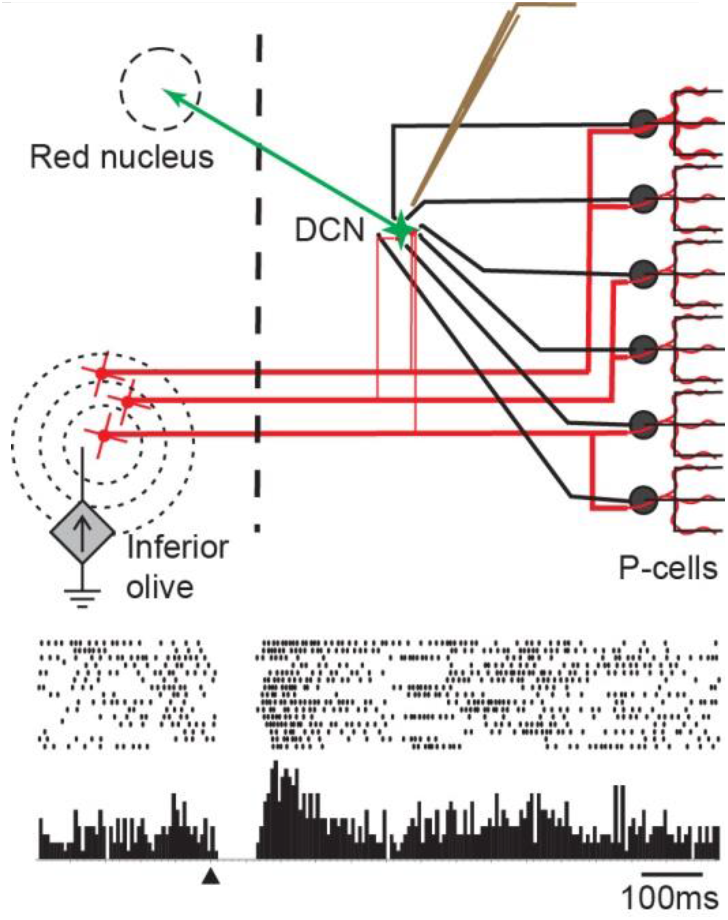
Effect of inferior olive stimulation on DCN neurons (interposed nucleus) in an anesthetized rat. Neurons in the inferior olive were stimulated while an electrode recorded activity of an excitatory neuron that projected to the red nucleus. Stimulation of the inferior olive produced a 50 ms pause in the activity of the DCN neuron. From Hoebeek et al. (2010), used by permission.

A series of experiments by Lang and colleagues (Blenkinsop and Lang, 2011; Tang et al., 2019, 2016) provided critical clues regarding how the activity in the olive produced the suppression in the DCN. Blenkinsop and Lang (2011) inserted an array of electrodes into the anesthetized rat’s cerebellar cortex along with a single electrode into the DCN. This allowed them to simultaneously record from 8-34 P-cells and 1-2 DCN neurons. In order to determine if one of the P-cells was connected to one of the DCN neurons, they identified complex spikes in each P-cell and used that event to align the spikes recorded from the DCN neuron. The result was a complex spike-triggered response in the DCN neuron (Fig. 3A).

**Figure 3.**
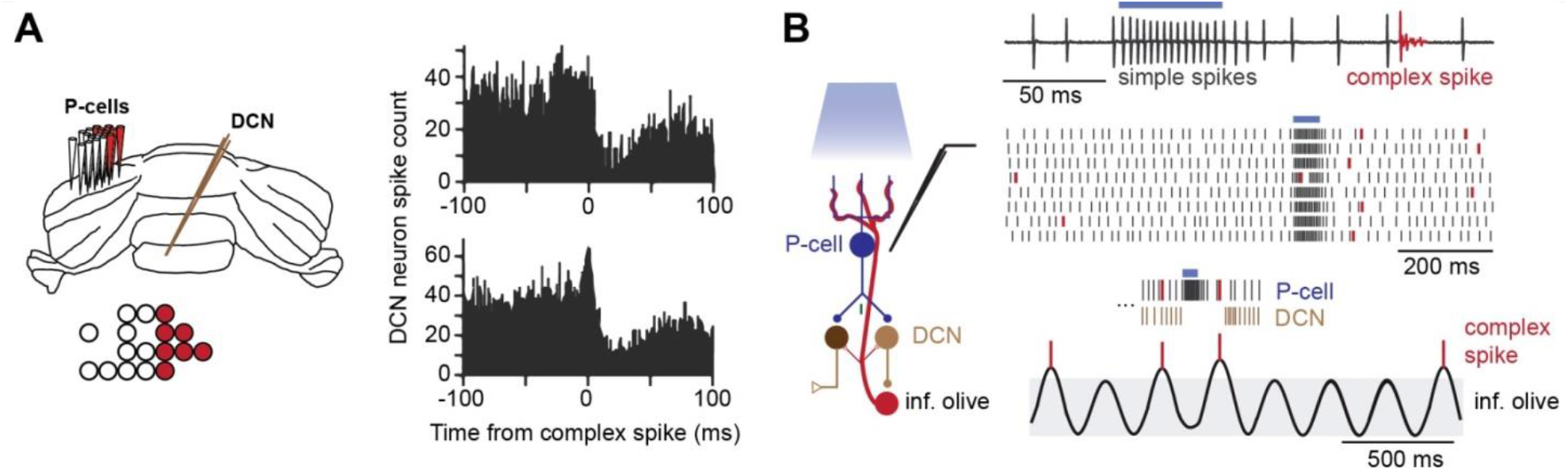
An olivary neuron that projects to a P-cell receives input from the DCN daughter neuron of that P-cell. **A**. Electrode array recordings from multiple P-cells and a single DCN neuron in an anesthetized rat. To determine if a P-cell projected to the DCN neuron, the complex spike in the P-cell was used as the trigger to align the spikes in the DCN neuron. The data on the right are from two instances in which the spike-triggered averaging suggested a P-cell to DCN projection. In 70% of the pairs the complex spike in the P-cell was followed by reduced activity in the DCN neuron (top right). In 21% of the pairs there was an initial spike in the DCN neuron followed by reduced activity. The P-cells that appeared connected to a single DCN neuron were usually clustered together along the rostra-caudal axis of the cerebellum, as shown on the left with red-filled electrodes in the array. From Blenkinsop and Lang (2011) and Tang et al. (2019), used by permission. **B**. Optogenetic stimulation of P-cells is followed by production of a complex spike. P-cells were optogenetically stimulated while activity was recorded with an electrode. Stimulation resulted in intense production of simple spikes, which was then followed at a latency of about 100 ms with a complex spike. A model suggests that P-cell stimulation strongly suppresses the DCN neuron, which in turn removes inhibition from the olivary neuron. The olivary neuron has a membrane potential that oscillates. Removal of inhibition allows the potential to reach threshold sooner, resulting in climbing fiber activity in the parent P-cell, and thus a complex spike. From Chaumont et al. (2013), used by permission.

In about 100 cases the spike-triggered response suggested that one of the P-cells was connected to one of the DCN neurons. In 70 of the 100 putative P-cell DCN neuron pairs the occurrence of a complex spike was followed by a suppression in the DCN, a suppression that lasted from 50-100 ms (Fig. 3A, right top subplot). In 24 pairs, the complex spike in the P-cell was coincident with a brief increase in firing rate of the DCN neuron, and then suppression (Fig. 3A, right bottom subplot). The authors interpreted the small increase as the effect of the weak projections from the olive to the nuclear neuron. Therefore, a single axon from the olive could project both to a DCN neuron, and to the P-cell that synapsed on that DCN neuron (Fig. 1C).

If the error associated with a DCN neuron’s output is conveyed to it via its parent P-cells, then we should observe complex spikes in the parent P-cell following an error by the DCN neuron. There is indirect evidence for this from optogenetic stimulation of P-cells. A 50 ms period of P-cell stimulation results in synchronized production of simple spikes, but when the P-cell stimulation ends, at 80-150 ms latency the P-cells produce a complex spike (Chaumont et al., 2013) (Fig. 3B, top subplot). The interpretation is that when many P-cells are simultaneously activated, they inhibit their downstream DCN neurons synchronously, preventing these neurons from firing. This suppression of the DCN neuron’s activity removes a source of inhibition at the olive. The removal of that inhibition allows the olivary cells to reach threshold earlier and fire, resulting in a complex spike in the P-cells (Fig. 3B, bottom subplot). This interpretation is consistent with a closed loop in which P-cells project to a DCN neuron, which then projects to an olive neuron, which then projects back to the parent P-cell (Fig. 3B), termed an olivocerebellar module (De Zeeuw et al., 1997).

These results suggest the following anatomy (Heck et al., 2013): the olive neuron that computes the error associated with a GABA-ergic DCN neuron’s output conveys that error via a single climbing fiber to a P-cell. That P-cell then projects back to that specific GABA-ergic DCN neuron, which then projects to the olive neuron that computed the error (Fig. 3B).

**Conjecture C3.** *The error associated with a GABA-ergic DCN neuron’s output is computed by an olive cell and then sent via climbing fibers to a few P-cells which then project back down to that specific DCN neuron*.

This conjecture describes an olivocerebellar module (De Zeeuw et al., 1997), and is the result of roughly three decades of anatomical observations (Ruigrok and Voogd, 2000). A single axon from the olive projects to approximately 7 P-cells, all located at similar distances from the midline, most of which are clustered within a single lobule (Sugihara et al., 1999, 2001). Molecular markers, notably the respiratory enzyme aldolase C (commonly known as zebrin II), define about 20 longitudinal compartments in the cerebellar cortex, each receiving inputs from specific regions of the inferior olive. P-cells that are located in the same compartment, and have similar climbing fiber receptive field characteristics, project to the same small area in one of the DCNs (Apps and Garwicz, 2000). Indeed, Lang and colleagues (Tang et al., 2019) noted that the P-cells that together projected to a single DCN neuron were not randomly distributed, but clustered along the rostra-caudal axis of the cerebellum (Fig. 3A). Tracing studies have identified specific projections from the olive to each compartment in the cerebellar cortex (Apps and Garwicz, 2005). The olive projects to both GABA-ergic and non-GABA-ergic neurons in the DCN (De Zeeuw et al., 1997). There are no molecular markers that have thus far shown compartmentalization of the DCN. However, DCN regions are divided based on tracing of inputs from the olive (Sugihara & Shinoda, 2007). P-cells that are in a given compartment, and thus receive an input form a region in the olive, project to a region in the DCN that also receives inputs from the same olive region (Sugihara et al., 2009). Finally, as we noted in conjecture C1, olivary projections to the DCN are reciprocal: if a region in the olive projects to a region in the DCN, that DCN region also project back to that specific region of the olive (Ruigrok and Voogd, 2000). Thus, we have the anatomical basis of the closed olivo-cortico-nuclear circuit that connects a longitudinal compartment of P-cells to a small subarea in the DCN (Sugihara, 2011).

In our theoretical framework, this anatomy serves a critical purpose: by synchronizing the complex spikes in the P-cells, the olive provides the DCN neurons with a reliable, life-long error signal, despite the fact that the olivary input to the DCN weakens with development (Najac and Raman, 2017), and may not play a significant role in adulthood (Lu et al., 2016).

Medina and Mauk (1999), building on ideas put forth by Miles and Lisberger (1981), had conjectured that whereas the climbing fiber input to the P-cell controls plasticity of parallel fiber inputs to that cell, it is the P-cell that controls the plasticity of mossy fiber inputs to its downstream DCN neuron. For this to happen, the mathematics require the P-cell to somehow communicate to its DCN neuron the fact that it has received an input in its climbing fiber. The results in Figs. 2 and 3 show that the signal from the olive to a P-cell can have a dramatic effect on the firing rates of nucleus neurons: following a single complex spike, there can be a 50 ms reduction in the firing rate of the DCN neuron. To be sure, some of these results are in the anesthetized state. Do they generalize to the awake, behaving animal?

ten Brinke et al. (2017) trained mice to associate an LED with an air puff and recorded from the interposed nucleus. During training, most task-related DCN neurons increased their discharge in response to the LED, and this increase was causally related to production of the eye-blink. The LED onset, which was a random event (thus producing a prediction error) and the air puff, both tended to produce a single complex spike in a few P-cells (Fig. 4A). However, in DCN neurons, the LED onset and especially the air puff tended to produce a transient pause in spiking (Fig. 4B). The DCN neurons that produced the air puff-induced pause also tended to show greater facilitation in their spiking during the LED period, presumably driving the eye-blink response. Thus, during learning of a behavior, some DCN neurons showed a transient pause that appeared related to arrival of a complex spike in P-cells. However, we do not know if the P-cells in Fig. 4A project to the specific DCN neurons shown in Fig. 4B. Thus, we do not know if in this data set there is a causal relationship between P-cell complex spikes and suppression in DCN activity.

**Figure 4.**
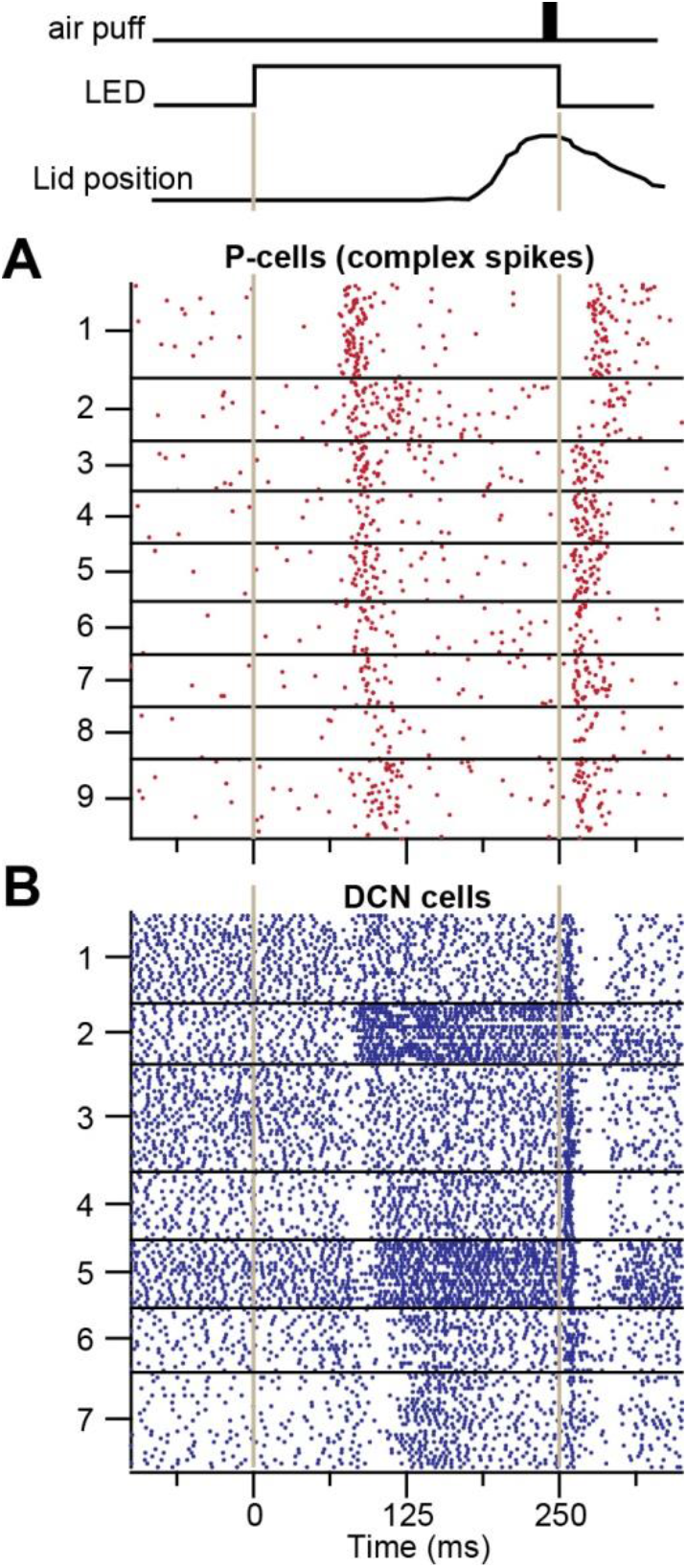
During learning of a behavior, some DCN neurons show a transient pause that appear related to arrival of error information via a complex spike in their parent P-cells. Recordings are from P-cells and neurons in the interposed nucleus as mice learned to associate an LED with an air puff. The LED onset and the air puff both tended to produce complex spikes in P-cells, which coincided with a transient pause in DCN neurons. From ten Brinke et al. (2017), used with permission.

Results of Tang et al. (2019) provide the needed evidence. They noted that in anesthetized animals, the P-cells that projected to a single DCN neuron tended to have higher levels of complex spike synchrony than P-cells that did not project to the same neuron (Fig. 5A). Indeed, when they simultaneously recorded from 6 or 7 P-cells that were putatively connected to a single DCN neuron, they found that if the complex spikes in the parent P-cells were not synchronous, a single complex spike by itself had little or no effect in the DCN neuron (Fig. 5C, subplot in which 1/7 P-cells had a complex spike). However, if by chance the complex spikes in some of the parent P-cells were synchronous, their simultaneous convergence produced a suppression (Fig. 5C, 4/7 P-cells had a complex spike). In the rare event that all 7 of the recorded P-cells happened to produce a complex spike simultaneously, the DCN neuron’s suppression was strong and long lasting (Fig. 5C, 7/7 condition).

**Figure 5.**
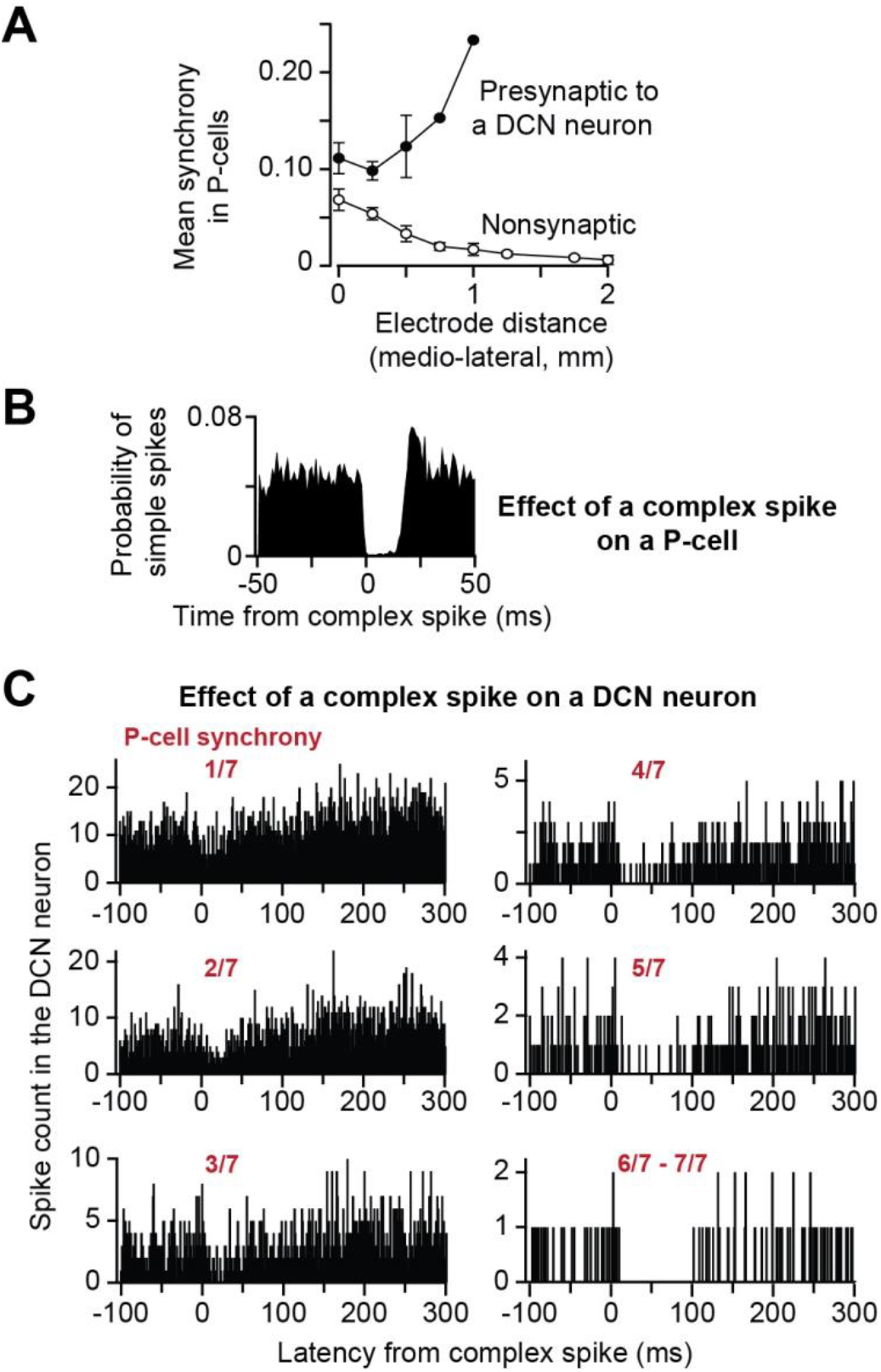
Effective transmission of the error signal to the nucleus requires complex spike synchrony. **A**. Groups of P-cells that project to a single DCN neuron tend to have higher levels of complex spike synchrony than groups of P-cells that do not project to the same neuron. From Tang et al. (2019), used with permission. **B**. Following a complex spike, there is a 10-20 ms period of total simple spike suppression in the P-cell. Data are from awake behaving marmoset. From Sedaghat-Nejad et al. (2019), used with permission. **C**. Effect of a complex spike is greater if the event is synchronized among the parent P-cells. From Tang et al. (2019), used with permission.

Note the similarity between the effect of a complex spike on a P-cell, and its effect on a DCN neuron: following a complex spike, both the P-cell (Fig. 5B) and the DCN neuron that it projects to (Fig. 5C) can experience a long-lasting suppression. In the case of a P-cell, a complex spike is followed by 10-20 ms of total suppression of simple spikes (Fig. 5B). This is partly due to the fact that a climbing fiber that innervates a P-cell also sends an axon collateral to a nearby molecular layer interneuron, which in turn inhibits that P-cell (Jörntell and Ekerot, 2002). In the case of a DCN neuron, a complex spike in the parent P-cell is followed by a reduction in the firing rate of the nucleus neuron, lasting about 50 ms, but only if that there is temporal synchrony among the complex spikes.

We are now in position to offer a tentative solution to puzzle P2.

**Conjecture C4.** *A DCN neuron is made aware of its erroneous output via synchronous complex spikes in the population of P-cells that project onto it*.

Around 50 P-cells project to a single nucleus neuron (in mice) (Person and Raman, 2012a). If these P-cells shared a common error signal from the olive, then they could solve an important problem: through simultaneity of their complex spikes, the P-cells could reliably transmit error information to the output layer.

There is evidence for the idea that unexpected perturbations increase synchrony of complex spikes, and that synchronous complex spikes are essential for normal learning. Van Der Giessen et al. (2008) produced mice that lacked gap-junctions, which are prominently expressed in the inferior olive. Gap-junction deficient mice showed impaired ability to learn to time their eye blink response to the end of a tone (at which point an air puff was directed to the eyes). Instead, the mice tended to close their eyes in response to the onset of the tone. The authors examined complex spike timing in the P-cells of lobule VI in the posterior lobe and found that in response to an air puff, the wildtype mice produced a single complex spike that was timed at around 30 ms average latency. In contrast, in the gap-junction deficient mice the air puff produced complex spikes that were distributed more widely in time with two peaks, one at 30 ms and another at 101 ms latency. Later work demonstrated that lack of gap junctions reduced the probability that complex spikes would be produced synchronously among P-cells in response to a perturbation during locomotion (De Gruijl et al., 2014).

In summary, it is possible that the olivocerebellar module, i.e., the loop from a P-cell to a DCN neuron to an olivary neuron back to the same P-cell, serves a fundamental purpose: to make it possible for the olive to convey the error in the output of a DCN neuron to a small group of P-cells, thus generating synchronous complex spikes which can then suppress activity in the DCN neuron that produced the erroneous output.

## Plasticity in the DCN

De Zeeuw et al. (2011) had conjectured that the complex spike induced suppression of activity in DCN (and the rebound that followed in some DCN neurons) played a key role in control of DCN plasticity. Indeed, Raman and colleagues (Person and Raman, 2010; Pugh and Raman, 2008) had observed that a sequential pattern of excitation followed by inhibition in a DCN neuron produced long term potentiation (LTP) in the mossy fiber synapses: when a period of excitation (150 ms) in the DCN neuron preceded a period of strong inhibition (250 ms), mossy fiber synapses strengthened. Similarly, in the medial vestibular nucleus (a DCN analogue for some regions of the cerebellar cortex), vestibular nerve synapses (a mossy fiber analogue) on the vestibular nucleus neurons exhibited LTP when nerve stimulation coincided with nucleus neuron hyperpolarization (McElvain et al., 2010). In addition, Zhang and Linden (2006) had found that a period of mossy fiber excitation alone led to long term depression (LTD) of the mossy fiber synapses.

Thus, we have a potential mechanism with which the error in the output of a DCN neuron can cause plasticity in that specific DCN neuron: 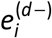 is detected by an olive cell and sent via climbing fibers to a population of P-cells whose synchronous complex spikes produce a long-lasting suppression of spiking in the DCN neuron. If that suppression is preceded by a period of excitation from mossy fibers, the result is an increase in the weight of the mossy fiber synapses on the DCN neuron. On the other hand, without this suppression there may be a decrease in the weight of the mossy fiber synapses.

Complex spike induced suppression of spiking in the DCN neuron must also affect the strength of the P-cell synaptic inputs. Mechanisms of plasticity in the P-cell synapses in the nucleus are poorly understood, but there is some evidence that this plasticity depends on activation patterns that resemble the suppression of DCN activity that follows a complex spike. Aizenman et al. (1998) found that in slices from the cerebellum of juvenile rats, following 200 ms of hyperpolarization that suppressed the nuclear cell’s spiking, there was rebound depolarization (i.e., a burst of spikes). The amount of this rebound depolarization dictated the direction of plasticity in the P-cell synapse: if the rebound was missing or weak, the synaptic strength was reduced. It is noteworthy that in vivo, generation of a complex spike in the parent P-cell not only leads to a period of suppression of spiking in the DCN neuron, this suppression is occasionally followed by a burst of spiking (Hoebeek et al., 2010). Results of Aizenman et al. (1998) imply that the magnitude of this rebound depolarization influences direction of weight change in the P-cell synapses upon the DCN neuron.

In summary, DCN neurons provide the output of the cerebellum and must be made aware of their errors. However, most of these neurons do not project to the olive, one of the locations where error may be computed. Furthermore, those that do project to the olive receive olivary projections that progressively weaken through development. As a result, at present we do not know how a DCN neuron learns to change its erroneous output. However, synchronous complex spikes in the parent P-cells can produce a 50 ms period of suppression in the DCN neuron. If the suppression is preceded by a period of excitation, the synaptic inputs from the mossy fibers to the DCN neuron strengthen. If the suppression is followed by rebound spiking, the synaptic inputs from the P-cells to the DCN neuron strengthen. Thus, control of plasticity at the DCN may be via synchronous complex spike activity among the population of P-cells that project onto it, providing an indirect pathway that communicates the error information from the olive to the DCN neurons.

## Membership criterion for P-cell populations

We are now in position to make a guess regarding the membership criterion for the P-cell population: in order to transmit the error signal reliably from the climbing fiber to the DCN, there is a need for complex spike synchrony, which is more likely if the population of P-cells that project to a given DCN neuron share a similar error signal from the olive. Furthermore, for the error signal to be a teacher for the downstream DCN neuron, it must be generated by olivary neurons that compute the error in that DCN neuron’s output.

The ideal scenario (from a computational perspective) would be the following. Suppose that an olive neuron computes the error associated with a given DCN neuron, i.e., receives an inhibitory synapse from that DCN neuron. The olive neuron’s axon to the cerebellum should split and synapse on every one of the 50 P-cells that project to that DCN neuron. However, a single olivary axon projects to only about 7 P-cells (Sugihara et al., 1999, 2001). Therefore, the population of P-cells that project to a nucleus neuron is bigger than what a single olive neuron can serve.

A typical DCN neuron receives inputs from approximately 8 olivary axons (Najac and Raman, 2017). If we imagine that each of these axons is a collateral from an olivary axon that travels to the cerebellar cortex and becomes the climbing fiber for 7 P-cells, then the olive to nucleus convergence ratio (8:1) combined with the olive to P-cell divergence ratio (1:7) produces roughly 56 P-cells that receive information about the error made by a single nucleus neuron. In a remarkable coincidence, about 50 P-cells converge on a single nucleus (in mice) (Person and Raman, 2012a). We thus arrive at our central conjecture regarding the membership criterion for P-cells that form a single population (Fig. 6A):

**Figure 6.**
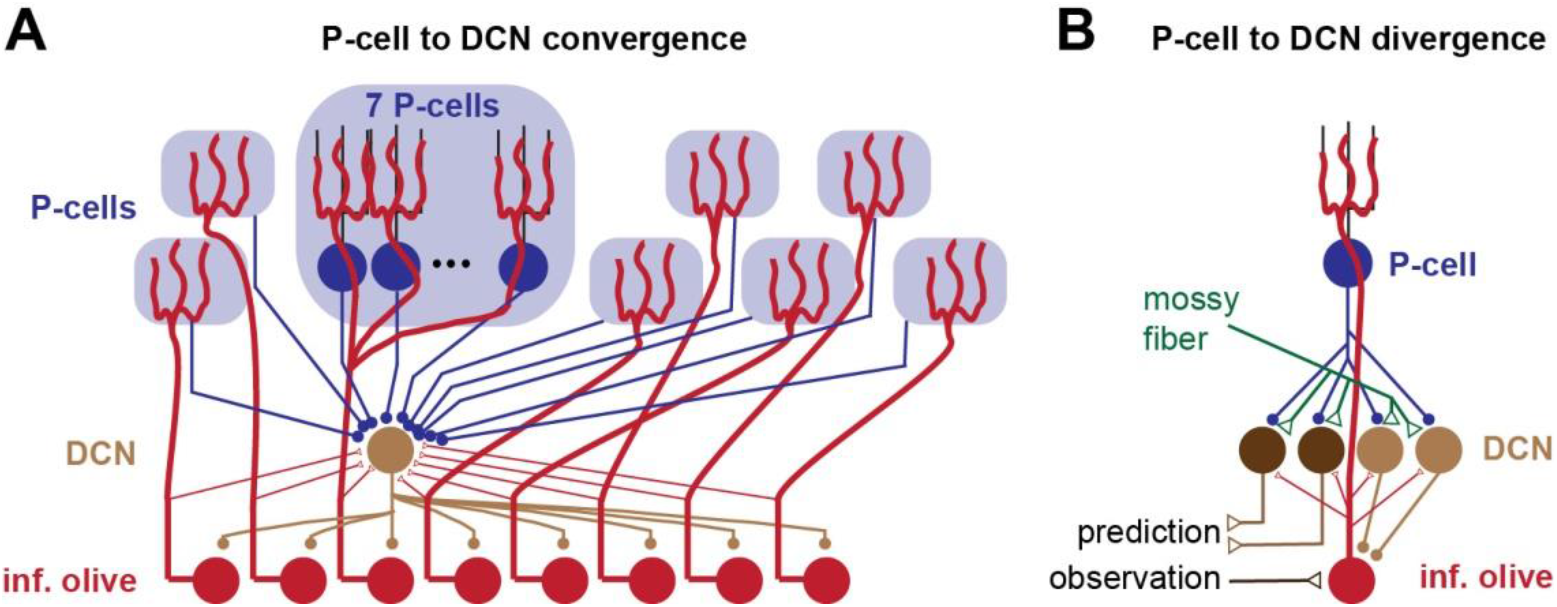
Membership criterion for population coding in the cerebellum. **A**. The population of P-cells that project onto a nucleus neuron is composed of those P-cells that receive climbing fibers from the olivary neurons that also project to that specific nucleus neuron. **B**. A single P-cell projects to only a few DCN neurons, which include both GABA-ergic and non-GABA-ergic cells. Efficient learning requires that the error signal that the P-cell receives represent these errors only. Here, the GABA-ergic DCN daughters of a given P-cell converge their axons on a single olive neuron, and that olive neuron serves as the teacher for the parent P-cell, providing it with its single climbing fiber. The olive cell in term sends excitatory projections to all the DCN daughters of the P-cell.

**Conjecture C5**. *The population of P-cells that project onto a DCN neuron is composed of those P-cells that receive climbing fibers from the olivary neurons that also receive projections from that specific nucleus neuron*.

In summary, a handful of olivary cells send axons to a single nucleus neuron, and each olivary cell also sends climbing fibers to a handful of P-cells. Perhaps through development, the olivary cells guide those P-cells to project to the same nucleus neuron (Najac and Raman, 2017). In this scenario, the P-cell population that projects to a single DCN neuron would consist of P-cells that receive a climbing fiber from one of those olivary cells. If most of these olivary cells also receive a projection from that DCN neuron, then the olivary cells are in position to compute the error (Eq. 1) in the DCN neuron’s output and provide that information via climbing fibers to the 50 P-cells that represent the parent of the DCN neuron. The P-cells would transform the error information into a complex spike, which if synchronized with other P-cells in the population, would result in a brief suppression of the DCN neuron’s activity that was responsible for the error. This in turn would produce plasticity in the DCN neuron’s mossy fiber and P-cell synapses. If these conjectures are true, then the responsibility to teach a DCN neuron would be relegated from the olive to the population of P-cells that converged on that DCN neuron.

## An appropriate error signal for a P-cell

The next step is to ask how we should teach the neurons in our middle layer, i.e., the P-cells. This may seem like a trivial task because each P-cell receives a climbing fiber, thus endowing it with a powerful channel to receive error information. Indeed, both the presence and absence of complex spikes synchronous with simple spikes produces parallel fiber plasticity (Jörntell and Ekerot, 2002). However, if we consider this question mathematically, we arrive at another useful inference.

Due to its placement in the middle layer, a P-cell does not have a direct role in producing an output from the cerebellum, and is not directly responsible for any error. Rather, it projects to a handful of DCN neurons that produce outputs, and thus contribute to our loss function (Eq. 1). Importantly, because a P-cell projects to only 4 or 5 DCN neurons (Person and Raman, 2012b), it contributes to errors made by only these specific output neurons. The mathematics require that the climbing fiber that the P-cell receives must carry error information related to activity of the DCN neurons that it projects to.

If this were an artificial neural network, we would send a P-cell a distinct error signal associated with each of the DCN neurons it projects to. However, a P-cell has only a single climbing fiber. Thus, we face a puzzle in how to teach the P-cells.

**Puzzle P3.** *A P-cell contributes to activity of multiple DCN neurons. Yet, it receives only a single climbing fiber. How can the P-cell learn to change its activity in order to reduce the errors associated with the DCN neurons it projects to?*

Suppose that P-cell *k* has activity 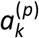, where the superscript refers to the fact that this unit is a P-cell. This P-cell projects to neurons *j* = {1, ⋯, *N*} in the DCN layer, contributing to their activities 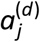. As a result, the gradient of the loss function with respect to activity of this P-cell is:

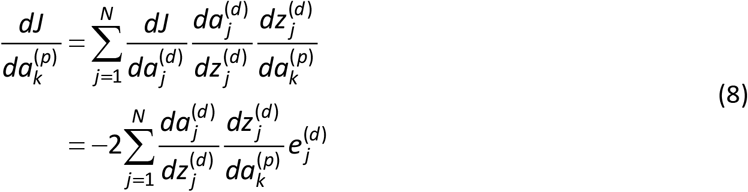

The above expression implies that the error that the P-cell must learn from is proportional to the weighted sum of the errors made by its downstream DCN neurons. Unfortunately, the P-cell receives only a single climbing fiber. Thus, this climbing fiber must represent the errors made by all the DCN neurons *j* = {1, ⋯, *N*} that the P-cell projects to.

Among the 4 or 5 DCN neurons that a single P-cell projects to, some are GABA-ergic, projecting to the olive, and some are non-GABA-ergic, projecting elsewhere in the brain (Teune et al., 1998). A potential solution to our puzzle is an anatomy in which the GABA-ergic DCN neurons of a given P-cell converge their axons on a single olive neuron, and that olive neuron serves as the teacher for the parent P-cell, providing it with its single climbing fiber (Fig. 6B). Alternatively, the GABA-ergic DCN neurons of a given P-cell may converge their axons on different olive neurons that are electrically coupled, one of which provides the climbing fiber to that P-cell.

For this to be an effective solution, the GABA-ergic and non-GABA-ergic DCN neurons of our single P-cell would have to be matched in their activity, which is indeed what we had conjectured earlier when we considered the problem of teaching the two groups of DCN neurons (conjecture C2). This wiring would raise the following requirement: the olive cell that provides the climbing fiber to a P-cell should be the same olive cell that receives inhibitory projections from every GABA-ergic DCN neuron that this P-cell projects to (Fig. 6B). This is consistent with the proposal that during development, the activity of the climbing fiber, and particularly the coincidence of complex spikes in the parent P-cell with the DCN neuron’s EPSCs from the olive, may be a guide in wiring the cerebellum (Lang et al., 2017; Lu et al., 2016).

In summary, efficient learning in a P-cell requires that its climbing fiber should come from an olive neuron that is the recipient of synapses from a specific group of DCN neurons: those that are “daughters” of that P-cell.

## Population coding of P-cells

Vernon Mountcastle used to start some of his lectures by reminding his students that “if you want to know what a neuron does, figure out what it is connected to.” That advice looms large in the motor system, where control is often via population coding: a few dozen neurons in a region of the brain project to a common output neuron, forming a population that encodes an aspect of behavior. In this population, the collective activity of all the neurons that are part of the population is important, not the activity of a single neuron. Unfortunately, we currently have no method that can label the membership of this population in the living brain. Instead, the common technique is to collect neurons into pseudo-populations based on the statistical properties of their discharge, for example, via principal component analysis, a technique in which one assigns a weight to each neuron’s output, and then labels the weighted sum of all activities as the population response (Churchland et al., 2010).

Our analysis here suggests that in the cerebellum, we can take an alternate approach, one that rests upon an *a priori* hypothesis about the role of the teacher (inferior olive) in organizing the students (the P-cells and their downstream DCN neurons). Our hypothesis states that P-cells that receive similar error signals are likely to be part of the same population (Fig. 6A). Recent work has tested this idea by organizing P-cells in the oculomotor vermis of macaques into populations that share a common teacher: a common complex spike tuning with respect to visual error. The results have demonstrated that whereas the simple spikes of individual P-cells may appear indecipherable, they become interpretable when the P-cells are organized into populations (Herzfeld et al., 2015, 2018).

Saccadic eye movements depend critically on the cerebellum, as evidenced by the deficits observed in patients with cerebellar damage (Leigh and Zee, 2015; Xu-Wilson et al., 2009) and monkeys with lesions (Barash et al., 1999; Kojima et al., 2010a). Given this evidence, it seemed likely that P-cell simple spikes should encode kinematic parameters of saccades (e.g., eye speed, amplitude, or direction). However, studies had found that individual P-cells showed little modulation of simple spikes with respect to saccade speed (Helmchen and Buttner, 1995) and direction (Thier et al., 2000). Indeed, simple spikes during saccades (Fig. 7A) presented a bewildering assortment of responses, including P-cells that produced a burst, P-cells that produced a pause, and P-cells that did both (Kojima et al., 2010b). Most puzzling was the fact that simple spikes were modulated well beyond saccade end (Thier et al., 2000), as shown in Fig. 7B.

**Figure 7.**
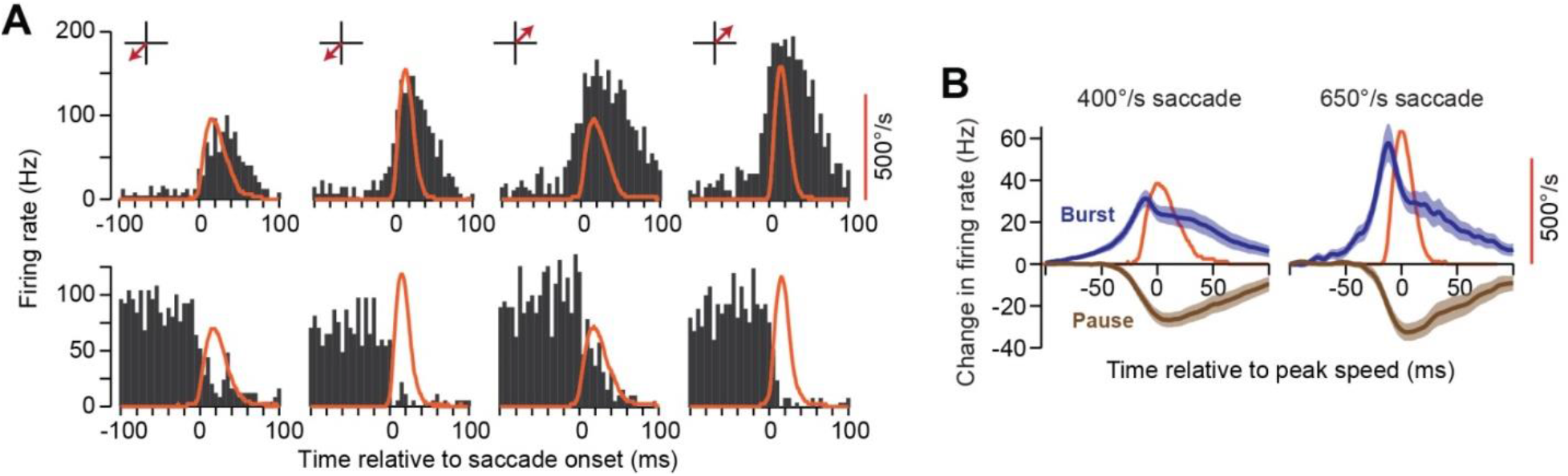
Activity of P-cells during a saccade. **A**. Saccades were made to various directions with various amplitudes and velocities. The red trace shows the eye velocity, and black bars indicate instantaneous firing rates of two P-cells. Each row is a single P-cell. Some P-cells exhibited a burst during the saccade, while others exhibited a pause. However, the duration of modulation tended to outlast the saccade by 50 ms or more. **B**. Averaged activity of burst and pause type P-cells during a saccade. From Herzfeld et al. (2015), used with permission.

These puzzles are not unique to saccades. Both the diversity of simple spike responses, and the fact that modulation of discharge outlasts the movement are also features of P-cell activity during wrist (Ishikawa et al., 2014; Mano and Yamamoto, 1980; Tomatsu et al., 2016) and arm movements (Hewitt et al., 2015). For example, Ishikawa et al. (2014) recorded from nearly 200 P-cells during wrist movements and found bursters, pausers, and P-cells that exhibited both bursting and pausing. They found that the discharge modulation outlasted the movement, exhibiting no obvious patterns with respect to movement direction. Thus, given that the cerebellum is critical for precise control of saccades (Robinson et al., 1993) as well as arm movements (Becker and Person, 2019; Chen et al., 2006; Viaro et al., 2017; Vilis and Hore, 1980), it is puzzling that simple spikes of individual P-cells are modulated long after the movement has ended.

A key to this puzzle may be population coding. Peter Thier and colleagues (Catz et al., 2008; Thier et al., 2000) were the first to sum activities of all recorded P-cells into a population response. They demonstrated that the population response was a good predictor of movement duration during saccades (Thier et al. 2000), as well as kinematics during pursuit (Dash et al., 2012). We extended this approach by applying the hypothesis that the inferior olive divided the P-cells into groups wherein all the P-cells within a group shared a common response to error.

To measure the error response of each P-cell, we relied on the observation that if a saccade concluded but the target was not on the fovea, some P-cells were likely to produce a complex spike (Junker et al., 2018; Kojima et al., 2010b; Soetedjo et al., 2008; Soetedjo and Fuchs, 2006). To measure the tuning of complex spikes, we induced visual errors. Following fixation, a target was presented in the periphery. As the monkey made a saccade toward the primary target, we jumped the target to a new position (Fig. 8A). During the 50-200 ms period following completion of the primary saccade, the visual error (target position with respect to fovea) induced a complex spike with a probability that depended on the direction of the error vector (Fig. 8B). At around 150 ms after completion of the primary saccade, the animal made a corrective saccade.

**Figure 8.**
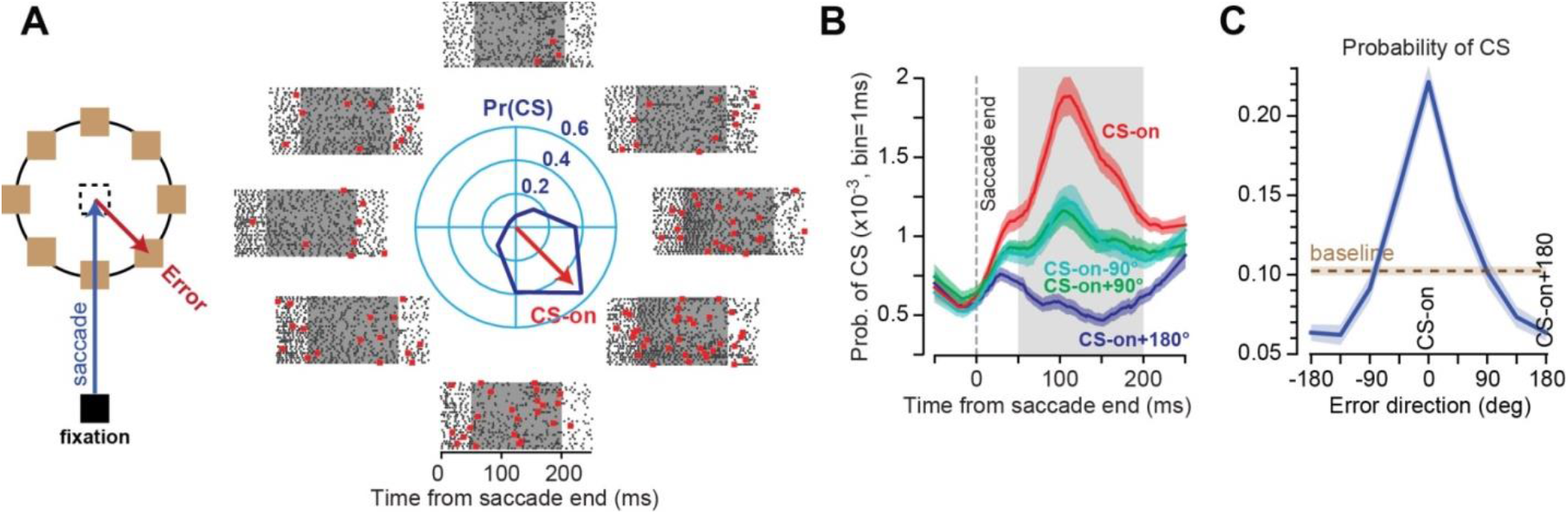
P-cells in the oculomotor vermis exhibit complex spike tuning with respect to visual error. **A**. Animals made a saccade to a target location and experienced a visual error following conclusion of the movement. The data show the simple (black dots) and complex spikes (red dots) of a P-cell in the period following saccade end. The center polar plot describes the probability of complex spike in the 50-200 ms period following saccade end. The direction of error that induced the greatest probability of complex spikes is noted by the red vector labeled CS-on. **B**. Probability of complex spike during the time following saccade end. Data are averaged from 72 P-cells. **C**. Probability of complex spike was averaged across 72 P-cells, aligned to CS-on. From Herzfeld et al. (2015), used with permission.

The jumping of the target often resulted in a complex spike, and then a corrective saccade. Thus, with these data alone we cannot say whether the complex spike was associated with the unexpected sensory event, the motor event that followed (i.e., the corrective saccade), or both. However, further experiments that we will review below suggest that the unexpected sensory event alone (Kaku et al., 2009; Soetedjo et al., 2009), without the motor correction (Tseng et al., 2007; Wallman and Fuchs, 1998), may be the main driver of the complex spikes.

Each P-cell preferred a specific error direction (Fig. 8C). The direction of error that produced the largest probability of complex spikes in the post-saccadic period for a given P-cell was labeled as CS-on. The complex spike tuning for all cells is plotted in Fig. 8C. We found that if the error was in direction CS-on, following saccade completion the probability of complex spikes peaked at around 100 ms, producing a response that was roughly twice the baseline. In contrast, if the error was in direction CS-on+180, the probability of complex spike decreased by roughly 40% below baseline. Importantly, the probability of a complex spike following saccade end was unrelated to direction of the preceding saccade. Rather, it was driven by the direction of the post-saccadic error. These results were acquired in macaques and then confirmed in marmosets (Sedaghat-Nejad et al., 2019). Thus, in two different primate species, P-cells in the oculomotor vermis exhibited a strong preference for direction of the visual error.

This preference for error appears to be a consistent feature of P-cells across species. In the cerebellum-like circuit of the electric fish, MG-cells, which are analogous to P-cells, produce broad spikes that are analogous to complex spikes. An unexpected sensory input produces broad spikes in some MG-cells, but suppresses the broad spikes in others (Muller et al., 2019). Thus, MG-cells in the electric fish also exhibit a tuning for error.

After we organized the P-cells in groups that shared similar complex spike tuning with respect to the visual error, a remarkable pattern emerged. The simple spikes produced by the population of P-cells appeared related to eye velocity (Fig. 9A). That is, whereas individual P-cells exhibited great diversity in their responses, with activity that was modulated long after the saccade had ended (Fig. 7A), as a population the simple spikes appeared to be associated with motion of the eyes rather precisely. For example, when the saccade was in the anti-preferred error direction (CS-on+180) of the P-cells, the combined simple spikes of the population increased before saccade onset, had a peak discharge that scaled with peak velocity of the eye (Fig. 9A), and then exhibited a brief period of reduce activity that coincided with deceleration of the eyes. It is remarkable that this pattern was not present in the activity of any single P-cell, but unmasked in the population, i.e., through the combined activities of the bursters, pausers, etc.

**Figure 9.**
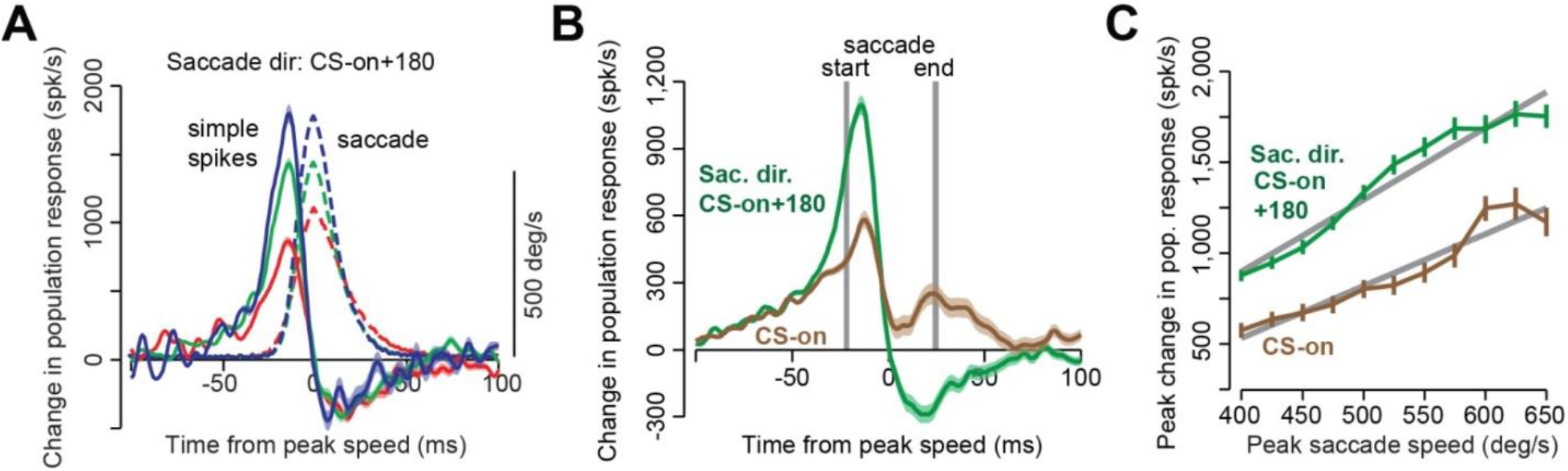
Simple spike population response of P-cells during a saccade. **A**. P-cells were organized into 50 member populations and their simple spike rates were summed as a saccade was made in direction CS-on+180. Saccades associated with three different peak velocities are shown. **B**. Simple spike population response are shown across all saccades made in direction CS-on and CS-on+180. C. Peak simple spike population response increased linearly with peak saccade velocity, but with a higher gain when the saccade was in direction CS-on+180. From Herzfeld et al. (2015), used with permission.

If saccade direction was aligned with the CS-on direction of the P-cells, the population still predicted peak eye velocity, but now with a lower gain (Fig. 9C), and without the reduced activity during the deceleration period. Therefore, the population response predicted real-time velocity of the eye, with a gain that multiplicatively depended on direction of motion. This encoding is termed a “gain-field”: the magnitude of the population response increased linearly with speed, and was cosine tuned in direction, with a multiplicative interaction between speed and direction. Gain fields are also employed by cells in the posterior parietal cortex to combine two pieces of information, position of the eye in the orbit and position of the stimulus on the fovea (Andersen et al., 1985).

In summary, in both macaques and marmosets, individual P-cells in the oculomotor vermis had simple spikes that were modulated for periods that lasted much longer than the saccade. The conjecture that the olive organized the P-cells (Fig. 6A) presented a hypothesis for population coding: P-cells that shared a similar error signal would group together to influence a DCN neuron. To find population membership, we measured each P-cell’s complex spike response to post-saccadic visual error. We assigned membership based on complex spike tuning, and then simply summed the simple spikes of the member P-cells. The resulting sum of simple spikes produced a population response that presented a consistent pattern. In direction CS-on+180, simple spikes increased before the eyes accelerated by an amount that predicted peak velocity of the eye, and then reduced activity as the eyes decelerated. In direction CS-on, simple spikes increased by a smaller amount during acceleration, predicted the peak velocity with a smaller gain, and then returned to near baseline during deceleration. Hence, when the P-cells were organized into populations that shared a similar preference for error, their simple spikes as a population appeared to play a role in controlling the real-time motion of the ongoing saccade. This framework for forming P-cells into populations await testing and confirmation in other paradigms and laboratories.

## Superior colliculus and the origin of the error signal in the complex spikes of saccade-related P-cells

Why do P-cells in the oculomotor vermis exhibit complex spikes that are tuned with respect to visual error? Recent experiments suggest that complex spike tuning reflects activity of neurons in the superior colliculus in response to events in the visual space (Kaku et al., 2009; Kojima and Soetedjo, 2018; Soetedjo et al., 2009). In this framework, activity in a specific region of the colliculus drives the contralateral inferior olive (Saint-Cyr and Courville, 1982), leading to production of complex spikes in a group of P-cells. That same collicular activity inhibits neurons on the contralateral colliculus (Mascetti and Arriagada, 1981; Munoz and Istvan, 1998; Takahashi et al., 2005), resulting in a reduction in production of complex spikes in a different group of P-cells. However, because collicular activity combines visual information from the retina with information from the basal ganglia and the cerebral cortex, the activity of the colliculus, and thus the activity that indirectly contributes to production of complex spikes, is much more than simply onset of a visual stimulus. Rather, because the superior colliculus integrates value, attention, and other variables with location of the visual stimulus, these “cognitive” modulation of the visual information may also be reflected in complex spikes.

Superior colliculus is a topographic structure, organized into regions that respond to visual inputs with respect to the fovea (Mays and Sparks, 1980). Suppose we place a target at location “a” on the left of the screen and ask a volunteer to make a saccade toward it (Fig. 10A). During this primary saccade, we erase target “a” and replace it with a target at “b”, located a few degrees to the left of “a”. As the primary saccade ends, imagine that the cerebellum predicts that the target should be near the fovea, and thus there should be activity among the foveal-related neurons in the rostral pole of the colliculus. However, the actual sensory consequence is different: the visual stimulus is a few degrees to the left of the fovea, and thus there is higher than expected activity in a group of neurons located in the right caudal superior colliculus (red region “b” in right colliculus, Fig. 10B), and lower than normal activity in a group of neurons in the left caudal superior colliculus (blue region in the left colliculus, Fig. 10B). The lower than normal collicular activity is a speculation based on the fact that caudal neurons in one colliculus inhibit the neurons on the contralateral side (Takahashi et al., 2005). Some of the affected neurons in the left and right superior colliculus project to neurons in the right and left olive, respectively (Saint-Cyr and Courville, 1982). Hence, in this hypothetical framework, the collicular map of the visual space is at least partly responsible for the complex spike tuning of the P-cells in the oculomotor vermis.

**Figure 10.**
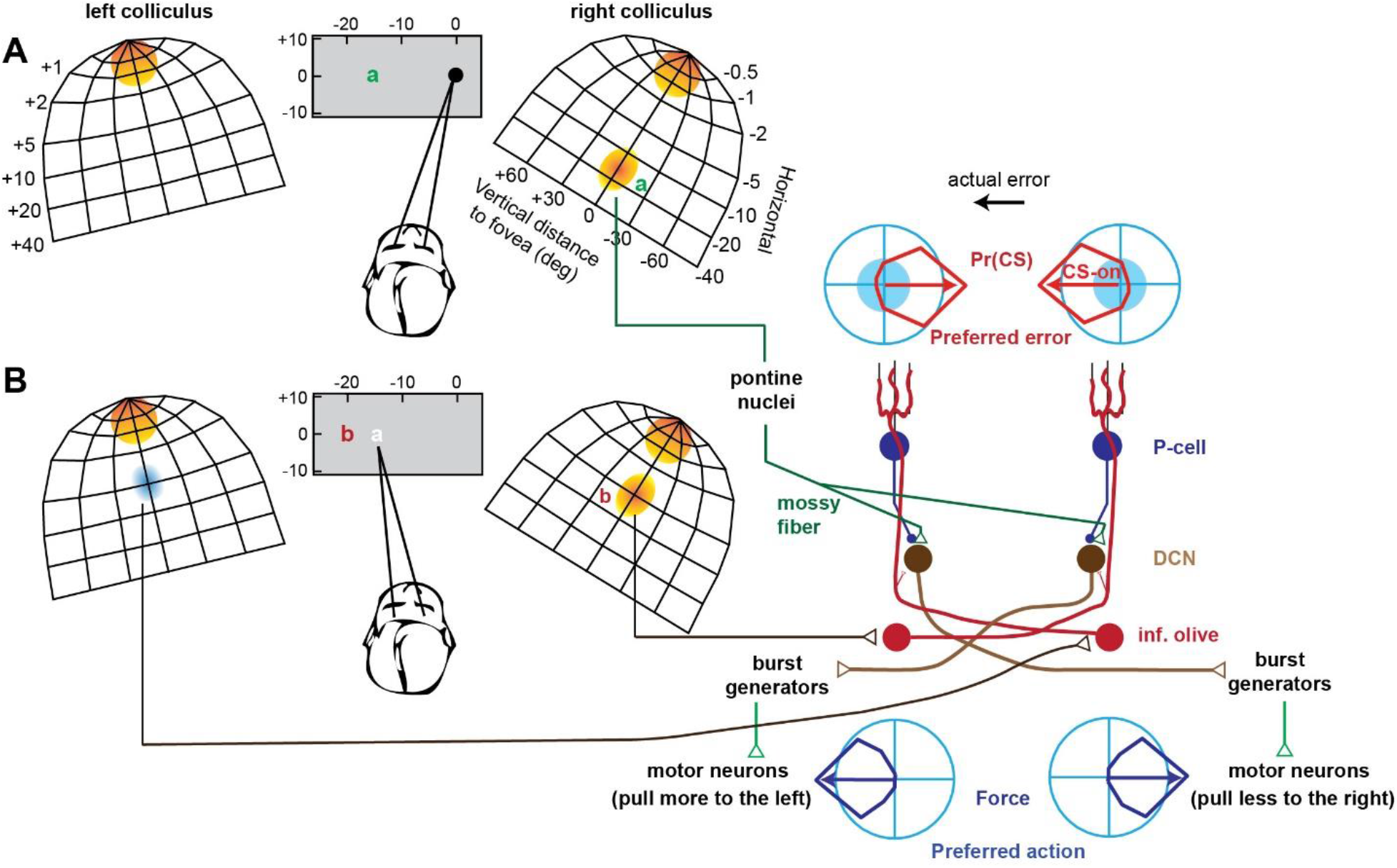
Hypothetical relationship between preferred error of a P-cell and the projections of its DCN daughters. **A**. A model of activity in the superior colliculus as the subject views a fixation point and a target is presented to the left at location “a”. During fixation, foveal related neurons in the rostral part of both colliculi are active. Presentation of the target activates neurons in the caudal region of the right colliculus (orange, labeled “a”), which results in a saccade. A copy of the saccade related activity is sent via mossy fibers to the cerebellum. **B**. At saccade end, the target is expected to be on the fovea, but is in fact at location “b”. This produces activity in a region labeled “b” in the right colliculus, which in turn inhibits a region on the contralateral colliculus. The unexpected activity on the right colliculus engages olivary neurons on the contralateral side, which produce a complex spike among P-cells on the right of the oculomotor vermis. The unexpected reduced activity on the left colliculus also affected the contralateral olive, which reduces the complex spike probability for P-cells to the left of the vermis. Presence of complex spike for the right P-cells slightly depresses the parallel fiber synapses that were activated during the preceding saccade. The lack of complex spike in the left P-cells slightly increases the weight of the parallel fiber synapse. The P-cells on the right project to DCN neurons that project to burst generators that activate motoneurons that pull the eyes to the left. On the next saccade, presentation of the same mossy fiber input now produces a slight reduction in the simple spikes of P-cells on the right (with respect to the previous trial), which results in greater force production to the left. Similarly, the P-cells on the left side of the vermis produce slightly more simple spikes, and this results in production of slightly less right-ward force. Hence, if there is a correspondence between the preferred action of the DCN neurons and the preferred error of their P-cell parents, a prediction error results in learning that correctly compensates for the experienced error. That is, the DCN neurons project to a group of downstream neurons that can produce an effect that will remedy the specific error that is of concern to the parent P-cells.

There is indirect experimental evidence in support of the idea that unexpected activity on the colliculus is the source of the complex spikes that drive learning in the saccade task. Soetedjo et al. (2009) and Kaku et al. (2009) timed sub-threshold stimulation with respect to saccade end and artificially produced activity in a small region of the colliculus. Their idea was to mimic collicular activity that would arise from a visual error and do this without eliciting the corrective saccade that would normally follow the visual error. For example, at around 80 ms following completion of the saccade, they stimulated a region of the superior colliculus that was associated with visual activity of magnitude 2° in a specific direction with respect to the fovea. They found that this sub-threshold stimulation led to saccade adaptation that corresponded to the direction and magnitude of this error vector. Their results established that unexpected activity in a small group of neurons in the superior colliculus following saccade end acted like an error, inducing learning.

To test whether unexpected collicular activity was necessary for learning, Kojima and Soetedjo (2018) deactivated a small region of the colliculus using muscimol. Their idea was to eliminate the ability of the colliculus to communicate to the inferior olive the presence of an unexpected visual event, and thus block the error signal. Notably, they took advantage of the topographic map of the colliculus and focused the disruption on a specific vector in the visual space. They found that the disruption led to near elimination of learning. That is, the animals were unable to learn from that specific error vector.

These results raise the possibility that a major source of complex spikes in the oculomotor vermis is unexpected activity in the superior colliculus. However, experiments that test for the conjecture that P-cell complex spike tuning is causally related to unexpected collicular activity have yet to be performed.

## Projections of DCN neurons and their relationship to complex spike tuning of the parent P-cells

We do not need the cerebellum to make a saccade (or any other movement). The superior colliculus is quite capable of activating the brainstem burst generators and the downstream motoneurons to produce a saccade. However, without the cerebellum’s contributions, the saccade is dysmetric. For example, if a DCN (caudal fastigial nucleus) on one side of the cerebellum is disabled, the saccade will overshoot the targets when they are shown toward the same side (hypermetria), and undershoot the targets if they are shown on the other side (hypometria) (Goffart et al., 2004; Guerrasio et al., 2010; Robinson et al., 1993). In particular, saccades to the side of the deactivated DCN are hypermetric and appear to be missing part of the deceleration commands that are needed to stop the eyes on target (Buzunov et al., 2013; Kojima et al., 2014). This is likely due to combined effects from both DCNs: the unaffected DCN produces unopposed agonist drive, while the lesioned DCN is unable to produce an antagonist drive (Bourrelly et al., 2018). Thus, the cerebellum adds something to the ongoing movement. What is it adding?

To answer this question, consider that a P-cell may have a preference for a particular aspect of error (tuning of its complex spikes), but that error is not its direct responsibility. Rather, the error arises because of a miscalculation in the output of the DCN neurons that the P-cell projects to. Where do these DCN neurons project to? Are the projections related to the preferred error (i.e., CS-on) of the parent P-cell?

Eq. (8) provides an important clue: the error signal that is conveyed via a climbing fiber to a P-cell must be associated with the olivary cell that computes the error in the output of the DCN neurons the P-cell projects to. But P-cells do not respond equally to all errors. Rather, they are most concerned with a specific error, as evidenced by their complex spike tuning. Perhaps the DCN neurons should produce downstream effects that will alleviate the specific error concerns of their parent P-cells.

In our example in Fig. 10A, we placed a target at location “a” and then jumped it to location “b”. This produced unexpected activity in certain regions of the colliculus. That activity was unexpected because it was not predicted by the GABA-ergic DCN neurons that inhibit the specific olive cells that these collicular regions project to. Hypothetically, the larger than expected activity in the right colliculus leads to increase in complex spike production among the P-cell parents of the right DCN neurons, whereas the smaller than expected activity in the left colliculus leads to suppression of complex spikes among the parents of the left DCN neurons.

Thus, a single unexpected event that occurs after completion of a saccade (the target is not on the fovea) is a miscalculation by two distinct groups of DCN neurons. Their miscalculations are conveyed to their parents P-cells via climbing fibers. In one group of P-cells, the error in the output of their DCN neurons results in an increase in their complex spike probability with respect to baseline (parents of the right DCN neurons), and in another group of P-cells (located on the left side) the same error results in a decrease in this probability with respect to baseline.

The change in complex spike probability (with respect to baseline) signals the P-cell that the DCN neurons that it projects to failed to correctly anticipate the sensory consequences of the movement. Therefore, if these DCN neurons are to correctly predict the visual consequences of this saccade, they must learn to anticipate the activity that will take place in neurons which reside in the caudal region of the superior colliculus, and cancel it when it is conveyed to the inferior olive. If they did that, they would inhibit the olive neurons that these collicular neurons project to, anticipating their activity and returning the complex spikes to baseline. As a result, if a P-cell exhibits a preference for leftward visual errors, this implies that its DCN daughters project to an olivary neuron that receives excitatory inputs from collicular neurons that encode a region to the left of the fovea (Fig. 10A).

However, it is not enough for the cerebellum to merely learn to correctly predict the sensory consequence of a movement. Rather, we need a mechanism with which to alter the motor commands and guide the saccade to the desired destination, placing the target on the fovea. That is, motor learning is not about eliminating sensory prediction errors by building more accurate predictors (termed forward models). It is about learning how to make better movements. This provides us with a clue as to where the non-GABA-ergic DCN daughters of a P-cell should project to.

**Conjecture C6.** *A DCN neuron should project to a group of neurons that can produce an effect that will remedy the specific error that is of concern to the parent P-cells*.

For example, if at saccade end the target is to the left of the fovea (Fig. 10B), and there is a P-cell population that prefers this error, then the non-GABA-ergic DCN daughters should be able to do something constructive about eliminating this error. They could do that if their axons projected to neurons that indirectly engaged muscles that pulled the eyes horizontally to the left (Fig. 10B), for example by projecting to burst generators that act on left-pulling abducens motoneurons. Similarly, for the P-cell population that prefers errors to the right, a constructive action would be for their DCN neurons to reduce the drive for right-ward pulling abducens motoneurons. With access to effectors, the non-GABA-ergic DCN neurons could profitably influence the saccade’s trajectory in a way that would help restore the complex spikes in their parent P-cells toward baseline.

Now let us consider how presence of this leftward error at saccade end should produce trial-to-trial learning. The leftward error increases probability of complex spikes for P-cells that prefer that error, slightly reducing their simple spikes on the next trial (push more to the left). The same error decreases the probability of complex spikes for P-cells for which this error is CS-on+180, slightly increasing their simple spikes on the next trial (push less to the right). As a result, on the next trial, as the target is shown at location “a” and the saccade is made, the cerebellum via its non-GABA-ergic DCN neuron adds a small leftward force to the ongoing motor commands, guiding it toward location “b”. If such an anatomy existed, the non-GABA-ergic neurons would be able to alter the saccade’s trajectory, reducing the sensory prediction error.

The suggested correspondence between the error that concerns the parents and the downstream influence of the DCN neurons helps us understand why the population of P-cells show reduced activity when the saccade is decelerating toward the CS-on+180 direction, but not when it is decelerating toward direction CS-on (Fig. 9B). In the oculomotor vermis of the cerebellum, P-cells that are on the right side of the midline project to the right caudal fastigial nucleus. These P-cells tend to prefer errors that are to the left part of the visual space (Herzfeld et al., 2015), and thus have their CS-on+180 to the right. According to conjecture C6, their DCN neurons indirectly affect effectors that produce a leftward force. When a rightward saccade is made, the P-cell population shows reduced activity during the deceleration period because this leads to increased activity in their DCN neurons, which in turn increases leftward forces, helping stop the saccade.

Is there evidence for this conjecture outside of the oculomotor system? Ekerot et al. (1995) examined the part of the interpositus nucleus that received inputs from P-cells which responded with complex spikes when the cat’s forelimb was touched or pinched on the skin. To determine the complex spike tuning of the P-cells, in the decerebrated cat they recorded local field potentials from the interpositus neurons and noted that providing a noxious pinch on a specific forelimb skin area produced an excitatory response. They used this as a proxy for the complex spike tuning of the P-cells. They then electrically stimulated these interpositus neurons and observed multi-joint movements of the forelimb. The movements were likely generated because stimulation engaged DCN neurons that projected to the red nucleus, and then neurons to the projected to the spinal cord via the rubrospinal tract.

When they compared the complex spike tuning of the P-cells (as inferred from the field potentials in the DCN) and the stimulation results, there was a general correspondence: the movement that was evoked from stimulation of a particular DCN region pulled the limb away, protecting the part of the skin which was pinched to produce the complex spike in the parent P-cells. For example, if the skin on the ventral forepaw was touched, generating potentials in a region of DCN that received input from P-cells that had a complex spike receptive field for that part of the skin, stimulation of that DCN region activated a palmar flexor of the wrist, producing a withdrawal of the forepaw away from the noxious stimulus. That is, the action produced by stimulation of DCN neurons appeared constructive in preventing the occurrence of the stimulus that produced complex spikes in the parent P-cells.

In summary, from a theoretical perspective the error that is transmitted to a P-cell must reflect the miscalculations that are made by only the DCN neurons that it projects to. P-cells may exhibit a preference for a specific error vector. To remedy this error, it would be useful if the DCN neurons produced actions that could remedy the error concerns of their parent P-cells. In the case of eye movements, this means that the DCN neurons should project to a location where they can indirectly engage motoneurons that pull the eyes in a direction that is parallel to the preferred visual error vector of their parent P-cells. That is, there should be a correspondence between the actions that can be influenced by the DCN neurons, and the errors that concern their P-cell parents.

## Effect of complex spikes on changing behavior

Conjecture C6 predicts that in the case of saccadic eye movements, P-cells that have a tuning for leftward visual errors (shown in Fig. 10 as being located on the right side of the cerebellum) should project to non-GABA-ergic DCN neurons that in turn project to neurons that through their activity produce a leftward force on the eyes. Indeed, many of the axons from the right caudal fastigial nucleus neurons cross the midline and act on excitatory and inhibitory burst neurons in the left pontomedullary reticular formation that through their projections on motoneurons produce a leftward force on the eyes (Noda et al., 1990; Quinet and Goffart, 2015, 2009; Scudder et al., 2000).

Because complex spikes are random events that may or may not occur after a given error, there is a way to test if there is a relationship between error tuning of a P-cell, and the downstream motor effects of the DCN neurons that the P-cell projects to.

Suppose in trial *n* a saccade is made and an error is observed following conclusion of that saccade. If that error is in the preferred direction of a P-cell, in some fraction of trials that P-cell will generate a complex spike, whereas in most trials it will not. Conjecture 6 predicts that if the complex spike did occur, it signaled the preferred error, which in turn produced plasticity in the P-cell and its DCN neurons. On trial *n*+1, the eyes should be pulled slightly in the direction specified by the CS-on vector of the parent P-cell.

We tested this idea in Herzfeld et al. (2018). After saccade completion, an error was present or absent, and the P-cell produced a complex spike, or not. We measured trial-to-trial change in behavior via the difference in motor output (eye velocity) from the trial in which the error was experienced to the subsequent trial in which the same target was presented.

We began with trials in which there was an error following saccade completion and the direction of that error was in the CS-on direction of the P-cell that was being recorded (Fig. 11A). In some trials, the P-cell did not produce a complex spike in the post-saccadic period (Fig. 11A, CS absent). In other trials, the P-cell did produce a complex spike (Fig. 11A, CS present). In both cases, in the subsequent trial the motor commands pulled the eyes more in the direction of error, which in this case happened to coincide with CS-on of the P-cell under study (Fig. 11A, left panel). The actual errors were similar in trials that the complex spike had or had not occurred. Yet, the change in the motor output from trial n to n+1 was significantly larger, i.e., the pull was stronger, if the P-cell produced a complex spike in the post-saccadic period of trial n (Fig. 11A, left panel, difference). A similar observation has been made in a different task (pursuit) when visual error was in direction CS-on of P-cells in the flocculus region of the cerebellum (Yang and Lisberger, 2014a).

**Figure 11.**
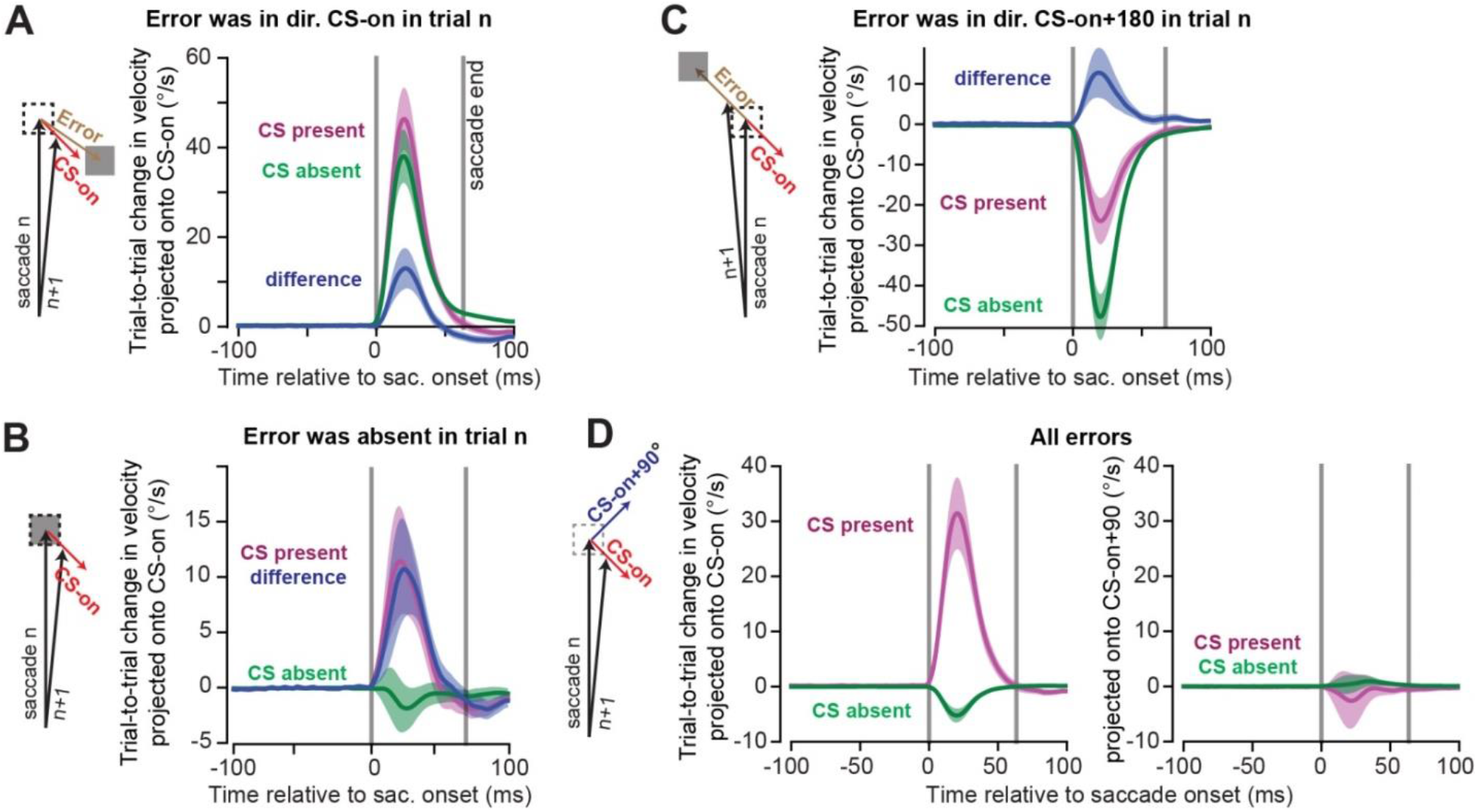
The influence of P-cells on motor output. **A.** Analysis of trials in which the error was in the CS-on direction of the P-cell. Eye trajectory (velocity vectors as a function of time) was measured during the saccade in the trial in which the error was experienced, as well as the trajectory in the subsequent trial in which saccade was made to the same target. To measure change in behavior, the trial-to-trial difference in the 2D sequence of velocity vectors was projected onto the CS-on direction of the P-cell. Following experience of an error, the next trial exhibited an increase in the velocity vector along the direction of error. However, if the P-cell produced a complex spike, velocity on the next trial was larger in direction CS-on compared to when a complex spike was absent. **B.** Same as (A) except for trials in which there was no post-saccadic error (|error| < 0.25°). Even without an error, presence of a complex spike led to increased motor output along direction CS-on of the P-cell that had produced the complex spike. **C.** Same as (A), except for trials in which errors were in CS-on+180° of the P-cell under study. If the error was opposite the preferred direction, presence of a complex spike still biased behavior in CS-on direction of that cell. **D.** Trial-to-trial change following any error. The trial-to-trial change in velocity was projected onto direction CS-on (left) and CS-on+90° (right) of the P-cell. When a CS was present, change in motor output was only in direction CS-on. From Herzfeld et al. (2018), used by permission.

Sometimes the movement was perfect: a saccade took place and the target was precisely on the fovea (Fig. 11B). Despite this lack of error, on a fraction of trials the P-cell nevertheless generated a complex spike in the post-saccadic period. Remarkably, in the subsequent trial the eyes were again pulled in the CS-on direction of that P-cell (Fig. 11B, CS present). The precise amount of change in behavior was the same whether an error was present or not (the difference curves are the same in Fig. 11A and Fig. 11B). Therefore, even without an error, the presence of a post-saccadic complex spike in a single P-cell was followed by a change in behavior in the subsequent trial. The eyes were pulled in the CS-on direction of that P-cell.

These results hinted that if a P-cell produced a post-saccadic complex spike, in the subsequent trial that P-cell and others in that population through their DCN neurons (indirectly) influenced a specific group of motoneurons, those that produced force along the P-cell’s CS-on direction. To test this hypothesis, we focused on trials in which error was in direction CS-on+180° (i.e., error was opposite the preferred error direction of the P-cell). Following experience of this error, the behavior in the subsequent trial changed in the direction of that error (Fig. 11C, left panel). Remarkably, the trial-to-trial change in behavior was significantly smaller if the P-cell had produced a complex spike. As a result, the difference between the trial-to-trial change in the motor commands that took place with and without a complex spike was always in direction CS-on, regardless of whether error was in direction CS-on (Fig. 11A), CS-on+180° (Fig. 11C), or absent altogether (Fig. 11B).

We therefore made our analysis blind to the error that the animal had experienced and instead labeled each trial based on whether the P-cell had or had not produced a post-saccadic complex spike. We found that if the P-cell produced a complex spike, the trial-to-trial change in saccade velocity was entirely in the CS-on direction of that P-cell (CS present, Fig. 11D), with no component along CS-on+90°. In contrast, if the P-cell did not produce a complex spike, the trial-to-trial change was in direction CS-on+180° of that P-cell (CS absent, Fig. 11D).

In summary, following production of a post-saccadic complex spike in a P-cell, in the subsequent trial the eyes were pulled along a vector that was parallel to the CS-on direction of that P-cell. Without a complex spike, in the subsequent trial the eyes were pushed away along that same vector. Importantly, this pattern was present even when there were no errors at the end of the saccade. This appears consist with the idea that there is a correspondence between the complex spike tuning of a P-cell, and the direction of action of the DCN neurons it projects to (Apps and Garwicz, 2005; Ekerot et al., 1995).

## Do complex spikes encode prediction errors?

The central assumption in our framework is that the inferior olive provides the error signal that serves as a teacher for the cerebellum. However, the idea that complex spikes signal a prediction error is controversial. For example, Horn et al. (2004) trained cats to reach and grasp a lever and then displaced the lever during some trials. They found no olivary neuron that discharged near the time of the error (when the paw would have grasped the lever), despite the fact that the neurons responded to taps on the paw. In addition, complex spikes are rare events that occur approximately once per second, producing a disparity between the richness of information conveyed by the simple spikes, and the poverty of the error signal transmitted by the complex spikes. Indeed, errors can double or halve in size without significant changes in the rate of complex spikes (Ke et al., 2009; Ojakangas and Ebner, 1992; Soetedjo et al., 2008).

In a saccade adaptation task, Catz et al. (2005) noted that the large errors at the onset of training produced only modest changes in complex spike probability among P-cells in the oculomotor vermis. Notably, while training produced reductions in error magnitude, the expected reductions in the probability of complex spikes did not materialize. Indeed, in tasks as diverse as arm movements, head movements, and eye movements, a consistent finding has been that error size can change without altering the probability of complex spikes (Ke et al., 2009; Keating and Thach, 1995; Soetedjo et al., 2008). From a machine learning perspective, it is quite problematic if the error signal is insensitive to error size.

Two clues shed light on this puzzle: the precise timing of the complex spikes, and the shape of their waveforms. Eccles et al. (1966) had stimulated the inferior olive and measured the complex spike response in P-cells, finding that an increase in the strength of olivary stimulation coincided with a reduction in the latency of the complex spike. This raised the possibility that complex spike timing might be an important variable with which the olive transmitted information about magnitude of error to the cerebellum. Indeed, experiments in slice preparations found that complex spike timing was a critical factor in determining the amount of plasticity at the parallel fiber to P-cell synapse (Suvrathan et al., 2016). That work demonstrated that complex spikes that arrived at a precise temporal window produced a greater effect on the simple spikes by maximizing the change in the strength of the recently active P-cell synapses.

A second clue was the shape of the complex spike waveform. Unlike simple spikes, complex spikes that are recorded near the dendrites of a P-cell have waveforms that have a variable number of spikelets (Davie et al., 2008; Monsivais et al., 2005), which in turn affects the duration of the complex spike and the subsequent rebound in simple spikes (Burroughs et al., 2017). Yang and Lisberger (2014b) measured the waveform of complex spikes from P-cells in the flocculus during a visual pursuit task. They noted that when an error occurred during pursuit (retinal slip), duration of the complex spike was longer as compared to when the animal was fixating and no error was present. Importantly, when there was an error during pursuit, the trial-to-trial amount of learning was greater when the duration of the error-induced complex spike was longer.

**Fig. 12.**
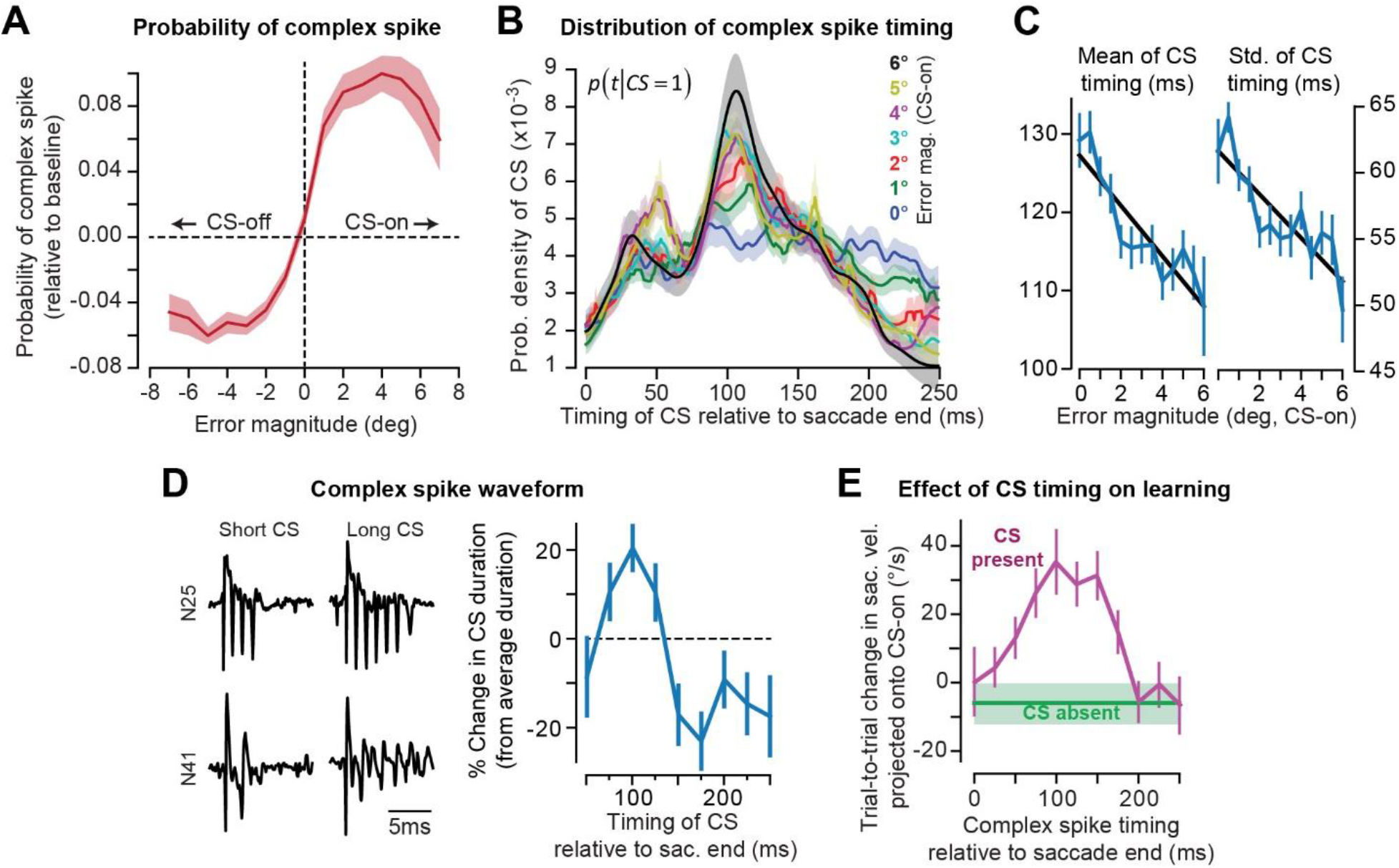
Encoding of the error vector by complex spikes. Monkeys made saccades to visual targets and experienced errors that varied in direction and amplitude (Fig. 8A). **A**. Changes in error magnitude produced little or no change in the probability of complex spikes. **B**. Timing distribution of complex spikes in the 250 ms period following completion of the primary saccade. As error magnitude increased, the timing of complex spikes exhibited less variability, becoming more focused at around 100 ms. **C**. Mean and standard deviation of complex spike timing as a function of error magnitude. As errors size increased, complex spikes occurred earlier and with less jitter in their timing. **D**. Variability in the complex spike waveform, shown for two representative P-cells. Changes in complex spike timing coincided with changes in waveform. Spikes that occurred at around 100 ms tended to have a longer duration. **E**. Effect of complex spike timing on trial-to-trial learning from error. The greatest amount of learning followed trials in which a complex spike occurred at around 100-150 ms. From Herzfeld et al. (2018), used by permission.

We recently examined timing and duration of complex spikes by systematically varying the direction and magnitude of the error vector in the task shown in Fig. 8A (Herzfeld et al. 2018). We found that the probability of complex spikes was strongly modulated by the direction of the visual error vector (Fig. 8B). However, changes in error magnitude did not produce an appreciable change in the probability of complex spikes (Fig. 12A). For example, despite the fact that error magnitude increased by 7 folds (from 1° to 7°), there were no significant changes in probability of complex spikes.

However, error magnitude affected the timing of the complex spikes. When the error magnitude was small, complex spike timing was distributed uniformly throughout the post-saccadic period (Fig. 12B). As error magnitude increased, the mean time of the complex spike shifted earlier, from 130 ms to around 100 ms with respect to saccade end (Fig. 12C, left panel). In addition, as error magnitude increased, the timing of complex spikes became more consistent, resulting in a reduction in the standard deviation of the timing distribution (Fig. 12C, right panel). As a result, small errors produced complex spikes that were distributed widely in time. In contrast, large errors produced complex spikes that clustered at a specific time: around 110ms after the onset of the error event (saccade end).

An example of variability in the waveform duration of complex spikes is shown in the left panel of Fig. 12D. A P-cell sometimes produced a complex spike that had many spikelets and a long duration, whereas in other times the same cell produced a complex spike with few spikelets and a short duration. Notably, Herzfeld et al. (2018) found that complex spikes that occurred at around 100 ms following the error event tended to be longer in duration (Fig. 12D, right panel).

These results suggested that error magnitude affected the precise timing and duration of complex spikes. If this was true, then there should be an effect of complex spike timing on trial-to-trial learning. Indeed, Herzfeld et al. (2018) noted that complex spike timing played a critical role in the process of learning: complex spikes that arrived at around 100 ms were followed by greater trial-to-trial learning than spikes that arrived earlier or later (Fig. 12E).

In summary, varying the magnitude of error does not produce an appreciable change in the probability of the complex spike. Rather, there is limited evidence for the idea that error magnitude may affect the precise timing and duration of complex spikes, which in turn regulate the amount of plasticity in the P-cell synapses. Because synchronizing complex spikes and thus coordinating their timing is also a key factor in regulating discharge of DCN neurons (Fig. 5C), the results taken together raise the likelihood that complex spike timing is an important variable with which the olive transmits error information to the cerebellum.

## Why do complex spikes carry information about the reward value of the stimulus?

The probability of a complex spike depends on many factors beyond the presence or absence of a prediction error. For example, while sudden occurrence of a stimulus by itself may not produce a complex spike in the P-cell of a naïve animal, it will do so if the animal has learned to associate that stimulus with an aversive event (Ohmae and Medina, 2015), with a greater reward (Heffley and Hall, 2019; Kostadinov et al., 2019; Larry et al., 2019), or a previous performance error (Junker et al., 2018). Withholding of reward when it was expected can also modulate complex spike rates (Heffley et al., 2018; Kostadinov et al., 2019). These results have demonstrated that the signal in the climbing fiber is not merely the difference between what was predicted and what was observed, at least not if this comparison is in terms of a simple sensory coordinate system. Rather, complex spikes also reflect a measure of the learned value of the stimulus. How does one reconcile these facts with the assumption that complex spikes encode a prediction error?

To illustrate this puzzle, consider the results of Larry et al. (2019) who trained monkeys in a pursuit task in which the amount of reward varied on each trial. Each trial began with a center target that changed color to indicate whether low or high reward would be available. As a population, P-cells showed an increase in probability of complex spikes at 200 ms following the cue that indicated high reward, and reduction of this probability when it indicated low reward. Why was the probability of complex spikes modulated by the reward value of the stimulus?

In the framework of Fig. 1, complex spikes reflect the difference between the excitatory inputs to the olive, and the inhibitory inputs from the DCN. Let us assume that in this and other visual tasks, some of the excitatory input to the olive is from the superior colliculus (Saint-Cyr and Courville, 1982). Thus, complex spikes would be generated to signal unexpected activity among a group of neurons in the superior colliculus. However, activities of collicular neurons are not simply a reflection of retinal input, but also inputs from the basal ganglia and the cerebral cortex. For example, response of neurons in the superior colliculus is greater if the stimulus is novel and threatening as compared to familiar and unthreatening (Lee et al., 2020). The response is greater if the stimulus is associated with reward (Ikeda and Hikosaka, 2003). This utility-dependent collicular response to the visual stimulus is likely because of value-dependent modulation of inhibition from the basal ganglia (Sato and Hikosaka, 2002) and excitation from the frontal eye fields (Glaser et al., 2016).

Upon presentation of a visual stimulus, the basal ganglia and the frontal eye fields evaluate the stimulus and alter the inhibition and excitation that they respectively impose on the superior colliculus (Glaser et al., 2016; Sato and Hikosaka, 2002). As a result, the collicular response to a visual stimulus depends on the location of the stimulus on the retina, as well as the utility that the cortex and the basal ganglia assign to that stimulus. This leads to the speculation that utility dependent activity on the colliculus may in turn be at least partly responsible for modulation of complex spike probability as a function of the reward value of the stimulus.

In summary, several recent reports have shown that complex spikes carry information about the reward value associated with a stimulus (Heffley and Hall, 2019; Kostadinov et al., 2019; Larry et al., 2019). To reconcile this fact with the idea that complex spikes carry an error signal, we can consider an olivary neuron that receives two inputs: inhibition from a DCN neuron, and excitation from a neuron in the superior colliculus. The DCN neuron produces an output that attempts to predict and cancel (at the level of the olive) the activity in a specific region of the superior colliculus. However, the collicular activity is not simply the location of the stimulus in the visual field, but also a reflection of its utility as evaluated by the frontal eye fields and the basal ganglia. Thus, an error in the activity of a DCN neuron would be reflected in complex spikes that carry information about both the location of the stimulus, and its utility.

## Implication: multiple timescales of adaptation

Let us try to broaden our scope and ask whether the conjecture regarding population coding might provide clues as to why behavior during learning exhibits certain remarkable features, namely multiple timescales, resistance to erasure, and spontaneous recovery.

We noted that presence of a complex spike in a P-cell was followed by a change in behavior, but the absence of a complex spike was also followed by a change, one that was much smaller and in the opposite direction (Fig. 11D). This is because both the presence and absence of a complex spike leads to P-cell plasticity, but in different directions, and with different rates. For example, Yang and Lisberger (2014a) trained monkeys to pursue a visual target and induced an unexpected change in the motion of the target. Sometimes the error was in direction CS-on for the P-cell, and sometimes it was in direction CS-on+180. If the error took place and a complex spike occurred, on the next trial the simple spike rate during pursuit exhibited a decrease (Fig. 13A, pursuit). If the complex spike did not occur, the simple spike rate during the subsequent trial exhibited a small increase. We observed roughly the same pattern during saccades (Herzfeld et al., 2018): if at saccade completion a complex spike was present, on the next trial the P-cell showed a reduced rate of simple spikes (Fig. 13A, saccade). (The effect of complex spike absence on the simple spikes was less clear in the saccade data.)

**Figure 13.**
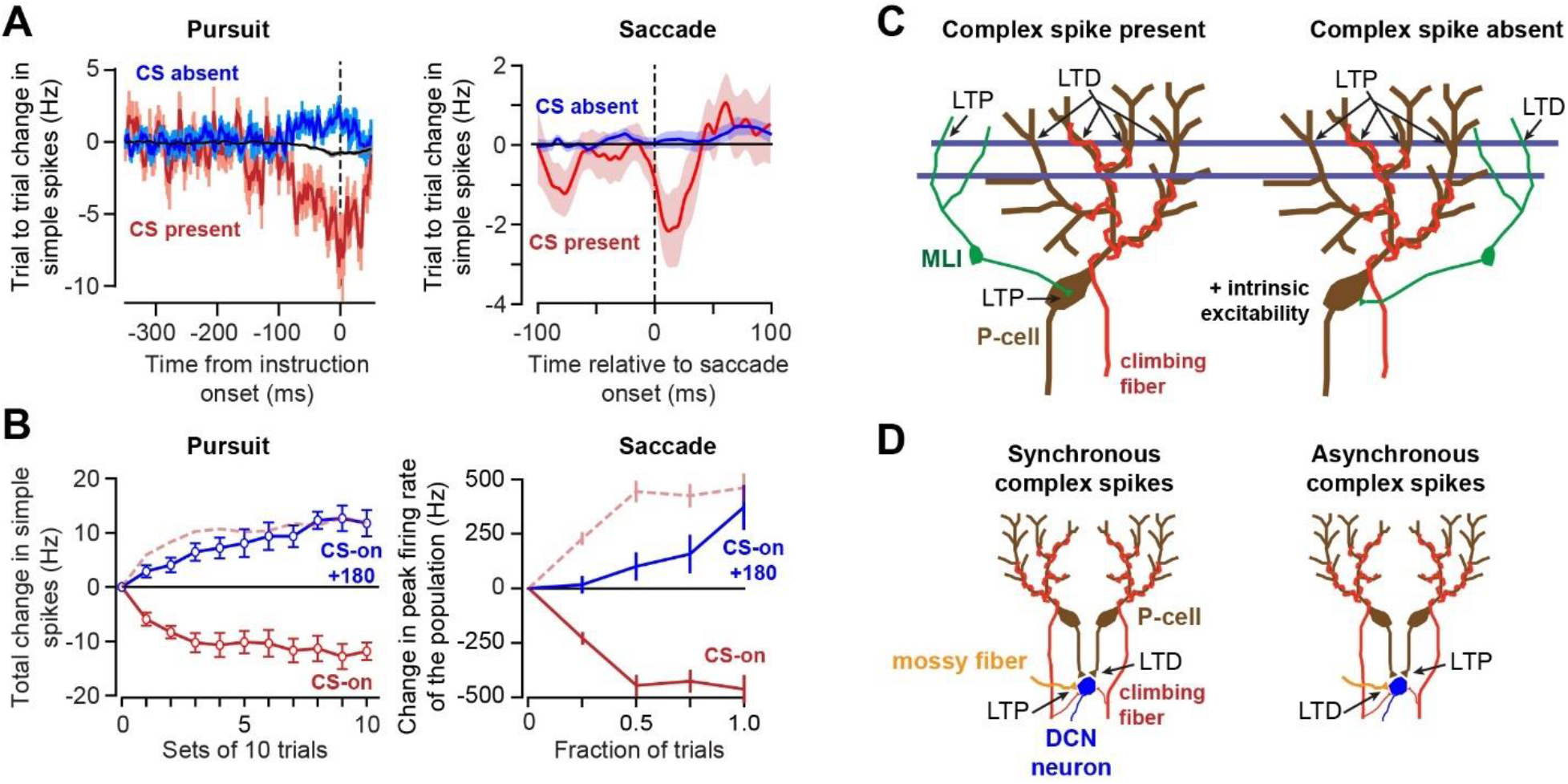
Change in simple spikes following presence or absence of a complex spike. Data are from pursuit and saccade tasks. **A**. Trial-to-trial change in simple spikes. In the pursuit task, the inter-trial interval was 2.5 sec. Time zero indicates the time during pursuit in which an unexpected change in the visual stimulus occurred. The black trace is the baseline relationship during pursuit. In the saccade task, change in simple spikes are plotted with respect to saccade onset. **B**. Total amount of change in simple spikes when the errors are in CS-on or CS-on+180 direction. For the pursuit task, the change is the expected value for a single P-cell, averaged across all P-cells. For the saccade task, the change is the sum for 50 P-cells that make an average population. The dashed lines are the CS-on data mirrored along the x-axis. It appears that the rate of change in simple spikes is faster when errors are in direction CS-on. **C**. Presence of climbing fiber activity (with parallel fiber activity) results in long-term depression (LTD) at the parallel fiber to P-cell synapse, long-term potentiation (LTP) at the parallel fiber to P-cell synapse, and rebound potentiation at the molecular layer interneuron (MLI) to P-cell synapse. When climbing fiber activity is absent, parallel fiber to P-cell synapses undergo LTP, LTD is induced at the parallel fiber synapse onto MLI neurons, and the intrinsic excitability of the P-cell is increased. Parts A and B are from Yang and Lisberger (2014), and Herzfeld et al. (2018), used by permission.

These changes in simple spike response of P-cells were likely due to the influence that climbing fiber activity has on induction of plasticity in the parallel fiber synapses on P-cells, as well as parallel fiber synapses on molecular layer interneurons (Fig. 13C, for a comprehensive review, see (Gao et al., 2012)). Parallel fiber synapses on P-cells undergo post-synaptic LTD following paired stimulation of parallel fiber and climbing fibers (Hansel et al., 2006; Jörntell and Ekerot, 2002). In contrast, post-synaptic LTP takes place when the parallel fibers are stimulated without climbing fiber activity (Coesmans et al., 2004). Thus, direction of plasticity at the parallel fiber P-cell synapse depends on presence or absence of complex spikes. In addition, stimulation of parallel fibers without climbing fiber stimulation can increase the intrinsic excitability of P-cell dendrites, reducing the simple spike pause duration that follows production of a complex spike (Grasselli et al., 2020, 2016).

Climbing fiber activity also influences the strength of the parallel fiber synapses onto molecular layer interneurons. These synapses undergo post-synaptic LTP when parallel fiber stimulation is coincident with depolarization of stellate cells, which is thought to be naturally occurring through spillover from activity in climbing fibers (Szapiro and Barbour, 2007). In contrast, post-synaptic LTD takes place in these synapses in response to low frequency parallel fiber activity without climbing fiber stimulation (Piochon et al., 2010).

In summary, presence of a complex spike is likely to reduce P-cell activity via induction of LTD at the parallel fiber P-cell synapse, and LTP at the parallel fiber molecular layer interneuron synapse. Absence of a complex spike is likely to increase P-cell activity via induction of LTP at the parallel fiber P-cell synapse, LTD at parallel fiber molecular layer interneuron synapse, and increase in the intrinsic excitability of the P-cell.

Yang and Lisberger (2014a) measured simple spike changes in P-cells over the course of 100 trials. They found that if the error was consistently in the CS-on direction of the P-cell, the simple spike rate during the pursuit declined steadily (Fig. 13B, pursuit). If the error was consistently in the CS-on+180 direction, the simple spike rate increased. Notably, the rate of increase in the simple spikes was slower than the rate of decrease during pursuit. We observed a similar pattern during saccades (Herzfeld et al., 2018): the simple spikes produced by the population of P-cells decreased for CS-on errors, and increased for CS-on+180 errors. However, the decrease occurred faster than the increase (Fig. 13B, saccade).

The differing sensitivities of simple spikes to presence or absence of a complex spike is an important feature because it suggests that a single error engages mechanisms that naturally exhibit multiple timescales, some fast (CS-on), others slow (CS-on+180). Let us show how this fact can contribute to the multiple timescales of behavior during adaptation.

Suppose we have two P-cell populations, one with a complex spike tuning to the left, and the other to the right (as in Fig. 10). Imagine that these P-cells are distinct populations that projects to two different DCN neurons. On trial *n* we make a saccade and during that saccade population 1 produces simple spike response 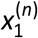. On the same trial population 2 produces response 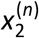. Following saccade completion we experience a post-saccadic error *e*^(*n*)^: the target is to the left of our fovea. This error is in direction CS-on for population 1, and CS-on+180 for population 2. The error produces plasticity in our two populations (and perhaps in their DCN daughters). On the next trial, the responses in the two populations are as follows:

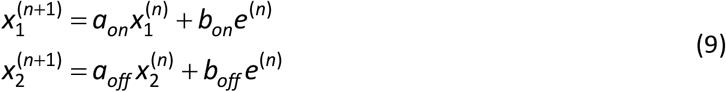

The term *b_on_* refers to the trial-to-trial change in the simple spike response when the error is in direction CS-on. The term *a_on_* refers to the trial-to-trial change due to passage of time between the two trials (decay toward baseline).

Eq. (9) has parameters that can be inferred from the available data. Following experience of error *e*^(*n*)^, the saccade is corrected by a small amount *q* on the subsequent trial. The adaptive response *q* is due to the contributions of these two populations. Suppose that if a population produced a complex spike in response to the error, it affected behavior on the next trial in the direction of its CS-on by amount *y*_1_. If it did not produce a complex spike, it affected behavior in the direction CS-on+180 by amount *y*_0_. Fig. 8C provides the probabilities associated with production of a complex spike. From these probabilities, we form the following two equations:

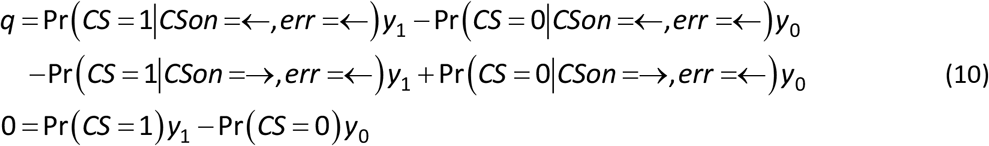

The first equation provides the probabilistic contributions of the two populations to learning from error. For example, the term Pr(*CS* = 1|*CSon* =←,*err* =←) specifies the probability of a complex spike in population 1, given that the CS-on direction is leftward, and error direction is leftward as well. The term Pr(*CS* = 1 *CSon* =→,*err* =←) specifies the probability of a complex spike in population 2, given that the CS-on direction is rightward, but the error direction is leftward. The values for each probability term can be acquired from Fig. 8C. The second equation provides the constraint associated with homeostatic regulation; sum total of plasticity, weighted by the overall probability of complex spikes, is zero.

The solution to Eq. (10) is *y*_1_ = 5.6*q* and *y*_0_ = 0.6*q*. This means that following an error, the population with a CS-on is responsible for 79% of the adaptive response, whereas the population with CS-on+180 is responsible for 21%. Therefore, given the leftward error, both populations learn from error, one by increasing the simple spike rate, and the other by decreasing it. However, because population 1 prefers the error, it will contribute roughly 3.5 times more to the adaptive response than population 2.

The next critical feature of the P-cell’s response to error is that between the trial in which the error is experienced and the subsequent trial, the error induced adaptation decays with passage of time. Yang and Lisberger (2014a) found that the trial-to-trial change in a P-cell’s adaptive response depended not only on production of a complex spike, but also on the passage of time between trials. The passage of time produced decay in the error induced change in the simple spike response.

We do not know whether the plasticity induced by production of a complex spike decays differently than the plasticity induced by lack of a complex spike. But we can make a guess by examining the data in Fig. 13B. After a long series of trials, the change in simple spikes reaches an asymptote. Remarkably, this asymptote is not different for CS-on errors that encourage production of complex spikes, and CS-on+180 errors that decrease it. However, the trial-to-trial rate of change is faster when the errors are in direction CS-on.

A constant asymptote but differing rates of convergence require that trial-by-trial amount of forgetting be larger for the process that learns more rapidly (Albert and Shadmehr, 2018; van der Kooij et al., 2015; Vaswani et al., 2015; Vaswani and Shadmehr, 2013). To see this, consider that a process that learns from error *e* by amount *b* but suffers from forgetting between trials by amount *a* (as in Eq. 9), exhibits a steady state *x*^(*ss*)^ that at asymptote is defined by:

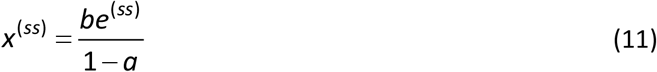

Eq. (10) and the data in Fig. 13B imply that when the error is in direction CS-on, the P-cell population that prefers this error will exhibit an error sensitivity that is larger than the population for which this error is in direction CS-on+180. That is, *b_on_* > *b_off_*. The data in Fig. 12B implies that 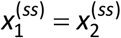. Hence, we infer that *a_on_* < *a_off_* and *b_on_* > *b_off_*.

This means that when an error is aligned with CS-on, sensitivity to that error in the P-cells and their downstream DCN neurons is greater than when the same error is aligned with CS-on+180. However, if the error is aligned to CS-on, the resulting adaptation decays at a faster rate as time passes from one trial to the next, as compared to when error is aligned to CS-on+180. Thus, given a single error, one population of P-cells learn a great deal (CS-on), but also forgets at a faster pace, whereas another population of P-cells learns less (CS-on+180), but retains a greater amount.

Behavior during learning of tasks that are believed to depend on the cerebellum has often hinted that the change in the motor output appears to be a sum of two or more adaptive states: some that learn slow but retain what they learned, and others that learn rapidly but are more susceptible to forgetting (Kording et al., 2007). It is curious that P-cell populations with CS-on and CS-on+180 error preference exhibit some of the properties of these theoretical states. Whether error-dependent adaptation in these two populations plays a causal role in the multiple timescales of behavior remains an open question.

## Implication: resistance to erasure and spontaneous recovery of memory

In numerous paradigms, from fear conditioning to motor adaptation, learning exhibits a remarkable property: acquisition of a novel behavior followed by its extinction does not erase the acquired memory. Rather, following passage of time, behavior reverts toward the originally learned pattern. That is, unlike an artificial neural network, in many biological systems one cannot unlearn a behavior by reversing the sign of the error vector (Pekny et al., 2011). Our framework provides clues as to what may be the cause of this ubiquitous feature of learning.

In the field of classical conditioning, bees can learn to associate an odor with nectar, extending their proboscis upon presentation of the odor (Stollhoff et al., 2005). They will extinguish this response if the odor is presented without the nectar. However, following passage of time, the bees once again extend their proboscis when they are presented with the odor. In the field of fear conditioning, a stimulus can be associated with a shock, inducing fear. This fear can be extinguished if the stimulus is presented without the shock (Schiller et al., 2010). However, fear of the stimulus returns following passage of time. In the field of motor learning, people and other animals respond to a perturbation by modifying their motor commands through learning from error. If the perturbation changes direction, reversing the error vector, behavior returns to baseline. However, with passage of time the behavior spontaneously reverts back and they once again produce the motor commands that they had originally learned (Criscimagna-Hemminger and Shadmehr, 2008; Kojima et al., 2004; Sarwary et al., 2018; Smith et al., 2006).

The central theme in these experiments is resistance to erasure, leading to spontaneous recovery. A mathematical model (Kording et al., 2007; Smith et al., 2006) had suggested that during learning, changes in behavior may be supported by two adaptive processes: a fast adaptive process that has high sensitivity to error along with poor retention, and a slow adaptive process that has poor sensitivity to error along with robust retention. In this model, learning is due to changes in both the fast and the slow processes. When error reverses direction, behavior returns to baseline, but the memory is not erased. Rather, the slow process retains what it had learned, but that learning is masked by learning in the fast process. With passage of time, the fast process decays, inducing spontaneous recovery of the previously learned behavior.

In our model of the cerebellum, training starts with errors that are in a constant direction, thus producing rapid changes in a specific P-cell population with CS-on aligned with that error, and slower change in other populations. During extinction training, the error reverses direction. This reversal of the error vector now produces complex spikes in a different population of P-cells. That is, because of preference for direction of error, training engages primarily one olivocerebellar module, whereas extinction is likely to engage another olivocerebellar module. The populations that had learned slowly now learn rapidly because the error is aligned with their CS-on. In contrast, the populations that had learned rapidly now learn slowly because the error is aligned with their CS-on+180. Critically, reversal of error direction produces adaptation that is approximately 3.5 times slower in the CS-on+180 direction as compared to CS-on. Thus, reversal of error cannot easily erase the history of past trials.

In summary, in this framework behavior exhibits apparent extinction not because P-cells have returned their activities to their baseline state, but because of a balance that has been reached between the changes in the CS-on and CS-on+180 populations of P-cells. With passage of time, the previously rapidly adapting population exhibits decay toward baseline, resulting in the spontaneous recovery of behavior. Thus, a consequence of preference for error in the cerebellum may be that behavior is more easily learned than unlearned: unlearning requires 4 or more times the amount of training than the amount that it took to acquire the original behavior. However, this potential link between spontaneous recovery and the rates of decay in the error-induced adaptation of the CS-on and CS-on+180 populations of P-cells is still a conjecture, waiting to be tested.

## Discussion

Activity of single neurons in the cerebral cortex and the cerebellum are often difficult to interpret as a function of the ongoing behavior. In the cerebral cortex, the current approach is to pool the neurons into pseudo-populations in which a weight is associated with each neuron’s response based on a statistical measure of its activity (e.g., principal components), and the weighted activities are summed to produce a population response. In the cerebellum, however, P-cells organize into small groups that converge on a single DCN neuron (Person and Raman, 2012a). Is there a principled way to understand population coding in the cerebellum?

Here, we viewed the cerebellum as a three-layer network in which simple spikes of P-cells carried information from the middle layer to the output layer. We made the assumption that the inferior olive compared the cerebellar output with a desired one, and thus produced an error sign. Using ideas from machine learning, we derived how this error signal must guide learning in the DCN neurons that produced the erroneous output, and the P-cells that projected to them. The theory highlighted a major problem: while olivary inputs to the P-cells are remarkably strong, the same inputs to the DCN neurons are quite weak (Lu et al., 2016). This disparity made it unclear how the synapses on the output layer neurons could change based on the signal in the olivary projections.

The anatomy suggested the error associated with a DCN neuron’s output was computed by an olive cell and then sent via climbing fibers to a few P-cells which then projected back to that specific DCN neuron (Chaumont et al., 2013). Perhaps the error in the output of a DCN neuron was conveyed to it indirectly through the complex spike that was produced by its parent P-cells (Heck et al., 2013). However, a complex spike is effective in influencing a DCN neuron’s activity only if it is synchronized with complex spikes that are generated in other P-cells in the same population (Tang et al. 2019). While this observation remains to be confirmed in other preparations, it suggests that the error in the output of a DCN neuron may be communicated to it via synchronous complex spikes in the parent P-cells. This requirement for complex spike synchrony implied that the parent P-cells of a DCN neuron could not be selected randomly. Rather, the olive must select the P-cells that form a population so that they share the same preference for error.

Thus, the idea is that the role of olivary input may be two folds: 1) provide complex spikes that act as a teaching signal that alters simple spike patterns, and 2) through complex spike synchrony among a small group of “parent P-cells” provide a teaching signal for the DCN neuron that is responsible for generating the erroneous output.

This framework made two predictions. The population of P-cells that projected onto a nucleus neuron was likely composed of those P-cells that received climbing fibers from the olivary neurons that also projected to that nucleus neuron (Fig. 6A). Furthermore, efficient learning in a P-cell required that its single climbing fiber should come from a specific olivary neuron: the olivary neuron that computed the errors associated with the DCN neurons that this specific P-cell projected to (Fig. 6B).

Many aspects of this framework remain untested. Yet, in one respect the framework has proven useful: decoding simple spike activity of P-cells as a population during voluntary movements (Herzfeld et al., 2018, 2015). During a saccade, individual P-cells exhibited simple spikes that burst, paused, or did both, with a modulation that outlasted the movement by 50 ms or more (Kojima et al., 2010b; Thier et al., 2000). However, when the P-cells were organized into populations based on their complex spike response to error, the sum total of simple spikes of the member P-cells presented a clear pattern: in direction CS-on+180, simple spikes increased before the eyes accelerated and then decreased as the eyes decelerated. In direction CS-on, simple spikes increased by a smaller amount during acceleration but returned to near baseline during deceleration. Thus, population response of P-cells during saccades appeared related to the acceleration needed to move the eyes, and the deceleration needed to stop it. These patterns were not present in any single P-cell, but revealed when P-cells were grouped based on their preference for error.

Because P-cells are not the output of the cerebellum, they do not directly control any aspect of behavior. To understand their contributions to behavior, it is useful to ask where the DCN neurons of a given population of P-cells project to. Trial-by-trial analysis of complex spikes (Herzfeld et al., 2018; Yang and Lisberger, 2014a) has provided evidence for the conjecture that there is a correspondence between the complex spike tuning of a P-cell, and the direction of action of the DCN neuron it projects to (Apps and Garwicz, 2005; Ekerot et al., 1995). It appears that the direction of error that a P-cell is concerned with (CS-on) is aligned with the direction of action of its DCN neuron. Thus, we have a simple heuristic: the output if a DCN neuron influences actions that are aligned with the specific errors that its parent P-cell is concerned with.

There may be a consequence of this anatomy on how the cerebellum learns from error. During the past two decades, numerous paradigms have quantified behavior during motor adaptation and noted a few consistencies. First, during a single session, errors decline as training proceeds, but there are at least two rates of learning, one fast, and the other slow. Second, if the errors reverse direction, thus causing “unlearning”, behavior returns to baseline (termed extinction training). However, reversal of error does not erase the memory, as evidenced by the fact that with passage of time, behavior spontaneously reverts back toward the initially learned behavior.

It is possible that the multiple timescales of learning arise from the fact that a single error engages multiple P-cell populations, some that produce a complex spike in response to that error, and others that suppress this production. Both induce plasticity that affects behavior, but the effects are asymmetric: simple spike rates change more after experience of a complex spike (as compared to when the complex spike is missing), and may also decay more with passage of time. In addition, the conjecture that the P-cells act as substitute teachers for their DCN neurons (Medina and Mauk, 1999) raises the possibility that even slower timescales of adaptation arise from plasticity in the DCN neurons (Herzfeld et al., 2020), though this conjecture also remains to be tested. The asymmetric response of a P-cell to presence or absence of a complex spike may be responsible for the fact that reversal of error during extinction training does not reverse the effects of past learning. Instead, it produces spontaneous recovery with passage of time.

To be sure, there are numerous limitations in our framework. For example, our central conjecture that complex spikes signal a prediction error is not without problems. Some complex spikes occur at the onset of salient events that by themselves have no obvious interpretation as an error (Welsh et al., 1995). A further complexity is that a single P-cell can produce complex spikes in response to sudden onset of many types of sensory inputs. For example, Ju et al. (2019) used 2-photon calcium imaging to measure complex spike driven responses in crus 1 region of awake mice and found that a response was elicited following air puff stimulation of the whiskers, tactile stimulation of the cheeks and lips, sound of the device that produced the air puff, and flash of an LED. If complex spikes represent a prediction error, they seem to lack specificity.

Another problem is that complex spikes, rare events that occur approximately once per second, appear poorly matched to reflect errors of DCN neurons, neurons which fire at rates of tens of spikes per second. The key appears to be complex spike timing. Small errors produce a complex spike that has high temporal variability, while large errors produce a complex spike that has low temporal jitter (Herzfeld et al., 2018; Najafi et al., 2014), synchronized with other P-cells (Najafi et al., 2014). The timing (Herzfeld et al., 2018; Suvrathan et al., 2016) and the number of spikelets (Rasmussen et al., 2013; Yang and Lisberger, 2014b) affect not only the amount of plasticity at the P-cell, but also the strength of suppression in the downstream DCN neuron, thus potentially affecting its plasticity.

## A potential problem with conveying error information via synchronous complex spikes

Our conjecture that synchronous complex spikes among a population of P-cells may be a method with which to transmit error information from the olive to a DCN neuron introduces a new problem: what happens if there is synchrony among the simple spikes of the parent P-cells? Indeed, we noted that optogenetic pulsed stimulation of P-cells induced simple spike synchrony that resulted in DCN plasticity (Lee et al., 2015) as well as behavioral learning (Nguyen-Vu et al., 2013).

On the one hand, the olive might rely on P-cells to serve as a substitute teacher for the DCN neuron that they converge upon, indirectly providing it with error information via its climbing fiber. On the other hand, the rest of the brain, via mossy fibers and the granule cells, rely on the P-cells to provide the DCN with a major source of information about the state of the body, thus sculpting the output of the DCN neurons. Hence, whereas most of the brain relies on the P-cells to provide simple spikes that influence the DCN’s output, the olive relies on the P-cells to provide spikes that teach the DCN about the errors of the same outputs. How can the P-cell serve both masters?

From a learning perspective, in order to disambiguate the complex spike transmitted error signal from other signals that the P-cell transmits to the DCN there is a need for suppressing synchrony among the simple spikes. In this way, if a synchronous event is detected at the DCN neuron, it can only be due to the error signal from the olive. However, from a control perspective, if the mossy fiber input via granule cells to the P-cells are to influence the output of a DCN neuron, then there is a need for synchrony among the simple spikes (Person and Raman, 2012b).

Let us consider the control perspective. P-cells fire at a baseline rate of around 50 Hz. In a decerebrated cat this background activity produces little change in the membrane potential of the downstream DCN neuron (Bengtsson et al., 2011). Through synchrony, simple spikes of P-cells can drive the activity of a DCN neuron (Person and Raman, 2012a). For example, if the simple spikes are asynchronous, the nucleus cell is inhibited, but if the simple spikes are briefly synchronous, at 8 ms latency the DCN neuron can experience a rebound that produces a spike. Thus, through synchronization of simple spikes the downstream DCN neuron can relay the specific timing of the P-cell activity to neurons outside of the cerebellum, thus providing a pathway to influence behavior.

Indeed in both anesthetized and behaving animals, nearby P-cells have cross-correlations that exhibit a level of synchrony among complex spikes as well as simple spikes (De Zeeuw et al., 1997; Ebner & Bloedel, 1981; Heck et al., 2007; Sedaghat-Nejad et al., 2019; Shin & De, 2006; Wise et al., 2010). For example, Heck et al. (2007) found that in rats, P-cells within a few hundred microns fired synchronously during the period of reach to grasp, but not during the period after grasp completion. Hong et al. (2016) noted evidence for simple spike synchrony around saccade onset. Simple spike synchrony exists even when chemical synapses in the cerebellar cortex are inactivated (Han et al., 2018), raising the possibility that some neighboring P-cells are electrically coupled.

However, it is unclear whether in the intact animal, control of DCN output relies on synchrony of simple spikes. Experimental results and simulations of Payne et al. (2019) suggest that whereas the mean rate of simple spikes is critical for control of eye velocity during a pursuit task, the timing of individual spikes plays a less significant role. Irregular simple spike timing of a population of P-cells, though produced synchronously through optogenetic stimulation, did not result in eye velocities that were different than expected from the averaged firing rates of the P-cells. If synchronous simple spikes are present during pursuit eye movements, they do not appear to provide an advantage for control of DCN output.

What about the idea that unambiguous transmission of error information via synchronous complex spikes might require a mechanism to suppress simple spike synchrony? Intriguingly, in the cerebellar cortex, one function of interneurons may be to reduce the synchrony of simple spikes. P-cells receive excitatory inputs from granule cells, and inhibitory inputs from small interneurons in the molecular layer, as well as Golgi cells. Hausser and Clark (1997) measured activity in P-cells and molecular layer interneurons in cerebellar slices, thus removing the influence of excitatory inputs. Without their external excitation, P-cells and the interneurons nevertheless produced spontaneous activity. Critically, the spontaneous activity of individual P-cells was highly irregular: inter-spike intervals (ISI) exhibited a long tail, indicating that the coefficient of variation of ISI was large. To measure the effect of the inhibitory interneurons on the P-cells, Hausser and Clark (1997) blocked inhibitory synapses and found that mean firing rates increased in both P-cell and the interneurons. Removal of inhibition transformed the ISI histogram into a pattern that exhibited a single narrow peak without a tail. That is, removal of inhibitory inputs to the P-cells increased their regularity.

Do the inhibitory interneurons play a role in reducing simple spike synchrony in the intact animal? Brown et al. (2019) made mutant mice that lacked stellate and basket cells, the principal interneurons in the cerebellar cortex. They recorded P-cell activity in the awake mice and found that despite loss of stellate cells, there was little or no change in the mean firing rate of P-cells. Rather, the mutation affected the variability of simple spikes: without the stellate cells, the P-cell simple spike ISI histogram had a reduced tail. This argument is admittedly weak because the pattern of irregularity may be shared among P-cells, and thus irregularity by itself is not an indicator of reduced synchrony. Regardless, the available results raise the possibility that a function of the interneurons in the cerebellar cortex may be to reduce simple spike synchrony.

We might imagine that by reducing simple spike synchrony, the molecular layer interneurons allow synchronous complex spikes to reliably convey error information to the DCN. If so, then removal of this inhibition might prevent normal learning in the DCN. Results of Wulff et al. (2009) are consistent with this. The authors produced mice in which P-cells lacked GABA receptors and observed an increase in the regularity of the simple spikes. These mice learned a gain-down VOR task normally on Day 1, but when tested on Day 2, they showed a deficit in the ability to retain what they had learned.

In summary, in the framework that we have described, the P-cells face a conflict in the two roles that they are asked to play. On the one hand, given the objective of transmitting error information from the middle layer (P-cells) to the output layer (DCN neurons), we need to encourage complex spike synchrony but avoid simple spike synchrony among the P-cells that converge upon a single DCN neuron. On the other hand, given the objective of driving activity of the output layer neurons, it is useful to encourage simple spike synchrony. Indeed, a function of the inhibitory interneurons in the molecular layer may be to transform the spontaneous simple spike activity of P-cells into an irregular pattern, which in turn may be required for normal learning in the DCN. However, synchrony of simple spikes has the advantage of allowing the P-cells to drive the activity of DCN neurons, thus affecting behavior.

Whether simple spikes are synchronous among the P-cells that form a population (as predicted by the control hypothesis), or asynchronous (as predicted by the error signal hypothesis), remains unknown.

## Other limitations

Despite our focus on the theoretical problem of learning in the DCN neurons, we did not consider the question of what the cerebellum’s output is specifying. For example, we focused on P-cell activity during saccades, which arguable is the one voluntary movement for which there is excellent information regarding the role of the various non-cerebellar structures (superior colliculus, brainstem burst generators, etc.). Yet, we did not discuss the activity of neurons in the DCN during saccades. There are two reasons for this. First, activity of individual DCN neurons are difficult to interpret. In the caudal fastigial nucleus, neurons burst (Fuchs et al., 1993; Helmchen et al., 1994; Sun et al., 2016), pause (Kleine et al., 2003), or do both, with a timing that may differ as a function of direction and speed of saccades. A second reason is that there is significant anatomical diversity in the projections of fastigial neurons: some project to the brainstem burst generators (Noda et al., 1990), while others project to the rostral region of the superior colliculus (May et al., 1990). This may account for the fact that disruption of caudal fastigial nucleus not only affects control of saccades, it also affects the equilibrium position of the eyes during fixation (Goffart et al., 2003; Guerrasio et al., 2010).

One path forward is to try and extend population coding of P-cells via their complex spike tuning into a method to infer population coding in the DCN. Because complex spikes tend to produce suppression of DCN activity, one may be able to use this error-dependent suppression to quantify tuning of a DCN neuron’s activity with respect to complex spikes in the parent P-cells. This error-dependent tuning may be a useful way to assign membership to DCN neurons that belong to the same population.

We focused on an elementary movement (saccades), but even simple gaze shifts involve coordination of motion between the eyes and the head. Visual errors that follow a head-free gaze shift produce adaptation of both saccades and head movements (Cecala and Freedman, 2009). In the framework presented here, these visual errors would engage P-cells that in turn influence DCN neurons that can produce an action that can remedy that error. Yet, stimulation of caudal fastigial nucleus produces eye saccades without significant motion of the head (Quinet and Goffart, 2009). At this writing it is unclear whether head-free and head-fixed gaze errors engage distinct populations of P-cells.

There is no denying that the vast majority of cerebellar output is destined to the cerebral cortex, where activity affects many aspects of behavior, including decision-making. For example, fastigial (Gao et al., 2018) or dentate nucleus (Chabrol et al., 2019) stimulation biases the delay period activity in the motor cortex as an animal evaluates the evidence associated with choosing a movement. Disruption of the interpositus nucleus eliminates working memory related activity in the prefrontal cortex (Siegel and Mauk, 2013). In humans, damage along the vermis has been associated with autism. Children who suffer from autism spectrum disorder, a developmental disorder that leads to impairments in social and communication skills, exhibit anatomic abnormalities in their cerebellum, including reduced number of P-cells (Whitney et al., 2008), particularly along the vermis (Marko et al., 2015; Scott et al., 2009).

How are P-cells contributing to social and communication skills? How are the activities in the DCN neurons supporting delay period activity of cortical cells during decision-making? These are some of the many unknown aspects of cerebellar function. It seems likely that some of the answers will be revealed once we better understand how the teacher (inferior olive) organizes the middle layer neurons (the P-cells) into populations that may serve as surrogate teachers for the actors (DCN neurons) that produce the output of the cerebellum.

## Acknowledgements

This work was supported by grants from the Office of Naval Research (N00014-15-1-2312), the National Institutes of Health (R01NS078311), and the National Science Foundation (CNS-1714623). I am grateful to Abigail Person for many discussions in which she pointed me to important papers and clarified ideas.

## References

Aizenman, C.D., Manis, P.B., Linden, D.J., 1998. Polarity of Long-Term Synaptic Gain Change Is Related to Postsynaptic Spike Firing at a Cerebellar Inhibitory Synapse. Neuron 21, 827–835. https://doi.org/10.1016/S0896-6273(00)80598-X

Albert, S.T., Shadmehr, R., 2018. Estimating properties of the fast and slow adaptive processes during sensorimotor adaptation. Journal of Neurophysiology 119, 1367–1393. https://doi.org/10.1152/jn.00197.2017

Andersen, R.A., Essick, G.K., Siegel, R.M., 1985. Encoding of spatial location by posterior parietal neurons. Science 230, 456–458.

Andersson, G., Armstrong, D.M., 1987. Complex spikes in Purkinje cells in the lateral vermis (b zone) of the cat cerebellum during locomotion. The Journal of Physiology 385, 107–134. https://doi.org/10.1113/jphysiol.1987.sp016487

Apps, R., Garwicz, M., 2005. Anatomical and physiological foundations of cerebellar information processing. Nat.Rev.Neurosci. 6, 297–311. https://doi.org/10.1038/nrn1646

Apps, R., Garwicz, M., 2000. Precise matching of olivo-cortical divergence and cortico-nuclear convergence between somatotopically corresponding areas in the medial C1 and medial C3 zones of the paravermal cerebellum. Eur.J.Neurosci. 12, 205–214. https://doi.org/10.1046/j.1460-9568.2000.00897.x

Barash, S., Melikyan, A., Sivakov, A., Zhang, M., Glickstein, M., Thier, P., 1999. Saccadic dysmetria and adaptation after lesions of the cerebellar cortex. J.Neurosci. 19, 10931–10939.

Bazzigaluppi, P., Ruigrok, T., Saisan, P., Zeeuw, C.I.D., Jeu, M. de, 2012. Properties of the Nucleo-Olivary Pathway: An In Vivo Whole-Cell Patch Clamp Study. PLOS ONE 7, e46360. https://doi.org/10.1371/journal.pone.0046360

Becker, M.I., Person, A.L., 2019. Cerebellar Control of Reach Kinematics for Endpoint Precision. Neuron 103, 335–348. https://doi.org/10.1016/j.neuron.2019.05.007

Bengtsson, F., Ekerot, C.-F., Jörntell, H., 2011. In Vivo Analysis of Inhibitory Synaptic Inputs and Rebounds in Deep Cerebellar Nuclear Neurons. PLoS One 6. https://doi.org/10.1371/journal.pone.0018822

Blenkinsop, T.A., Lang, E.J., 2011. Synaptic action of the olivocerebellar system on cerebellar nuclear spike activity. J.Neurosci. 31, 14708–14720. https://doi.org/10.1523/JNEUROSCI.3323-11.2011

Boele, H.-J., Koekkoek, S.K.E., Zeeuw, C.I.D., Ruigrok, T.J.H., 2013. Axonal Sprouting and Formation of Terminals in the Adult Cerebellum during Associative Motor Learning. J. Neurosci. 33, 17897–17907. https://doi.org/10.1523/JNEUROSCI.0511-13.2013

Bourrelly, C., Quinet, J., Goffart, L., 2018. The caudal fastigial nucleus and the steering of saccades toward a moving visual target. Journal of Neurophysiology 120, 421–438. https://doi.org/10.1152/jn.00141.2018

Brown, A.M., Arancillo, M., Lin, T., Catt, D.R., Zhou, J., Lackey, E.P., Stay, T.L., Zuo, Z., White, J.J., Sillitoe, R.V., 2019. Molecular layer interneurons shape the spike activity of cerebellar Purkinje cells. Sci.Rep. 9, 1742. https://doi.org/10.1038/s41598-018-38264-1

Burroughs, A., Wise, A.K., Xiao, J., Houghton, C., Tang, T., Suh, C.Y., Lang, E.J., Apps, R., Cerminara, N.L., 2017. The dynamic relationship between cerebellar Purkinje cell simple spikes and the spikelet number of complex spikes. J.Physiol 595, 283–299. https://doi.org/10.1113/JP272259

Buzunov, E., Mueller, A., Straube, A., Robinson, F.R., 2013. When during horizontal saccades in monkey does cerebellar output affect movement? Brain Res. 1503, 33–42. https://doi.org/10.1016/j.brainres.2013.02.001

Carta, I., Chen, C.H., Schott, A.L., Dorizan, S., Khodakhah, K., 2019. Cerebellar modulation of the reward circuitry and social behavior. Science 363. https://doi.org/10.1126/science.aav0581

Carulli, D., Broersen, R., Winter, F. de, Muir, E.M., Mešković, M., Waal, M. de, Vries, S. de, Boele, H.-J., Canto, C.B., Zeeuw, C.I.D., Verhaagen, J., 2020. Cerebellar plasticity and associative memories are controlled by perineuronal nets. PNAS 117, 6855–6865. https://doi.org/10.1073/pnas.1916163117

Catz, N., Dicke, P.W., Thier, P., 2008. Cerebellar-dependent motor learning is based on pruning a Purkinje cell population response. Proc.Natl.Acad.Sci.U.S.A 105, 7309–7314.

Catz, N., Dicke, P.W., Thier, P., 2005. Cerebellar complex spike firing is suitable to induce as well as to stabilize motor learning. Curr.Biol 15, 2179–2189.

Cecala, A.L., Freedman, E.G., 2009. Head-Unrestrained Gaze Adaptation in the Rhesus Macaque. Journal of Neurophysiology 101, 164–183. https://doi.org/10.1152/jn.90735.2008

Chabrol, F.P., Blot, A., Mrsic-Flogel, T.D., 2019. Cerebellar Contribution to Preparatory Activity in Motor Neocortex. Neuron 103, 506–519. https://doi.org/10.1016/j.neuron.2019.05.022

Chaumont, J., Guyon, N., Valera, A.M., Dugue, G.P., Popa, D., Marcaggi, P., Gautheron, V., Reibel-Foisset, S., Dieudonne, S., Stephan, A., Barrot, M., Cassel, J.C., Dupont, J.L., Doussau, F., Poulain, B., Selimi, F., Lena, C., Isope, P., 2013. Clusters of cerebellar Purkinje cells control their afferent climbing fiber discharge. Proc.Natl.Acad.Sci.U.S.A. 110, 16223–16228. https://doi.org/10.1073/pnas.1302310110

Chen, H., Hua, S.E., Smith, M.A., Lenz, F.A., Shadmehr, R., 2006. Effects of human cerebellar thalamus disruption on adaptive control of reaching. Cereb.Cortex 16, 1462–1473.

Churchland, M.M., Cunningham, J.P., Kaufman, M.T., Ryu, S.I., Shenoy, K.V., 2010. Cortical preparatory activity: representation of movement or first cog in a dynamical machine? Neuron 68, 387–400. https://doi.org/10.1016/j.neuron.2010.09.015

Coesmans, M., Weber, J.T., De Zeeuw, C.I., Hansel, C., 2004. Bidirectional Parallel Fiber Plasticity in the Cerebellum under Climbing Fiber Control. Neuron 44, 691–700. https://doi.org/10.1016/j.neuron.2004.10.031

Criscimagna-Hemminger, S.E., Shadmehr, R., 2008. Consolidation patterns of human motor memory. J.Neurosci. 28, 9610–9618.

Dash, S., Catz, N., Dicke, P.W., Thier, P., 2012. Encoding of smooth-pursuit eye movement initiation by a population of vermal Purkinje cells. Cereb.Cortex 22, 877–891. https://doi.org/10.1093/cercor/bhr153

Davie, J.T., Clark, B.A., Hausser, M., 2008. The origin of the complex spike in cerebellar Purkinje cells. J.Neurosci. 28, 7599–7609. https://doi.org/10.1523/JNEUROSCI.0559-08.2008

De Gruijl, J.R., Hoogland, T.M., Zeeuw, C.I.D., 2014. Behavioral Correlates of Complex Spike Synchrony in Cerebellar Microzones. J. Neurosci. 34, 8937–8947. https://doi.org/10.1523/JNEUROSCI.5064-13.2014

De Schutter, E., 1995. Cerebellar long-term depression might normalize excitation of Purkinje cells: a hypothesis. Trends in Neurosciences 18, 291–295. https://doi.org/10.1016/0166-2236(95)93916-L

De Zeeuw, C.I., Berrebi, A.S., 1995. Postsynaptic Targets of Purkinje Cell Terminals in the Cerebellar and Vestibular Nuclei of the Rat. European Journal of Neuroscience 7, 2322–2333. https://doi.org/10.1111/j.1460-9568.1995.tb00653.x

De Zeeuw, C.I., Hoebeek, F.E., Bosman, L.W., Schonewille, M., Witter, L., Koekkoek, S.K., 2011. Spatiotemporal firing patterns in the cerebellum. Nat.Rev.Neurosci. 12, 327–344. https://doi.org/10.1038/nrn3011

de Zeeuw, C.I., Holstege, J.C., Calkoen, F., Ruigrok, T.J.H., Voogd, J., 1988. A new combination of WGA-HRP anterograde tracing and GABA immunocytochemistry applied to afferents of the cat inferior olive at the ultrastructural level. Brain Research 447, 369–375. https://doi.org/10.1016/0006-8993(88)91142-0

De Zeeuw, C.I., Koekkoek, S.K., Wylie, D.R., Simpson, J.I., 1997. Association between dendritic lamellar bodies and complex spike synchrony in the olivocerebellar system. J.Neurophysiol. 77, 1747–1758. https://doi.org/10.1152/jn.1997.77.4.1747

De Zeeuw, C.I., Simpson, J.I., Hoogenraad, C.C., Galjart, N., Koekkoek, S.K., Ruigrok, T.J., 1998. Microcircuitry and function of the inferior olive. Trends Neurosci. 21, 391–400.

De Zeeuw, C.L., Van Alphen, A.M., Hawkins, R.K., Ruigrok, T.J., 1997. Climbing fibre collaterals contact neurons in the cerebellar nuclei that provide a GABAergic feedback to the inferior olive. Neuroscience 80, 981–986. https://doi.org/10.1016/s0306-4522(97)00249-2

Doya, K., 1999. What are the computations of the cerebellum, the basal ganglia and the cerebral cortex? Neural Networks. 12, 961–974.

Ebner, T.J., Bloedel, J.R., 1981. Correlation between activity of Purkinje cells and its modification by natural peripheral stimuli. J.Neurophysiol. 45, 948–961. https://doi.org/10.1152/jn.1981.45.5.948

Eccles, J.C., Llinás, R., Sasaki, K., 1966. The excitatory synaptic action of climbing fibres on the Purkinje cells of the cerebellum. The Journal of Physiology 182, 268–296. https://doi.org/10.1113/jphysiol.1966.sp007824

Ekerot, C.F., Jorntell, H., Garwicz, M., 1995. Functional relation between corticonuclear input and movements evoked on microstimulation in cerebellar nucleus interpositus anterior in the cat. Exp.Brain Res. 106, 365–376.

Fuchs, A.F., Robinson, F.R., Straube, A., 1993. Role of the caudal fastigial nucleus in saccade generation. I. Neuronal discharge pattern. J.Neurophysiol. 70, 1723–1740.

Gao, Z., Davis, C., Thomas, A.M., Economo, M.N., Abrego, A.M., Svoboda, K., De Zeeuw, C.I., Li, N., 2018. A cortico-cerebellar loop for motor planning. Nature 563, 113–116. https://doi.org/10.1038/s41586-018-0633-x

Gao, Z., van Beugen, B.J., De Zeeuw, C.I., 2012. Distributed synergistic plasticity and cerebellar learning. Nature Reviews Neuroscience 13, 619–635. https://doi.org/10.1038/nrn3312

Glaser, J.I., Wood, D.K., Lawlor, P.N., Ramkumar, P., Kording, K.P., Segraves, M.A., 2016. Role of expected reward in frontal eye field during natural scene search. J.Neurophysiol. 116, 645–657. https://doi.org/10.1152/jn.00119.2016

Goffart, L., Chen, L.L., Sparks, D.L., 2004. Deficits in saccades and fixation during muscimol inactivation of the caudal fastigial nucleus in the rhesus monkey. J.Neurophysiol. 92, 3351–3367.

Goffart, L., Chen, L.L., Sparks, D.L., 2003. Saccade dysmetria during functional perturbation of the caudal fastigial nucleus in the monkey. Ann.N.Y.Acad.Sci. 1004, 220–228.

Grasselli, G., Boele, H.-J., Titley, H.K., Bradford, N., Beers, L. van, Jay, L., Beekhof, G.C., Busch, S.E., Zeeuw, C.I.D., Schonewille, M., Hansel, C., 2020. SK2 channels in cerebellar Purkinje cells contribute to excitability modulation in motor-learning–specific memory traces. PLOS Biology 18, e3000596. https://doi.org/10.1371/journal.pbio.3000596

Grasselli, G., He, Q., Wan, V., Adelman, J.P., Ohtsuki, G., Hansel, C., 2016. Activity-Dependent Plasticity of Spike Pauses in Cerebellar Purkinje Cells. Cell Reports 14, 2546–2553. https://doi.org/10.1016/j.celrep.2016.02.054

Guerrasio, L., Quinet, J., Buttner, U., Goffart, L., 2010. Fastigial oculomotor region and the control of foveation during fixation. J.Neurophysiol. 103, 1988–2001. https://doi.org/10.1152/jn.00771.2009

Guo, C., Witter, L., Rudolph, S., Elliott, H.L., Ennis, K.A., Regehr, W.G., 2016. Purkinje Cells Directly Inhibit Granule Cells in Specialized Regions of the Cerebellar Cortex. Neuron 91, 1330–1341. https://doi.org/10.1016/j.neuron.2016.08.011

Han, K.S., Guo, C., Chen, C.H., Witter, L., Osorno, T., Regehr, W.G., 2018. Ephaptic Coupling Promotes Synchronous Firing of Cerebellar Purkinje Cells. Neuron 100, 564–578. https://doi.org/10.1016/j.neuron.2018.09.018

Hansel, C., de Jeu, M., Belmeguenai, A., Houtman, S.H., Buitendijk, G.H.S., Andreev, D., De Zeeuw, C.I., Elgersma, Y., 2006. αCaMKII Is Essential for Cerebellar LTD and Motor Learning. Neuron 51, 835–843. https://doi.org/10.1016/j.neuron.2006.08.013

Hausser, M., Clark, B.A., 1997. Tonic synaptic inhibition modulates neuronal output pattern and spatiotemporal synaptic integration. Neuron 19, 665–678. https://doi.org/10.1016/s0896-6273(00)80379-7

Heck, D.H., De Zeeuw, C.I., Jaeger, D., Khodakhah, K., Person, A.L., 2013. The neuronal code(s) of the cerebellum. J.Neurosci. 33, 17603–17609. https://doi.org/10.1523/JNEUROSCI.2759-13.2013

Heck, D.H., Thach, W.T., Keating, J.G., 2007. On-beam synchrony in the cerebellum as the mechanism for the timing and coordination of movement. Proc.Natl.Acad.Sci.U.S.A 104, 7658–7663. https://doi.org/10.1073/pnas.0609966104

Heffley, W., Hall, C., 2019. Classical conditioning drives learned reward prediction signals in climbing fibers across the lateral cerebellum. eLife 8:e46764.

Heffley, W., Song, E.Y., Xu, Z., Taylor, B.N., Hughes, M.A., McKinney, A., Joshua, M., Hull, C., 2018. Coordinated cerebellar climbing fiber activity signals learned sensorimotor predictions. Nat Neurosci 21, 1431–1441. https://doi.org/10.1038/s41593-018-0228-8

Helmchen, C., Buttner, U., 1995. Saccade-related Purkinje cell activity in the oculomotor vermis during spontaneous eye movements in light and darkness. Exp.Brain Res. 103, 198–208.

Helmchen, C., Straube, A., Buttner, U., 1994. Saccade-related activity in the fastigial oculomotor region of the macaque monkey during spontaneous eye movements in light and darkness. Exp.Brain Res. 98, 474–482.

Herzfeld, D.J., Hall, N.J., Tringides, M., Lisberger, S.G., 2020. Principles of operation of a cerebellar learning circuit. eLife 9, e55217. https://doi.org/10.7554/eLife.55217

Herzfeld, D.J., Kojima, Y., Soetedjo, R., Shadmehr, R., 2018. Encoding of error and learning to correct that error by the Purkinje cells of the cerebellum. Nat.Neurosci. 21, 736–743. https://doi.org/10.1038/s41593-018-0136-y

Herzfeld, D.J., Kojima, Y., Soetedjo, R., Shadmehr, R., 2015. Encoding of action by the Purkinje cells of the cerebellum. Nature 526, 439–442.

Hewitt, A.L., Popa, L.S., Ebner, T.J., 2015. Changes in Purkinje cell simple spike encoding of reach kinematics during adaption to a mechanical perturbation. J.Neurosci. 35, 1106–1124. https://doi.org/10.1523/JNEUROSCI.2579-14.2015

Hewitt, A.L., Popa, L.S., Pasalar, S., Hendrix, C.M., Ebner, T.J., 2011. Representation of limb kinematics in Purkinje cell simple spike discharge is conserved across multiple tasks. J.Neurophysiol. 106, 2232–2247. https://doi.org/10.1152/jn.00886.2010

Hoebeek, F.E., Witter, L., Ruigrok, T.J.H., Zeeuw, C.I.D., 2010. Differential olivo-cerebellar cortical control of rebound activity in the cerebellar nuclei. PNAS 107, 8410–8415. https://doi.org/10.1073/pnas.0907118107

Hong, S., Negrello, M., Junker, M., Smilgin, A., Thier, P., De, S.E., 2016. Multiplexed coding by cerebellar Purkinje neurons. eLife 5. https://doi.org/10.7554/eLife.13810

Horn, K.M., Pong, M., Gibson, A.R., 2004. Discharge of inferior olive cells during reaching errors and perturbations. Brain Research 996, 148–158. https://doi.org/10.1016/j.brainres.2003.10.021

Ikeda, T., Hikosaka, O., 2003. Reward-dependent gain and bias of visual responses in primate superior colliculus. Neuron 39, 693–700.

Ishikawa, T., Tomatsu, S., Tsunoda, Y., Lee, J., Hoffman, D.S., Kakei, S., 2014. Releasing dentate nucleus cells from Purkinje cell inhibition generates output from the cerebrocerebellum. PLoS.One. 9, e108774. https://doi.org/10.1371/journal.pone.0108774

Ito, M., Simpson, J.I., 1971. Discharges in Purkinje cell axons during climbing fiber activation. Brain Research 31, 215–219. https://doi.org/10.1016/0006-8993(71)90648-2

Jirenhed, D.A., Hesslow, G., 2016. Are Purkinje Cell Pauses Drivers of Classically Conditioned Blink Responses? Cerebellum. 15, 526–534. https://doi.org/10.1007/s12311-015-0722-4

Jörntell, H., Ekerot, C.-F., 2002. Reciprocal Bidirectional Plasticity of Parallel Fiber Receptive Fields in Cerebellar Purkinje Cells and Their Afferent Interneurons. Neuron 34, 797–806. https://doi.org/10.1016/S0896-6273(02)00713-4

Ju, C., Bosman, L.W.J., Hoogland, T.M., Velauthapillai, A., Murugesan, P., Warnaar, P., Genderen, R.M. van, Negrello, M., Zeeuw, C.I.D., 2019. Neurons of the inferior olive respond to broad classes of sensory input while subject to homeostatic control. The Journal of Physiology 597, 2483–2514. https://doi.org/10.1113/JP277413

Junker, M., Endres, D., Sun, Z.P., Dicke, P.W., Giese, M., Thier, P., 2018. Learning from the past: A reverberation of past errors in the cerebellar climbing fiber signal. PLOS Biology 16, e2004344. https://doi.org/10.1371/journal.pbio.2004344

Kaku, Y., Yoshida, K., Iwamoto, Y., 2009. Learning signals from the superior colliculus for adaptation of saccadic eye movements in the monkey. J.Neurosci. 29, 5266–5275. https://doi.org/10.1523/JNEUROSCI.0661-09.2009

Kase, M., Noda, H., Suzuki, D.A., Miller, D.C., 1979. Target velocity signals of visual tracking in vermal Purkinje cells of the monkey. Science 205, 717–720. https://doi.org/10.1126/science.111350

Ke, M.C., Guo, C.C., Raymond, J.L., 2009. Elimination of climbing fiber instructive signals during motor learning. Nat.Neurosci. 12, 1171–1179. https://doi.org/10.1038/nn.2366

Keating, J.G., Thach, W.T., 1995. Nonclock behavior of inferior olive neurons: interspike interval of Purkinje cell complex spike discharge in the awake behaving monkey is random. J.Neurophysiol. 73, 1329–1340. https://doi.org/10.1152/jn.1995.73.4.1329

Khaliq, Z.M., Raman, I.M., 2005. Axonal Propagation of Simple and Complex Spikes in Cerebellar Purkinje Neurons. J. Neurosci. 25, 454–463. https://doi.org/10.1523/JNEUROSCI.3045-04.2005

Khosrovani, S., Giessen, R.S.V.D., Zeeuw, C.I.D., Jeu, M.T.G.D., 2007. In vivo mouse inferior olive neurons exhibit heterogeneous subthreshold oscillations and spiking patterns. PNAS 104, 15911–15916. https://doi.org/10.1073/pnas.0702727104

Kim, J.J., Krupa, D.J., Thompson, R.F., 1998. Inhibitory cerebello-olivary projections and blocking effect in classical conditioning. Science 279, 570–573.

Kitazawa, S., Kimura, T., Yin, P.B., 1998. Cerebellar complex spikes encode both destinations and errors in arm movements. Nature 392, 494.

Kleine, J.F., Guan, Y., Buttner, U., 2003. Saccade-related neurons in the primate fastigial nucleus: what do they encode? J.Neurophysiol. 90, 3137–3154. https://doi.org/10.1152/jn.00021.2003

Kojima, Y., Iwamoto, Y., Yoshida, K., 2004. Memory of learning facilitates saccadic adaptation in the monkey. J.Neurosci. 24, 7531–7539.

Kojima, Y., Robinson, F.R., Soetedjo, R., 2014. Cerebellar fastigial nucleus influence on ipsilateral abducens activity during saccades. J.Neurophysiol. 111, 1553–1563. https://doi.org/10.1152/jn.00567.2013

Kojima, Y., Soetedjo, R., 2018. Elimination of the error signal in the superior colliculus impairs saccade motor learning. Proc.Natl.Acad.Sci.U.S.A 115, E8987–E8995. https://doi.org/10.1073/pnas.1806215115

Kojima, Y., Soetedjo, R., 2017. Change in sensitivity to visual error in superior colliculus during saccade adaptation. Sci.Rep. 7, 9566. https://doi.org/10.1038/s41598-017-10242-z

Kojima, Y., Soetedjo, R., Fuchs, A.F., 2010a. Effects of GABA agonist and antagonist injections into the oculomotor vermis on horizontal saccades. Brain Res. 1366, 93–100. https://doi.org/10.1016/j.brainres.2010.10.027

Kojima, Y., Soetedjo, R., Fuchs, A.F., 2010b. Changes in simple spike activity of some Purkinje cells in the oculomotor vermis during saccade adaptation are appropriate to participate in motor learning. J.Neurosci. 30, 3715–3727. https://doi.org/10/3715

Kording, K.P., Tenenbaum, J.B., Shadmehr, R., 2007. The dynamics of memory as a consequence of optimal adaptation to a changing body. Nat.Neurosci. 10, 779–786.

Kostadinov, D., Beau, M., Pozo, M.B., Hausser, M., 2019. Predictive and reactive reward signals conveyed by climbing fiber inputs to cerebellar Purkinje cells. Nat.Neurosci. 22, 950–962. https://doi.org/10.1038/s41593-019-0381-8

Kralj-Hans, I., Baizer, J.S., Swales, C., Glickstein, M., 2007. Independent roles for the dorsal paraflocculus and vermal lobule VII of the cerebellum in visuomotor coordination. Exp.Brain Res. 177, 209–222. https://doi.org/10.1007/s00221-006-0661-x

Lang, E.J., Apps, R., Bengtsson, F., Cerminara, N.L., De Zeeuw, C.I., Ebner, T.J., Heck, D.H., Jaeger, D., Jörntell, H., Kawato, M., Otis, T.S., Ozyildirim, O., Popa, L.S., Reeves, A.M.B., Schweighofer, N., Sugihara, I., Xiao, J., 2017. The Roles of the Olivocerebellar Pathway in Motor Learning and Motor Control. A Consensus Paper. Cerebellum 16, 230–252. https://doi.org/10.1007/s12311-016-0787-8

Larry, N., Yarkoni, M., Lixenberg, A., Joshua, M., 2019. Cerebellar climbing fibers encode expected reward size. eLife 8. https://doi.org/10.7554/eLife.46870

Lee, K.H., Mathews, P.J., Reeves, A.M.B., Choe, K.Y., Jami, S.A., Serrano, R.E., Otis, T.S., 2015. Circuit Mechanisms Underlying Motor Memory Formation in the Cerebellum. Neuron 86, 529–540. https://doi.org/10.1016/j.neuron.2015.03.010

Lee, K.H., Tran, A., Turan, Z., Meister, M., 2020. The sifting of visual information in the superior colliculus. eLife 9, e50678. https://doi.org/10.7554/eLife.50678

Lefler, Y., Yarom, Y., Uusisaari, M.Y., 2014. Cerebellar Inhibitory Input to the Inferior Olive Decreases Electrical Coupling and Blocks Subthreshold Oscillations. Neuron 81, 1389–1400. https://doi.org/10.1016/j.neuron.2014.02.032

Leigh, R.J., Zee, D.S., 2015. The neurology of eye movements. Oxford University Press.

Lu, H., Yang, B., Jaeger, D., 2016. Cerebellar Nuclei Neurons Show Only Small Excitatory Responses to Optogenetic Olivary Stimulation in Transgenic Mice: In Vivo and In Vitro Studies. Front Neural Circuits 10. https://doi.org/10.3389/fncir.2016.00021

Mano, N., Yamamoto, K., 1980. Simple-spike activity of cerebellar Purkinje cells related to visually guided wrist tracking movement in the monkey. J.Neurophysiol. 43, 713–728.

Marko, M.K., Crocetti, D., Hulst, T., Donchin, O., Shadmehr, R., Mostofsky, S.H., 2015. Behavioural and neural basis of anomalous motor learning in children with autism. Brain 138, 784–797. https://doi.org/10.1093/brain/awu394

Mascetti, G.G., Arriagada, J.R., 1981. Tectotectal interactions through the commissure of the superior colliculi: An electrophysiological study. Experimental Neurology 71, 122–133. https://doi.org/10.1016/0014-4886(81)90075-3

Mauk, M.D., Donegan, N.H., 1997. A model of Pavlovian eyelid conditioning based on the synaptic organization of the cerebellum. Learn.Mem. 4, 130–158.

May, P.J., Hartwich-Young, R., Nelson, J., Sparks, D.L., Porter, J.D., 1990. Cerebellotectal pathways in the macaque: implications for collicular generation of saccades. Neuroscience 36, 305–324.

Mays, L.E., Sparks, D.L., 1980. Dissociation of visual and saccade-related responses in superior colliculus neurons. J.Neurophysiol. 43, 207–232.

McElvain, L.E., Bagnall, M.W., Sakatos, A., du Lac, S., 2010. Bidirectional Plasticity Gated by Hyperpolarization Controls the Gain of Postsynaptic Firing Responses at Central Vestibular Nerve Synapses. Neuron 68, 763–775. https://doi.org/10.1016/j.neuron.2010.09.025

Medina, J.F., 2011. The multiple roles of Purkinje cells in sensori-motor calibration: to predict, teach and command. Current Opinion in Neurobiology, Sensory and motor systems 21, 616–622. https://doi.org/10.1016/j.conb.2011.05.025

Medina, J.F., Garcia, K.S., Mauk, M.D., 2001. A mechanism for savings in the cerebellum. J.Neurosci. 21.

Medina, J.F., Mauk, M.D., 1999. Simulations of cerebellar motor learning: computational analysis of plasticity at the mossy fiber to deep nucleus synapse. J.Neurosci. 19, 7140–7151.

Miles, F.A., Lisberger, S.G., 1981. Plasticity in the vestibulo-ocular reflex: a new hypothesis. Annu.Rev.Neurosci. 4, 273–299. https://doi.org/10.1146/annurev.ne.04.030181.001421

Monsivais, P., Clark, B.A., Roth, A., Häusser, M., 2005. Determinants of Action Potential Propagation in Cerebellar Purkinje Cell Axons. J. Neurosci. 25, 464–472. https://doi.org/10.1523/JNEUROSCI.3871-04.2005

Muller, S.Z., Zadina, A.N., Abbott, L.F., Sawtell, N.B., 2019. Continual Learning in a Multi-Layer Network of an Electric Fish. Cell. https://doi.org/10.1016/j.cell.2019.10.020

Munoz, D.P., Istvan, P.J., 1998. Lateral inhibitory interactions in the intermediate layers of the monkey superior colliculus. J.Neurophysiol. 79, 1193–1209.

Najac, M., Raman, I.M., 2017. Synaptic excitation by climbing fibre collaterals in the cerebellar nuclei of juvenile and adult mice. The Journal of Physiology 6703–6718. https://doi.org/10.1113/JP274598@10.1002/(ISSN)1469-7793(CAT)VirtualIssues(VI)EditorsChoice

Najafi, F., Giovannucci, A., Wang, S.S., Medina, J.F., 2014. Coding of stimulus strength via analog calcium signals in Purkinje cell dendrites of awake mice. eLife 3, e03663. https://doi.org/10.7554/eLife.03663

Nguyen-Vu, T.D., Kimpo, R.R., Rinaldi, J.M., Kohli, A., Zeng, H., Deisseroth, K., Raymond, J.L., 2013. Cerebellar Purkinje cell activity drives motor learning. Nat.Neurosci. 16, 1734–1736. https://doi.org/10.1038/nn.3576

Noda, H., Sugita, S., Ikeda, Y., 1990. Afferent and efferent connections of the oculomotor region of the fastigial nucleus in the macaque monkey. J.Comp Neurol. 302, 330–348. https://doi.org/10.1002/cne.903020211

O’Hearn, E., Molliver, M.E., 1997. The Olivocerebellar Projection Mediates Ibogaine-Induced Degeneration of Purkinje Cells: A Model of Indirect, Trans-Synaptic Excitotoxicity. J. Neurosci. 17, 8828–8841. https://doi.org/10.1523/JNEUROSCI.17-22-08828.1997

Ohmae, S., Medina, J.F., 2015. Climbing fibers encode a temporal-difference prediction error during cerebellar learning in mice. Nat.Neurosci. 18, 1798–1803. https://doi.org/10.1038/nn.4167

Ohyama, T., Mauk, M.D., 2001. Latent acquisition of timed responses in cerebellar cortex. J.Neurosci. 21, 682–690.

Ohyama, T., Nores, W.L., Mauk, M.D., 2003. Stimulus generalization of conditioned eyelid responses produced without cerebellar cortex: implications for plasticity in the cerebellar nuclei. Learn.Mem. 10, 346–354.

Ojakangas, C.L., Ebner, T.J., 1992. Purkinje cell complex and simple spike changes during a voluntary arm movement learning task in the monkey. J.Neurophysiol. 68, 2222–2236.

Payne, H.L., French, R.L., Guo, C.C., Nguyen-Vu, T.B., Manninen, T., Raymond, J.L., 2019. Cerebellar Purkinje cells control eye movements with a rapid rate code that is invariant to spike irregularity. eLife 8. https://doi.org/10.7554/eLife.37102

Pekny, S.E., Criscimagna-Hemminger, S.E., Shadmehr, R., 2011. Protection and expression of human motor memories. J.Neurosci. 31, 13829–13839. https://doi.org/10.1523/JNEUROSCI.1704-11.2011

Person, A.L., Raman, I.M., 2012a. Purkinje neuron synchrony elicits time-locked spiking in the cerebellar nuclei. Nature 481, 502–505. https://doi.org/10.1038/nature10732

Person, A.L., Raman, I.M., 2012b. Synchrony and neural coding in cerebellar circuits. Front Neural.Circuits. 6, 97. https://doi.org/10.3389/fncir.2012.00097

Person, A.L., Raman, I.M., 2010. Deactivation of L-type Ca Current by Inhibition Controls LTP at Excitatory Synapses in the Cerebellar Nuclei. Neuron 66, 550–559. https://doi.org/10.1016/j.neuron.2010.04.024

Piochon, C., Levenes, C., Ohtsuki, G., Hansel, C., 2010. Purkinje Cell NMDA Receptors Assume a Key Role in Synaptic Gain Control in the Mature Cerebellum. J. Neurosci. 30, 15330–15335. https://doi.org/10.1523/JNEUROSCI.4344-10.2010

Popa, L.S., Hewitt, A.L., Ebner, T.J., 2012. Predictive and Feedback Performance Errors Are Signaled in the Simple Spike Discharge of Individual Purkinje Cells. J. Neurosci. 32, 15345–15358. https://doi.org/10.1523/JNEUROSCI.2151-12.2012

Popa, L.S., Streng, M.L., Ebner, T.J., 2017. Long-Term Predictive and Feedback Encoding of Motor Signals in the Simple Spike Discharge of Purkinje Cells. eNeuro 4. https://doi.org/10.1523/ENEURO.0036-17.2017

Pugh, J.R., Raman, I.M., 2008. Mechanisms of Potentiation of Mossy Fiber EPSCs in the Cerebellar Nuclei by Coincident Synaptic Excitation and Inhibition. J. Neurosci. 28, 10549–10560. https://doi.org/10.1523/JNEUROSCI.2061-08.2008

Quinet, J., Goffart, L., 2015. Cerebellar control of saccade dynamics: contribution of the fastigial oculomotor region. J.Neurophysiol. 113, 3323–3336. https://doi.org/10.1152/jn.01021.2014

Quinet, J., Goffart, L., 2009. Electrical microstimulation of the fastigial oculomotor region in the head-unrestrained monkey. J.Neurophysiol. 102, 320–336. https://doi.org/10.1152/jn.90716.2008

Rasmussen, A., Hesslow, G., 2014. Feedback control of learning by the cerebello-olivary pathway. Prog.Brain Res. 210, 103–119. https://doi.org/10.1016/B978-0-444-63356-9.00005-4

Rasmussen, A., Jirenhed, D.A., Zucca, R., Johansson, F., Svensson, P., Hesslow, G., 2013. Number of spikes in climbing fibers determines the direction of cerebellar learning. J.Neurosci. 33, 13436–13440. https://doi.org/10.1523/JNEUROSCI.1527-13.2013

Rasmussen, A., Zucca, R., Johansson, F., Jirenhed, D.A., Hesslow, G., 2015. Purkinje cell activity during classical conditioning with different conditional stimuli explains central tenet of Rescorla-Wagner model [corrected]. Proc.Natl.Acad.Sci.U.S.A 112, 14060–14065. https://doi.org/10.1073/pnas.1516986112

Raymond, J.L., Medina, J.F., 2018. Computational Principles of Supervised Learning in the Cerebellum. Annual Review of Neuroscience 41, 233–253. https://doi.org/10.1146/annurev-neuro-080317-061948

Robinson, F.R., Straube, A., Fuchs, A.F., 1993. Role of the caudal fastigial nucleus in saccade generation. II. Effects of muscimol inactivation. J.Neurophysiol. 70, 1741–1758.

Ruigrok, T.J.H., Voogd, J., 2000. Organization of projections from the inferior olive to the cerebellar nuclei in the rat. Journal of Comparative Neurology 426, 209–228. https://doi.org/10.1002/1096-9861(20001016)426:2<209::AID-CNE4>3.0.CO;2-0

Saint-Cyr, J.A., Courville, J., 1982. Descending projections to the inferior olive from the mesencephalon and superior colliculus in the cat. An autoradiographic study. Exp.Brain Res. 45, 333–348.

Sarwary, A.M.E., Wischnewski, M., Schutter, D.J.L.G., Selen, L.P.J., Medendorp, W.P., 2018. Corticospinal correlates of fast and slow adaptive processes in motor learning. Journal of Neurophysiology 120, 2011–2019. https://doi.org/10.1152/jn.00488.2018

Sato, M., Hikosaka, O., 2002. Role of primate substantia nigra pars reticulata in reward-oriented saccadic eye movement. J.Neurosci. 22, 2363–2373.

Schiller, D., Monfils, M.H., Raio, C.M., Johnson, D.C., LeDoux, J.E., Phelps, E.A., 2010. Preventing the return of fear in humans using reconsolidation update mechanisms. Nature 463, 49–53.

Scott, J.A., Schumann, C.M., Goodlin-Jones, B.L., Amaral, D.G., 2009. A comprehensive volumetric analysis of the cerebellum in children and adolescents with autism spectrum disorder. Autism Res. 2, 246–257. https://doi.org/10.1002/aur.97

Scudder, C.A., McGee, D.M., Balaban, C.D., 2000. Connections of monkey saccade-related fastigial nucleus neurons revealed by anatomical and physiological methods. Soc.Neurosci.Abs. 26.

Sedaghat-Nejad, E., Herzfeld, D.J., Hage, P., Karbasi, K., Palin, T., Wang, X., Shadmehr, R., 2019. Behavioral training of marmosets and electrophysiological recording from the cerebellum. J.Neurophysiol. 122, 1502–1517.

Shin, S.L., De, S.E., 2006. Dynamic synchronization of Purkinje cell simple spikes. J.Neurophysiol. 96, 3485–3491. https://doi.org/10.1152/jn.00570.2006

Shutoh, F., Ohki, M., Kitazawa, H., Itohara, S., Nagao, S., 2006. Memory trace of motor learning shifts transsynaptically from cerebellar cortex to nuclei for consolidation. Neuroscience 139, 767–777.

Siegel, J.J., Mauk, M.D., 2013. Persistent activity in prefrontal cortex during trace eyelid conditioning: dissociating responses that reflect cerebellar output from those that do not. J.Neurosci. 33, 15272–15284. https://doi.org/10.1523/JNEUROSCI.1238-13.2013

Simpson, J.I., Wylie, D.R., Zeeuw, C.I.D., 1996. On climbing fiber signals and their consequence(s). Behavioral and Brain Sciences 19, 384–398. https://doi.org/10.1017/S0140525X00081486

Slemmer, J.E., De Zeeuw, C.I., Weber, J.T., 2005. Don’t get too excited: mechanisms of glutamate-mediated Purkinje cell death, in: Progress in Brain Research, Creating Coordination in the Cerebellum. Elsevier, pp. 367–390. https://doi.org/10.1016/S0079-6123(04)48029-7

Smith, M.A., Ghazizadeh, A., Shadmehr, R., 2006. Interacting adaptive processes with different timescales underlie short-term motor learning. PLoS.Biol. 4, e179.

Soetedjo, R., Fuchs, A.F., 2006. Complex spike activity of purkinje cells in the oculomotor vermis during behavioral adaptation of monkey saccades. J Neurosci. 26, 7741–7755.

Soetedjo, R., Fuchs, A.F., Kojima, Y., 2009. Subthreshold activation of the superior colliculus drives saccade motor learning. J.Neurosci. 29, 15213–15222. https://doi.org/10.1523/JNEUROSCI.4296-09.2009

Soetedjo, R., Kojima, Y., Fuchs, A.F., 2008. Complex spike activity in the oculomotor vermis of the cerebellum: a vectorial error signal for saccade motor learning? J.Neurophysiol. 100, 1949–1966.

Stollhoff, N., Menzel, R., Eisenhardt, D., 2005. Spontaneous recovery from extinction depends on the reconsolidation of the acquisition memory in an appetitive learning paradigm in the honeybee (Apis mellifera). J.Neurosci. 25, 4485–4492.

Streng, M.L., Popa, L.S., Ebner, T.J., 2018. Modulation of sensory prediction error in Purkinje cells during visual feedback manipulations. Nature Communications 9, 1099. https://doi.org/10.1038/s41467-018-03541-0

Sugihara, I., 2011. Compartmentalization of the Deep Cerebellar Nuclei Based on Afferent Projections and Aldolase C Expression. Cerebellum 10, 449–463. https://doi.org/10.1007/s12311-010-0226-1

Sugihara, I., Fujita, H., Na, J., Quy, P.N., Li, B.-Y., Ikeda, D., 2009. Projection of reconstructed single purkinje cell axons in relation to the cortical and nuclear aldolase C compartments of the rat cerebellum. Journal of Comparative Neurology 512, 282–304. https://doi.org/10.1002/cne.21889

Sugihara, I., Shinoda, Y., 2007. Molecular, Topographic, and Functional Organization of the Cerebellar Nuclei: Analysis by Three-Dimensional Mapping of the Olivonuclear Projection and Aldolase C Labeling. J. Neurosci. 27, 9696–9710. https://doi.org/10.1523/JNEUROSCI.1579-07.2007

Sugihara, I., Wu, H., Shinoda, Y., 1999. Morphology of single olivocerebellar axons labeled with biotinylated dextran amine in the rat. J.Comp.Neurol. 414, 131–148. https://doi.org/10.1002/(SICI)1096-9861(19991115)414:2<131::AID-CNE1>3.0.CO;2-F

Sugihara, I., Wu, H.-S., Shinoda, Y., 2001. The Entire Trajectories of Single Olivocerebellar Axons in the Cerebellar Cortex and their Contribution to Cerebellar Compartmentalization. J. Neurosci. 21, 7715–7723. https://doi.org/10.1523/JNEUROSCI.21-19-07715.2001

Sun, Z., Junker, M., Dicke, P.W., Thier, P., 2016. Individual neurons in the caudal fastigial oculomotor region convey information on both macro- and microsaccades. Eur.J.Neurosci. 44, 2531–2542. https://doi.org/10.1111/ejn.13289

Suvrathan, A., Payne, H.L., Raymond, J.L., 2016. Timing Rules for Synaptic Plasticity Matched to Behavioral Function. Neuron 92, 959–967. https://doi.org/10.1016/j.neuron.2016.10.022

Szapiro, G., Barbour, B., 2007. Multiple climbing fibers signal to molecular layer interneurons exclusively via glutamate spillover. Nat.Neurosci. 10, 735–742. https://doi.org/10.1038/nn1907

Takahashi, M., Sugiuchi, Y., Izawa, Y., Shinoda, Y., 2005. Commissural excitation and inhibition by the superior colliculus in tectoreticular neurons projecting to omnipause neuron and inhibitory burst neuron regions. J.Neurophysiol. 94, 1707–1726. https://doi.org/10.1152/jn.00347.2005

Tang, T., Blenkinsop, T.A., Lang, E.J., 2019. Complex spike synchrony dependent modulation of rat deep cerebellar nuclear activity. eLife 8. https://doi.org/10.7554/eLife.40101

Tang, T., Suh, C.Y., Blenkinsop, T.A., Lang, E.J., 2016. Synchrony is Key: Complex Spike Inhibition of the Deep Cerebellar Nuclei. Cerebellum. 15, 10–13. https://doi.org/10.1007/s12311-015-0743-z

Telgkamp, P., Raman, I.M., 2002. Depression of inhibitory synaptic transmission between Purkinje cells and neurons of the cerebellar nuclei. J.Neurosci. 22, 8447–8457.

ten Brinke, M.M., Heiney, S.A., Wang, X., Proietti-Onori, M., Boele, H.-J., Bakermans, J., Medina, J.F., Gao, Z., De Zeeuw, C.I., 2017. Dynamic modulation of activity in cerebellar nuclei neurons during pavlovian eyeblink conditioning in mice. eLife 6, e28132. https://doi.org/10.7554/eLife.28132

Teune, T.M., Burg, J. van der, Zeeuw, C.I. de, Voogd, J., Ruigrok, T.J.H., 1998. Single Purkinje cell can innervate multiple classes of projection neurons in the cerebellar nuclei of the rat: A light microscopic and ultrastructural triple-tracer study in the rat. Journal of Comparative Neurology 392, 164–178. https://doi.org/10.1002/(SICI)1096-9861(19980309)392:2<164::AID-CNE2>3.0.CO;2-0

Thach, W.T., 1967. Somatosensory receptive fields of single units in cat cerebellar cortex. J.Neurophysiol. 30, 675–696. https://doi.org/10.1152/jn.1967.30.4.675

Thier, P., Dicke, P.W., Haas, R., Barash, S., 2000. Encoding of movement time by populations of cerebellar Purkinje cells. Nature 405, 72–76.

Tomatsu, S., Ishikawa, T., Tsunoda, Y., Lee, J., Hoffman, D.S., Kakei, S., 2016. Information processing in the hemisphere of the cerebellar cortex for control of wrist movement. J.Neurophysiol. 115, 255–270. https://doi.org/10.1152/jn.00530.2015

Tseng, Y.W., Diedrichsen, J., Krakauer, J.W., Shadmehr, R., Bastian, A.J., 2007. Sensory prediction errors drive cerebellum-dependent adaptation of reaching. J.Neurophysiol. 98, 54–62.

Van Der Giessen, R.S., Koekkoek, S.K., van Dorp, S., De Gruijl, J.R., Cupido, A., Khosrovani, S., Dortland, B., Wellershaus, K., Degen, J., Deuchars, J., Fuchs, E.C., Monyer, H., Willecke, K., De Jeu, M.T.G., De Zeeuw, C.I., 2008. Role of Olivary Electrical Coupling in Cerebellar Motor Learning. Neuron 58, 599–612. https://doi.org/10.1016/j.neuron.2008.03.016

van der Kooij, K., Brenner, E., van Beers, R.J., Smeets, J.B.J., 2015. Visuomotor Adaptation: How Forgetting Keeps Us Conservative. PLoS One 10. https://doi.org/10.1371/journal.pone.0117901

Vaswani, P.A., Shadmehr, R., 2013. Decay of motor memories in the absence of error. J.Neurosci. 33, 7700–7709. https://doi.org/10.1523/JNEUROSCI.0124-13.2013

Vaswani, P.A., Shmuelof, L., Haith, A.M., Delnicki, R.J., Huang, V.S., Mazzoni, P., Shadmehr, R., Krakauer, J.W., 2015. Persistent residual errors in motor adaptation tasks: reversion to baseline and exploratory escape. J.Neurosci. 35, 6969–6977. https://doi.org/10.1523/JNEUROSCI.2656-14.2015

Viaro, R., Bonazzi, L., Maggiolini, E., Franchi, G., 2017. Cerebellar Modulation of Cortically Evoked Complex Movements in Rats. Cereb.Cortex 27, 3525–3541. https://doi.org/10.1093/cercor/bhw167

Vilis, T., Hore, J., 1980. Central neural mechanisms contributing to cerebellar tremor produced by limb perturbations. J.Neurophysiol. 43, 279–291.

Wallman, J., Fuchs, A.F., 1998. Saccadic gain modification: visual error drives motor adaptation. J Neurophysiol. 80, 2405–2416.

Welsh, J.P., Lang, E.J., Suglhara, I., Llinas, R., 1995. Dynamic organization of motor control within the olivocerebellar system. Nature 374, 453–457. https://doi.org/10.1038/374453a0

Whitney, E.R., Kemper, T.L., Bauman, M.L., Rosene, D.L., Blatt, G.J., 2008. Cerebellar Purkinje cells are reduced in a subpopulation of autistic brains: a stereological experiment using calbindin-D28k. Cerebellum. 7, 406–416. https://doi.org/10.1007/s12311-008-0043-y

Wise, A.K., Cerminara, N.L., Marple-Horvat, D.E., Apps, R., 2010. Mechanisms of synchronous activity in cerebellar Purkinje cells. J.Physiol 588, 2373–2390. https://doi.org/10.1113/jphysiol.2010.189704

Witter, L., Rudolph, S., Pressler, R.T., Lahlaf, S.I., Regehr, W.G., 2016. Purkinje Cell Collaterals Enable Output Signals from the Cerebellar Cortex to Feed Back to Purkinje Cells and Interneurons. Neuron 91, 312–319. https://doi.org/10.1016/j.neuron.2016.05.037

Wulff, P., Schonewille, M., Renzi, M., Viltono, L., Sassoè-Pognetto, M., Badura, A., Gao, Z., Hoebeek, F.E., van Dorp, S., Wisden, W., Farrant, M., De Zeeuw, C.I., 2009. Synaptic inhibition of Purkinje cells mediates consolidation of vestibulo-cerebellar motor learning. Nature Neuroscience 12, 1042–1049. https://doi.org/10.1038/nn.2348

Xu-Wilson, M., Chen-Harris, H., Zee, D.S., Shadmehr, R., 2009. Cerebellar contributions to adaptive control of saccades in humans. J.Neurosci. 29, 12930–12939.

Yang, Y., Lisberger, S.G., 2014a. Role of Plasticity at Different Sites across the Time Course of Cerebellar Motor Learning. J. Neurosci. 34, 7077–7090. https://doi.org/10.1523/JNEUROSCI.0017-14.2014

Yang, Y., Lisberger, S.G., 2014b. Purkinje-cell plasticity and cerebellar motor learning are graded by complex-spike duration. Nature. 510, 529–532. https://doi.org/10.1038/nature13282

Zhang, W., Linden, D.J., 2006. Long-Term Depression at the Mossy Fiber–Deep Cerebellar Nucleus Synapse. J. Neurosci. 26, 6935–6944. https://doi.org/10.1523/JNEUROSCI.0784-06.2006

